# Mapping odorant sensitivities reveals a sparse but structured representation of olfactory chemical space by sensory input to the mouse olfactory bulb

**DOI:** 10.1101/2022.05.11.491539

**Authors:** Shawn D. Burton, Audrey Brown, Thomas P. Eiting, Isaac A. Youngstrom, Thomas C. Rust, Michael Schmuker, Matt Wachowiak

## Abstract

In olfactory systems, convergence of sensory neurons onto glomeruli generates a map of odorant receptor identity. How glomerular maps relate to sensory space remains unclear. We sought to better characterize this relationship in the mouse olfactory system by defining glomeruli in terms of the odorants to which they are most sensitive. Using high-throughput odorant delivery and ultrasensitive imaging of sensory inputs, we imaged responses to 185 odorants presented at concentrations determined to activate only one or a few glomeruli across the dorsal olfactory bulb. The resulting datasets defined the tuning properties of glomeruli - and, by inference, their cognate odorant receptors - in a low-concentration regime, and yielded consensus maps of glomerular sensitivity across a wide range of chemical space. Glomeruli were extremely narrowly tuned, with ~25% responding to only one odorant, and extremely sensitive, responding to their effective odorants at sub-picomolar to nanomolar concentrations. Such narrow tuning in this concentration regime allowed for reliable functional identification of many glomeruli based on a single diagnostic odorant. At the same time, the response spectra of glomeruli responding to multiple odorants was best predicted by straightforward odorant structural features, and glomeruli sensitive to distinct odorants with common structural features were spatially clustered. These results define an underlying structure to the primary representation of sensory space by the mouse olfactory system.

## INTRODUCTION

Across animals, the first central representation of olfactory stimuli arises from the convergence of olfactory sensory neurons (OSNs) that express the same odorant receptor (OR) onto glomeruli of the olfactory bulb (OB) or antennal lobe, generating a map of OR identity across glomeruli. Characterizing odorant responses at the level of OSN input to glomeruli thus enables probing the functional properties of ORs in the intact animal, as well as understanding of how the representation of olfactory information is structured across the OSN population prior to its processing by central circuits. In the fly olfactory system, comprehensive characterization of OR-defined OSNs mapped to their cognate glomeruli has been foundational for understanding how olfactory information is represented and transformed by successive stages of central processing (Fishilevich and Vosshall, 2005; Hallem and Carlson, 2006; Caron et al., 2013). Achieving a similar level of characterization has been difficult in the mammalian olfactory system: to date, only 3-5% of mammalian ORs have been functionally characterized and mapped to their cognate glomeruli in vivo (Peterlin et al., 2014; Shirasu et al., 2014; Saito et al., 2017).

An additional challenge in the study of olfaction is the complexity of olfactory stimulus space, which includes a large number of compounds (>>10^4^ - >>10^9^) (Mayhew et al., 2022)that are not easily organized along physical dimensions such as wavelength or frequency. A related confound is the strong dependence of OSN specificity on odorant concentration and a lack of consensus on meaningful concentration ranges at which to characterize odor coding strategies (Meister and Bonhoeffer, 2001; Wachowiak and Cohen, 2001). With few exceptions (Si et al., 2019) previous studies have characterized OSN or glomerular responses to odorants presented at one or a few concentrations, typically far above threshold for evoking neural activity (Wachowiak and Cohen, 2001; Nara et al., 2011; Ma et al., 2012; Chae et al., 2019; Pashkovski et al., 2020; Soelter et al., 2020). The consensus from these and other studies is that odorant identity is encoded by combinatorial patterns of OSN and glomerular activity; however, details of such a coding strategy, including the logic of OSN tuning properties, the nature of glomerular maps across the OB surface, and the dimensionality of odorant representations remain unclear.

A useful approach to characterizing sensory response properties and central representations in other sensory systems is to measure neural responses relative to the parts of sensory space to which they are most sensitive – for example, characteristic frequencies in the auditory system (Evans et al., 1965). This approach avoids confounds from arbitrarily-chosen stimulus intensities, facilitates comparison across levels and approaches, and can more clearly reveal organizational features such as topographic mapping across neural space and transformations by neural circuits (M H Goldstein et al., 1970; Stiebler et al., 1997; Kandler et al., 2009). In olfaction, high-affinity odorant-glomerulus or odorant-OR interactions have been identified for only a handful of glomeruli or ORs (Oka et al., 2006; Zhang et al., 2012; Zhang et al., 2013; Peterlin et al., 2014; Sato-Akuhara et al., 2016; Horio et al., 2019; Soelter et al., 2020).

Here, we sought to identify ‘primary’ odorants (i.e., the odorant or odorants to which a glomerulus is most sensitive) for glomeruli of the dorsal OB, using ultrasensitive mapping of OSN inputs to glomeruli in anesthetized mice and efficient screening of a large, chemically diverse panel of odorants. We imaged glomerular responses to 185 odorants in single preparations and, for each odorant, determined a concentration that evoked activation of a small number of glomeruli, thus defining primary odorants for the majority of glomeruli across the dorsal OB. This approach yielded several foundational datasets, including: 1) consensus maps of glomerular odorant representations in the low (sub-nanomolar) concentration range, 2) an atlas of glomerular sensitivities for the dorsal OB across many odorants, and 3) a set of approximately two-dozen individual glomeruli that are robustly identified across animals using their activation by a single diagnostic odorant-concentration combination. These datasets revealed that OSN inputs to OB glomeruli – and, by extension, their cognate ORs – are exquisitely sensitive and selective to their primary odorants, such that representations of olfactory sensory space in this concentration regime are sparse and high-dimensional. Further, co-tuning of glomeruli to their few high-sensitivity odorants, as well as spatial maps of odorant sensitivities, revealed an underlying structure to these sparse representations that reflected relatively straightforward physicochemical features of odorants. This sparse but structured organization identifies an accessible framework for further analyses of how sensory information is represented and processed in in the mammalian olfactory system.

## RESULTS

### Generating consensus maps of odorant sensitivity across dorsal OB glomeruli

To map OSN inputs to OB glomeruli with high sensitivity and consistency across animals, we used tetracycline transactivator-amplified expression of the Ca^2+^ reporter GCaMP6s in all mature OSNs (OMP-IRES-tTA; tetO-GCaMP6s mice; see Methods). Consistent with earlier reports using the OMP-IRES-tTA driver line (Ma et al., 2014; Inagaki et al., 2020), this expression strategy did not appear to affect targeting of OSNs to their cognate glomeruli (Zhu et al., 2021). Odorant-evoked GCaMP6s signals were imaged with widefield epifluorescence across the dorsal surface of both OBs simultaneously in anesthetized mice, using artificial inhalation to ensure consistent odorant sampling (Eiting and Wachowiak, 2018).

We used a flexible, high-throughput odorant delivery system (Burton et al., 2019) to present a chemically diverse panel of 185 odorants (plus blank and solvent controls) to each experimental preparation. The panel covered a wide range of odorant chemical space as defined by physicochemical descriptors taken from a list of compounds curated for use in flavors and fragrances (**Fig. 1A**) (Pashkovski et al., 2020) (see Methods), and included a diversity of chemical classes as defined by functional group and other structural features (**Table S1, Supplementary Document S1**). Rather than deliver odorants at a single arbitrarily- or empirically-chosen concentration, we used a rational search strategy that allowed adjusting the concentration of each odorant across a >1000-fold (sub-pico-to nanomolar) range in order to identify high-sensitivity odorant-glomerulus interactions and to, ideally, pair each glomerulus with its primary odorant. Guided by recent studies indicating mouse perceptual thresholds in the picomolar range for at least some odorants (Dewan et al., 2018; Williams and Dewan, 2020), we initially set estimated delivered odorant concentrations to ~1 pM (see Methods). Across a series of pilot experiments, concentrations were then systematically increased (or occasionally decreased) by tenfold steps up to ~1 nM to identify the lowest concentration for each odorant capable of reliably activating at least one glomerulus, with responses averaged across at least three trials (typically, four) per concentration and odorant. Further increases in concentration were not considered as these were unlikely to reveal high-sensitivity interactions. The resulting concentrations were then used to screen the odorant panel across four final mice (eight OBs), with additional variations in concentration tested for many odorants (62-77 per mouse) to achieve comparable activation patterns while accounting for inter-animal variability. Final analyzed concentrations were identical across animals for the majority of odorants (145/185), and none differed by more than ten-fold across the four mice.

**Figure 1.**
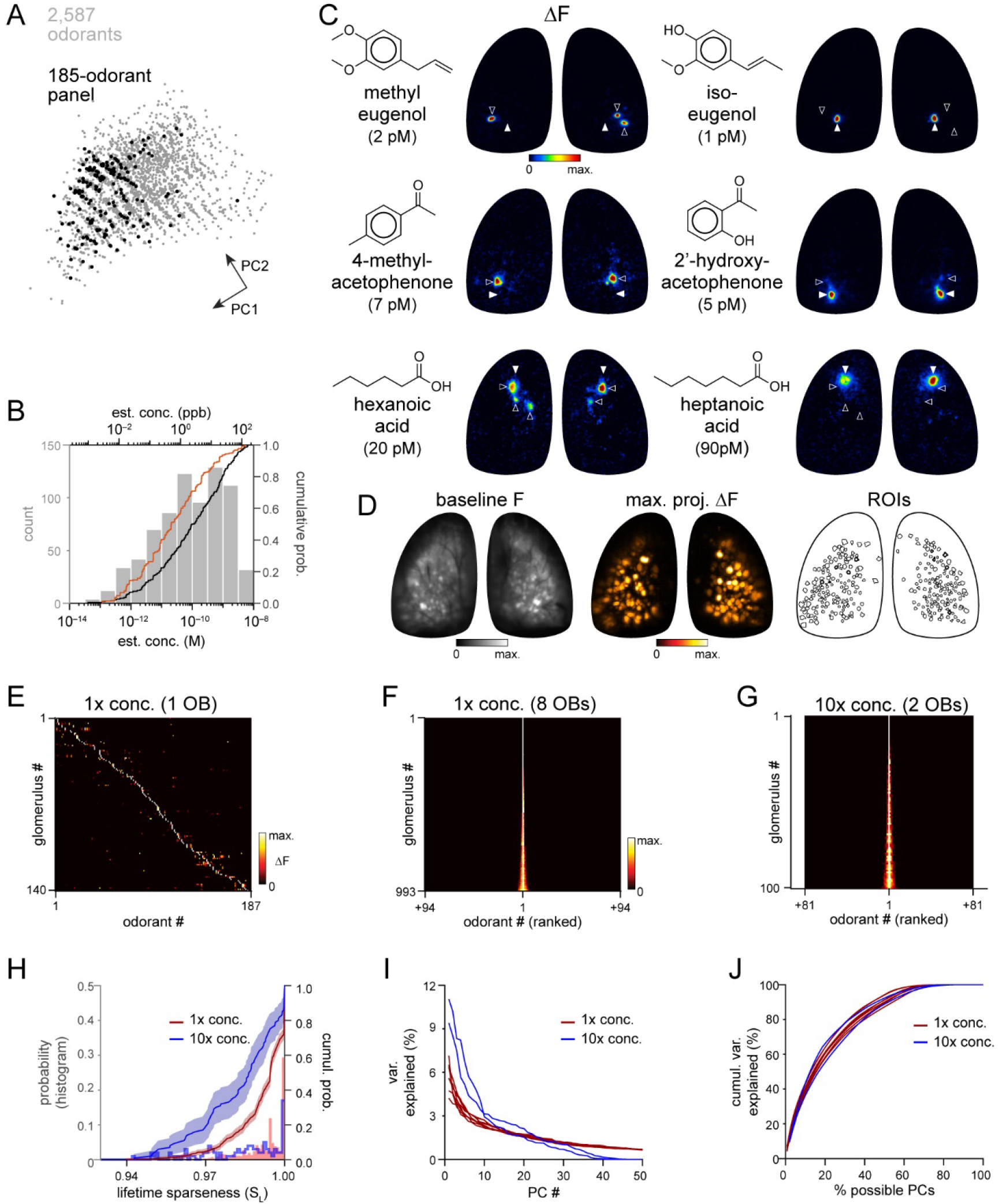
High sensitivity and narrow tuning of olfactory sensory input to OB glomeruli. **A**. Coverage of physicochemical space by the 185-odorant panel. Grey points, 2587 odorants plotted in the first two principal components of the physicochemical descriptor matrix, as in (Pashkovski et al., 2020) (see Methods). Black points indicate tested odorants. **B**. Estimated delivered concentrations used across the odorant panel. Histogram and black cumulative distribution function show concentrations of each presented odorant across four preparations (n = 740). Red plot shows cumulative distribution function of minimal effective concentration for each responsive glomerulus (n = 993). **C**. Response maps evoked by single odorants. Top three rows show distinct but neighboring glomeruli activated by structurally similar odorants. All images taken from the same preparation. Estimated concentrations are rounded to single-significant digit precision. **D**. Left: baseline fluorescence of the mouse shown in (C). Middle: maximal projection of responses across the 185-odorant panel. Right: ROIs of responsive glomeruli. **E**. Matrix of responses across all responsive glomeruli in one OB. Each row (glomerulus) is normalized to the maximal response across the odorant panel. Glomeruli are sorted in order of their maximally-activating odorant, producing a pseudo-diagonalized matrix. Odorants are ordered according to nominal structural classification (see Methods). Matrix includes responses to empty and solvent controls. **F, G**. Response spectra of all imaged glomeruli (rows) across the odorant panel (columns), normalized by maximal response, for 1x concentration epifluorescence dataset and the 10x concentration two-photon dataset (separate preparations; 10x two-photon data imaged from a smaller field of view containing fewer glomeruli). Odorant order sorted by response amplitude; glomerular order sorted by lifetime sparseness. **H**. Histogram and cumulative distribution plots of lifetime sparseness values for all responsive glomeruli for the odorant panel presented at original, 1x concentrations (red; n = 993 glomeruli) and at 10x concentrations (blue; n = 100). Shading denotes 95% confidence intervals (calculated using ‘ecdf’ function in Matlab). **I**. Response variance explained versus PC number for each imaged OB. Red plots: 1x concentrations, n = 8 OBs; blue plots: 10x concentrations, n = 2 OBs. **J**. Variance explained versus fraction of possible PCs for same data as in (I).

Nearly all odorants tested proved effective within the picomolar-to-nanomolar concentration range: 163 of 185 odorants (88%) evoked responses in one or more dorsal glomeruli per OB per mouse, and only twelve odorants proved ineffective (i.e., failed to elicit a response in any OB). Qualitatively, patterns of glomerular activation (defined by the location and number of activated glomeruli) were remarkably consistent across all eight OBs, with only modest differences in overall response magnitudes between left and right OBs in the same mouse. We used the resulting dataset as a resource for defining consensus high-sensitivity response maps for this large odorant panel across the dorsal OB (**Document S2**).

### Sensory inputs to glomeruli are highly sensitive and narrowly tuned to their primary odorants

Across all odorants, final estimated concentrations ranged from 4 × 10^−14^ to 4 × 10^−9^ M (median: 1 × 10^−10^ M, 2.4 ppb) (**Fig. 1B**; **Table S1**). The cumulative number of glomeruli activated across the odorant panel ranged from 103 to 142 per OB (median: 126) and covered the extent of the dorsal OB (**Fig. 1C, D; Fig. S1**), indicating that high odorant sensitivity is a general feature of OSN inputs to glomeruli. Indeed, glomerular sensitivity (i.e., the concentration at which each glomerulus responded to its primary odorant) was distributed across even lower concentrations, with a median of 2 × 10^−11^ M (0.5 ppb) (**Fig. 1B**). These concentrations are, overall, substantially lower than those used to characterize odorant representations in earlier studies, by as much as 4-5 orders of magnitude (**Fig. S2**). We next assessed how canonical features of olfactory stimulus coding at the level of OSN input to glomeruli manifest in this concentration regime, beginning with the tuning of individual glomeruli and the nature of glomerular representations of individual odorants.

Our experimental approach was designed to yield sparse glomerular responses to each odorant. Nevertheless, the degree of sparseness was striking, with each effective odorant evoking input to only a few glomeruli (median: 2 glomeruli per odorant; quartiles: 1-3 glomeruli per odorant; 1290 responses across 8 OBs) (**Fig. S3A**). Population sparseness (S_P_), a measure of the selectivity of glomerular activation for a given odorant, was exceptionally high (median S_P_: 0.993; quartiles: 0.989-0.997, calculated from mean S_P_ per odorant across 8 OBs) (**Fig. S3B**). This result implies that individual glomeruli are narrowly tuned across the entire odorant panel. Indeed, highly selective tuning was evident in individual response maps, with individual glomeruli often responding strongly to a given odorant and not at all to structurally similar odorants (**Fig. 1C**).

Earlier characterizations of OR tuning have identified a mix of narrowly- and broadly-tuned ORs (Hallem and Carlson, 2006; Saito et al., 2009; Nara et al., 2011). Here, we observed narrow tuning of OSN inputs to nearly all glomeruli, and for glomeruli tuned to nearly all odorants (**Fig. 1E, F; Fig. S1**). Lifetime sparseness (S_L_), a measure of stimulus selectivity that reflects the distribution of response magnitudes across a stimulus set and ranges from 0 (no stimulus selectivity) to 1 (response to only a single stimulus) (Davison and Katz, 2007; Schlief and Wilson, 2007), was extremely high across the glomerular population, with a median value of 0.995 (mean of medians across 8 OBs; mean quartiles: 0.988-1) (**Fig. 1H**). 25-30% of glomeruli in a given OB (mean ± s.d.: 29 ± 6%, 288/1004 total glomeruli) responded to only one odorant from the panel, and only 19% of glomeruli (187/1004) responded to more than 5 odorants (**Fig. S3C**).

Sparse responses and narrow tuning of glomeruli was not due to an ‘iceberg effect’, in which only a few odorants evoked responses detectable above the signal-to-noise ratio of our imaging approach, as response amplitudes were typically many times greater than baseline variance levels. 75% of glomeruli that responded to more than one odorant (n = 716) had a dynamic range (i.e., the ratio between the strongest and weakest odorant responses) greater than 2, 50% had a range greater than 4, and 20% had a range greater than 10, indicating a high dynamic range in odorant responsiveness even in this low-concentration regime. S_L_ values were also consistently high and independent of both dynamic range and maximum response amplitude (**Fig. S3D, E**), indicating that sparse responses are not an artifact of low signal-to-noise ratios or of weakly-responding glomeruli.

We further tested the concentration-dependence of OSN tuning by presenting 159 odorants of the panel at tenfold higher concentrations in two separate preparations (the remaining 26 odorants were presented at the same concentration as the widefield imaging set but omitted from this concentration-dependent analysis; see Table S1), using two-photon imaging from a central subregion of the dorsal OB in order to more precisely attribute signals to specific glomeruli (**Fig. S4**). Glomerular tuning was only slightly broader at these higher concentrations, with 12% of glomeruli activated by only one of the 159 odorants and a median S_L_ of 0.986 (quartiles: 0.971-0.995; 100 responsive glomeruli, 2 mice) (**Fig. 1G, H**). Narrow OSN tuning was thus not restricted to carefully-chosen perithreshold concentrations, suggesting that sparse and selective glomerular activation is a robust feature of odorant representations in low-concentration regimes.

Previous studies examining neural responses to higher concentrations have proposed that odorant identity can be reliably encoded within a structured, lower-dimensional neural coding space (Chae et al., 2019; Si et al., 2019; Pashkovski et al., 2020). However, glomerulus-odorant response matrices in our low-concentration datasets appeared high-dimensional, at least in the Euclidean domain. The first principal component (PC) in each OB dataset accounted for only 4-7% of the total variance (**Fig. 1I**), and 54-67 PCs (44-55% of possible PCs, given 100-140 glomeruli per OB) were required to account for 90% of the response variance (**Fig. 1J**), with similar results observed for the tenfold higher concentration two-photon dataset. These results substantially differ from the 21 PCs required to account for 90% of the response variance in a pseudopopulation of 871 glomeruli (i.e., ~2.4% of possible PCs) in a comparable analysis of glomerular responses to substantially higher odorant concentrations (Chae et al., 2019). Effective dimensionality (ED), a measure of response covariance across a dataset (Litwin-Kumar et al., 2017), was also substantially higher in our main dataset (mean ± s.d.: 48.9±4.4, n=8 OBs) than that reported for odorant responses imaged from OB projections to piriform cortex using higher odorant concentrations (ED: 13.5, n=22 odorants × 3160 axonal boutons) (Pashkovski et al., 2020).

### Diagnostic odorants enable widespread functional identification of glomeruli

The narrow tuning and high sensitivity of glomeruli to their primary odorants suggests that many glomeruli can be identified simply by their response to a single diagnostic odorant delivered at low concentration. Examples of five such diagnostic odorants evoking consistent singular- or near-singular activation of glomeruli are shown in **Fig. 2A**. These glomeruli appear in a consistent location in each OB and exhibit near-identical response spectra across the full odorant panel (**Fig. 2B**). For example, the aromatic ester phenylacetate (~3 pM) strongly activated a single glomerulus in the central OB, with identical response spectra across the 8 OBs (**Fig. 2A, B**). Likewise, the odorant methyl tiglate (~10 pM) strongly activated a single glomerulus in the central-medial OB in each of the 8 OBs imaged; this glomerulus was also strongly- and singly-activated by ethyl, hexyl, and isopropyl tiglate, as well as trans-2-methyl-2-butenal and 2-methyl 2-pentenoic acid (**Fig. 2A, B**). These observations suggest that, in many cases, a single diagnostic odorant can be used to identify putatively cognate glomeruli across animals.

**Figure 2.**
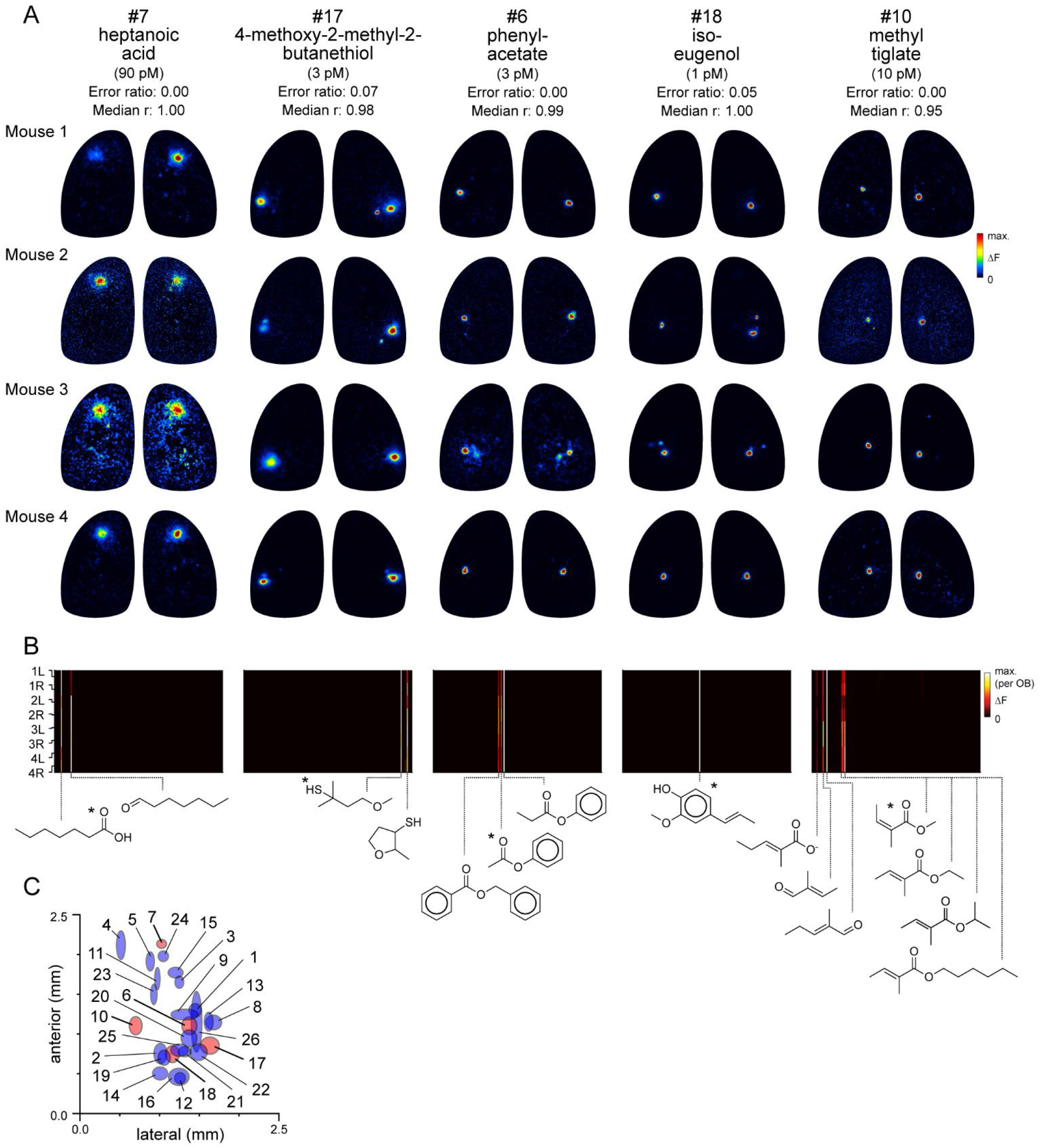
Functional identification of glomeruli using singular activation by diagnostic odorants. **A**. Response maps evoked by five odorants, shown for each of four mice, eliciting singular or near-singular activation of a glomerulus in a consistent location in each OB. See Text for definition of error ratio and median r. **B**. Response spectra for each glomerulus in (A) across the 185-odorant panel, shown for each of the 8 imaged OBs. Pseudocolor scale is normalized to the maximal response in the glomerulus for each preparation. Structures of effective odorants are shown at bottom. Asterisk indicates diagnostic odorant shown in (A). **C**. Mean locations of all functionally identified glomeruli, referenced to the midline and caudal sinus of the OB. Spot width and height indicate jitter (s.d.) of medial-lateral and rostro-caudal location across the 8 OBs. Identified glomeruli from (A) are shown in red; all other glomeruli shown in blue.

To identify additional such glomeruli and their diagnostic odorants from the dataset, we first screened odorants using highly conservative criteria for sparseness and response reliability (see Methods). For each of the 80 odorants passing this screen, we tested the odorant’s ability to identify glomeruli across OBs by comparing the response spectrum of the strongest-activated glomerulus by that odorant across the full 185-odorant panel with that of all other glomeruli in each OB. For 19 odorants, the strongest (or only) activated glomerulus identified the glomerulus with the highest-correlated response spectrum in 100% of comparisons (median correlation coefficient [Pearson’s r] across OBs: 0.95 ± 0.04; mean ± s.d., n = 19). Relaxing these criteria slightly to allow for inherent variability in responsiveness across the odorant panel – to a cutoff of 80% match between the strongest-activated and most-correlated glomerulus across OBs (error ratio <0.2; see Methods) and a median correlation coefficient in response spectra of >0.8 – yielded 22 additional odorants (41 odorants total) that identified 26 unique glomeruli (**Document S3**; **Table S2**). **Table S3** lists the remaining odorants that elicited reliably sparse glomerular activity but which did not meet our conservative criteria for functional identification. Notably, while spatial location was not used as a diagnostic criterion, glomeruli identified with this approach appeared in a similar location on the dorsal OB, with a spatial jitter consistent with that characterized for OR-defined OSN projections (**Fig. 2C; Table S2**) (Zapiec and Mombaerts, 2015). Diagnostic odorant-concentration pairs also evoked singular activation of the same putatively identified glomeruli in awake, head-fixed mice (**Fig. S5**). Thus, the high selectivity and sensitivity of OSNs for their primary odorants allows for simple and robust identification of cognate glomeruli across OBs using a single odorant-concentration pair.

We then further explored the logic of OSN tuning by analyzing the response spectra of the identified glomeruli, compiled across the 8 OBs (**Fig. 3A**). Consensus response spectra were defined by the median response to each odorant across the 8 OBs. The median S_L_ across the 26 glomeruli was 0.99 (mean ± SD : 0.988±0.011), indicating that the functionally-identified glomeruli exhibited similarly narrow tuning as the general glomerular population. Seven of the 26 glomeruli responded to only a single odorant. Of the remaining 19 glomeruli, effective co-tuned odorants often shared common structural features such as functional group, carbon chain length (for aliphatic odorants), ring structure, or heteroatom. At the same time, nearly all glomeruli were exquisitely selective to their co-tuned odorants, often showing a strong response to one and no response to other structurally similar odorants. Several of the identified glomeruli appeared relatively broadly tuned to heterocyclic compounds including pyrazines and thiazoles (e.g., glomeruli 13, 14, and 16, **Document S3**). In addition, several glomeruli showed high sensitivity to odorants where chemical similarity was less obvious; for example, glomeruli 15, 19, and 20 were sensitive to both aromatic and aliphatic compounds, and glomerulus 14 was highly sensitive to the cyclic terpenoid (R)-(+)-pulegone as well as several pyrazines.

**Figure 3.**
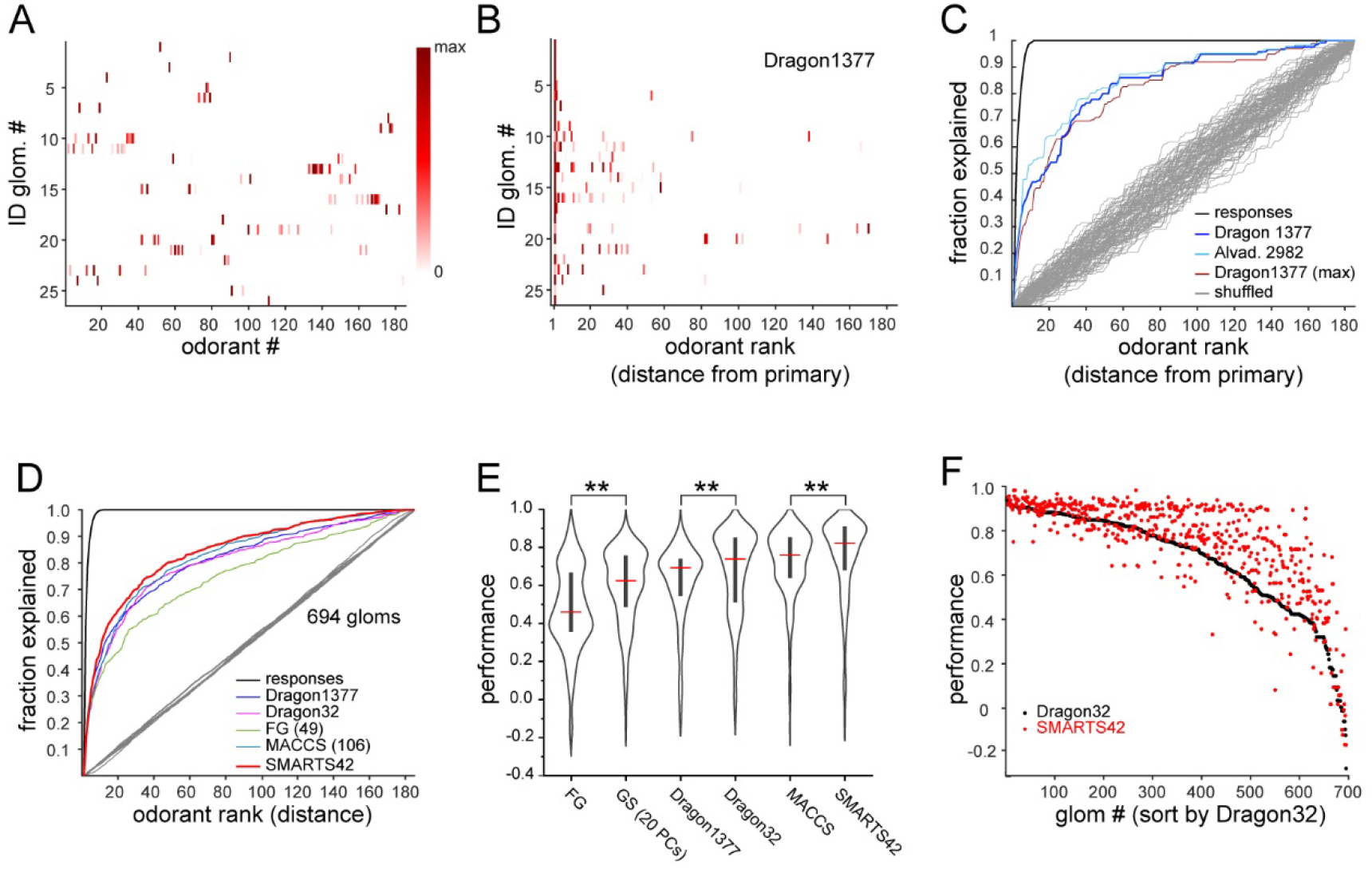
Predicting odorant response specificity from physicochemical feature sets. **A**. Median response spectra of all functionally-identified glomeruli. Each spectrum (row) is normalized to its maximal odorant response. Odorants ordered as in Fig. 1G. **B**. Median response spectra of functionally-identified glomeruli, with odorants ordered by physicochemical descriptor distance from the primary odorant for each glomerulus (odorant that evokes a response at the lowest concentration), using the Dragon1377 descriptor set. **C**. Mean receiver-operating characteristic (ROC) curves across all 19 co-tuned identified glomeruli, showing fraction of all effective odorant responses explained as a function of ranked odorant distance from the primary odorant. Colored lines show performance curves for the Dragon1377 descriptor set, a larger 2982-element descriptor set (Alvad. 2982), and the Dragon1377 set distances relative to the strongest-activating odorant (Dragon1377 (max)), as described in the Text. Black plot shows cumulative response function of binarized odorant responses, with odorants ranked by response magnitude, representing perfect response prediction. Grey plots show performance using random ranking of odorants (100 iterations). **D**. Mean ROC curves for all non-singularly activated glomeruli (n = 694) across the 8 OBs, relative to the primary odorant for each glomerulus, for different physicochemical descriptor sets, as defined in Text. Grey plots show mean performance across random rankings. **E**. Distribution of performance scores across all 694 glomeruli for different descriptor sets. Red bar, median; Center bar, interquartile range; envelope, smoothed point density. Asterisks indicate significant difference between sets (Kruskal-Wallis test comparing all descriptor set performance metrics: p<2 × 10^−16^; chi-squared statistic, 859; df = 6; p=0.002 for MACCS vs. Dragon32; p<1 × 10^−25^ for MACCS versus all other descriptor sets (post-hoc Dunn tests, Benjamani-Hochberg correction for multiple comparisons); only differences between incrementally higher medians are shown for clarity). **F**. Prediction performance scores for each of the 694 non-singularly activated glomeruli for the 32-element optimized Dragon descriptor subset (Dragon32; black points) and the 42-element SMARTS set (SMARTS42; red points), with glomeruli ordered by performance in the Dragon32 descriptor space.

To further test the concentration-dependence of glomerular tuning, we located six of the functionally-identified glomeruli using epifluorescence widefield imaging at their diagnostic (‘1x’) odorant concentrations, and then characterized their response spectra to the tenfold-higher concentration odorant panel (‘10x’) using two-photon imaging (n=1-2 OBs), as described above (**Fig. S4**). Consistent with the summary analysis across all glomeruli, the response spectrum of each glomerulus at these 10x concentrations was nearly identical to the median response spectrum at the 1x concentrations, with smaller-magnitude responses occasionally recruited in response to some additional odorants (**Fig. S6**). This result further supports the conclusion that narrow tuning is a robust feature of OSN inputs to OB glomeruli.

### Basic structural features predict co-tuning of glomeruli to their high-sensitivity odorants

Earlier studies have used sets of physicochemical descriptors of odorants to infer relationships between the chemical space of odorants and their neural representations, OR tuning, or odor perception (Haddad et al., 2008; Saito et al., 2009; Chae et al., 2019; Soelter et al., 2020; Gerkin, 2021). Given the exceptionally narrow tuning of glomerular inputs in our dataset, we used this approach to explore the chemical relationships between the small number of odorants to which OSN populations were most sensitive. We first focused on the 19 functionally-identified glomeruli that were responsive to more than one odorant, sorting the median response spectra of each glomerulus according to odorant distance in physicochemical descriptor space, relative to the primary odorant for that glomerulus. We initially used a subset of 1377 descriptors from the E-Dragon web app (Todeschini and Consonni, 2003; Tetko et al., 2005) chosen previously from a large-scale characterization of mammalian OR binding properties (Haddad et al., 2008; Saito et al., 2009; Chae et al., 2019; Soelter et al., 2020; Gerkin, 2021). This descriptor set appeared moderately effective at predicting glomerulus co-tuning given the primary odorant identity, although some glomeruli were co-tuned to odorants distributed across large distances in the descriptor space, and all glomeruli failed to respond to numerous odorants located more closely within this space (**Fig. 3B**).

To quantitatively assess the ability of these physicochemical descriptors to predict glomerulus co-tuning, we derived a performance metric from receiver-operating characteristic analysis (see Methods) based on the cumulative fraction of responses explained by each successively ranked odorant, relative to shuffled odorant ordering (**Fig. 3C**). The performance metric had a value of 1 for perfect prediction of odorant responses and 0 for prediction no different from chance. Performance metrics for functionally-identified glomeruli were well above chance levels (median: 0.73; quartiles: 0.60-0.96). Similar results were obtained with models generated from expanded physicochemical descriptor databases (2982 descriptors, Alvadesc) (Pashkovski et al., 2020) (median: 0.75; quartiles: 0.63-0.93), and when querying response spectra relative to the strongest-activating (rather than most-sensitive, or primary) odorant (median: 0.82; quartiles: 0.66-0.96) (**Fig. 3C, Fig. S7A**).

We next extended this analysis to the full dataset of glomerular responses. Performance metrics were calculated for all glomeruli responding to more than one odorant at a level greater than 10% of its maximal response (n = 694 glomeruli). To avoid assuming a priori knowledge of the primary odorant for a glomerulus, for this analysis we measured performance using each effective odorant as the query odorant, using the median of these values as the performance metric for each glomerulus (see Methods). Finally, we compared the quality of different models of odorant chemical space at predicting glomerulus co-tuning, using a number of additional physicochemical descriptor sets. Specifically, we used an optimized subset of 32 descriptors selected from fitting to earlier published odorant response datasets (Haddad et al., 2008), a subset (49) of the 154 descriptors defining odorant functional groups (Saito et al., 2009), and a subset (108) of the 166 MACCS chemical feature keyset bits (Durant et al., 2002) (see Methods). We also defined odorant distances using the first 20 principal components of the chemical space defined by physicochemical descriptors for the curated set of 2624 odorants, as in Fig. 1A (**Fig. S7B, D**). Of these, the MACCS feature keys had significantly higher performance metrics than the other descriptor sets (**Fig. 3D, E**).

It was notable that the set of 108 MACCS keys outperformed exhaustive physicochemical descriptor sets and performed equivalently to the 32-descriptor set that was optimized by fitting to neural response data (Haddad et al., 2008). Inspired by this result, we defined a new set of structural features using the SMARTS chemical pattern matching language (Daylight Chemical Information Systems Inc.). This set shared some features with the 49-element functional group descriptors (‘FG’), with an additional emphasis on resolving larger substructural motifs that are common in odorant compounds, including the presence and length of an aliphatic carbon chain; heteroatom substitutions, ortho-, meta- and para-substituted rings, and a number of terpenoid scaffolds (**Table S4**). The resulting ‘SMARTS42’ set was small (42 features), but notably had a higher predictive quality than all other models, with higher performance metrics across the entire glomerular dataset (**Fig. 3E, F**), as well as for the identified glomeruli (**Fig. S7C, E, F**) (SMARTS42 vs. MACCS, p=4.0 × 10^−7^, corrected post-hoc Dunn test). These results suggest that, despite the extremely narrow tuning of glomeruli overall, the response spectrum of a given glomerulus in low-concentration regimes is moderately well-predicted from relatively straightforward structural features.

Given this result we hypothesized that, despite the overall high dimensionality of odorant representations arising from narrowly-tuned glomeruli, any correlational structure in odorant representations is likely to reflect such basic structural features. Indeed, odorant response correlation matrices revealed blocks of correlated responses that corresponded to major structural classes of odorants. In particular, the highest correlations occurred among aliphatic acids, aldehydes and esters; primary amines; and pyrazines and thiazoles/pyrroles/pyridines (‘heterocyclic N-S’) (**Fig. 4A**). As each odorant activated only a few glomeruli, correlation analysis was poorly suited to further investigating the structure of odorant response matrices. Instead, since correlation coefficients essentially reflected co-tuning of individual glomeruli to a few odorants, we constructed pairwise odorant co-tuning matrices directly based on the number of glomeruli in a given OB that were co-activated by each odorant pair, regardless of response magnitude. **Fig. 4B** shows the mean pairwise odorant co-tuning matrix across the 8 OBs, thresholded at 0.875 co-tuned glomeruli per OB (i.e., co-tuning in at least 7 of 8 OBs), to identify the most reliable co-tuning relationships among glomeruli. The co-tuning matrix revealed a similar structure to that of the correlation matrix, with blocks of co-tuning among the same structurally-defined odorant classes.

**Figure 4.**
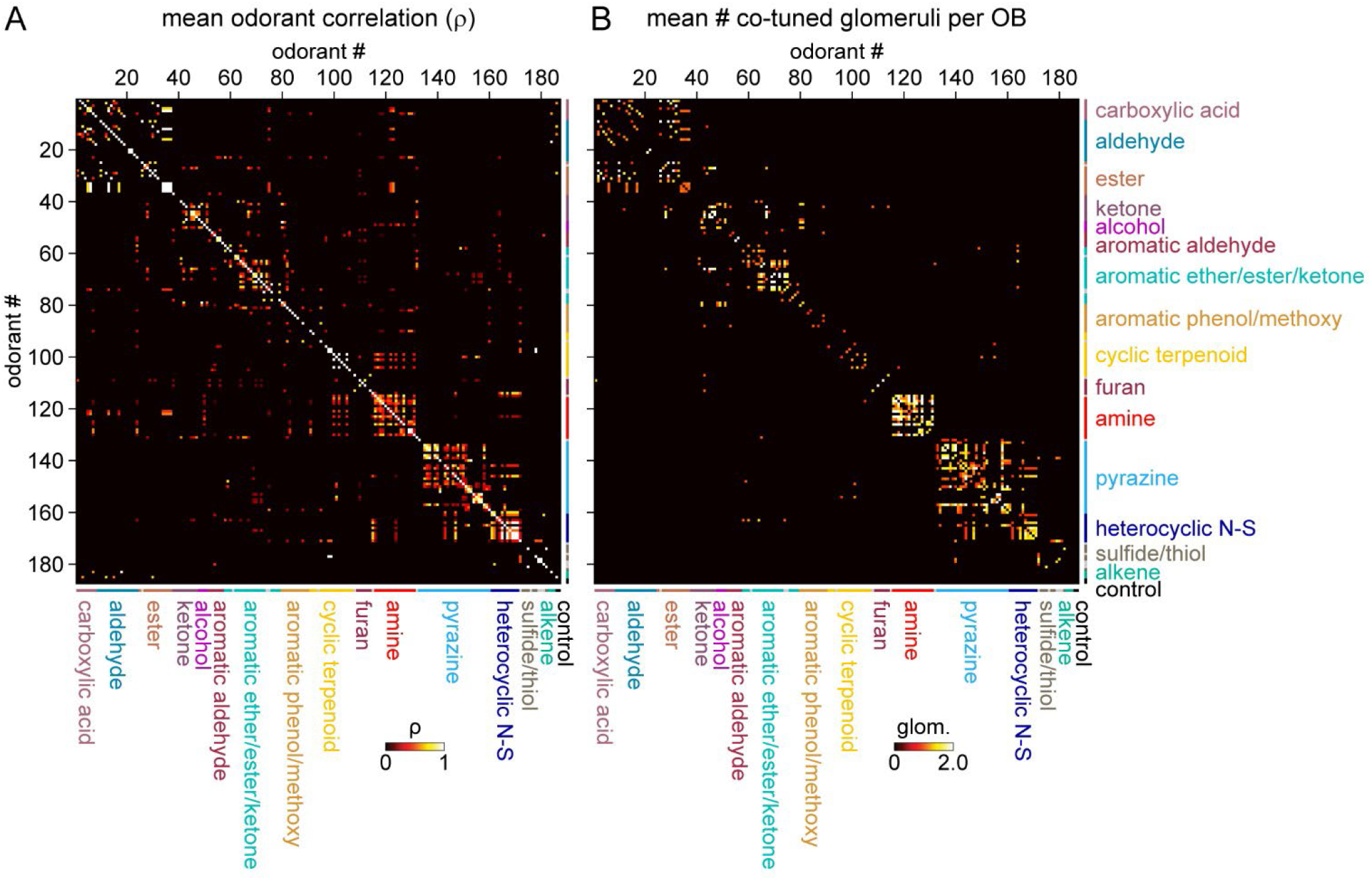
Sparse tuning of glomerular inputs is heterogeneously structured. **A**. Mean glomerular response correlation matrix for the 185-odorant panel. Values show Spearman’s rank correlation (ρ), averaged across all 8 OBs. Odorants ordered by structural classification as in previous Figures. **B**. Odorant co-tuning matrix for the 185-odorant panel. Values show mean number of glomeruli co-activated by each odorant pair; matrix is thresholded at 0.875 (<1 co-tuned glomerulus per OB) to highlight the strongest odorant co-tuning relationships.

For further visualization we generated a network graph of odorant co-tuning relationships, using co-tuning as the adjacency matrix input, with odorants as nodes and mean number of co-tuned glomeruli as edge weights (**Fig. 5A**). This visualization confirmed that glomerular tuning was dominated by particular structural relationships between co-tuned odorants. In particular, there was prominent co-tuning between carboxylic acids, aldehydes and esters; between primary amines – including both acyclic and cyclic amines; and between the heterocyclic pyrazines, thiazoles and pyrroles/pyridines (all heterocyclic aromatic compounds containing one or two nitrogens or a nitrogen and sulfur heteroatom). Notably, these co-tuning relationships were not well-predicted by the chemical space of odorants defined by computed physicochemical descriptors: for example, the amine odorants were distributed across a large extent of the Good Scents/2982-descriptor space but showed almost no co-tuning to other odorant classes (**Fig. 5A, B**). Conversely, the aromatic-containing odorants overlapped within a relatively small extent of descriptor space, but there was little co-tuning between aromatic compounds with different functional groups (**Fig. 5A, B**). Other notable aspects of glomerular tuning appeared generalizable. For example, co-tuning among the acids, aldehydes and esters was common between odorants with the same or similar carbon chain length (e.g., butyric acid, butyraldehyde, butyric acid esters). However, these glomeruli responded to the acids at 100-1000x lower concentrations than their aldehyde or ester counterparts (**Document S2, Table S1**). In addition, while glomeruli sensitive to the sulfur- and nitrogen-containing thiazoles, pyrazines and pyrrole/pyridines showed extensive co-tuning among these classes, they were not sensitive to sulfur- and nitrogen-containing thiols and amines, respectively. Overall, this analysis reveals a basic structure underlying the tuning of the OR repertoire that largely reflects relatively straightforward structural relationships among odorants spanning chemical space.

**Figure 5.**
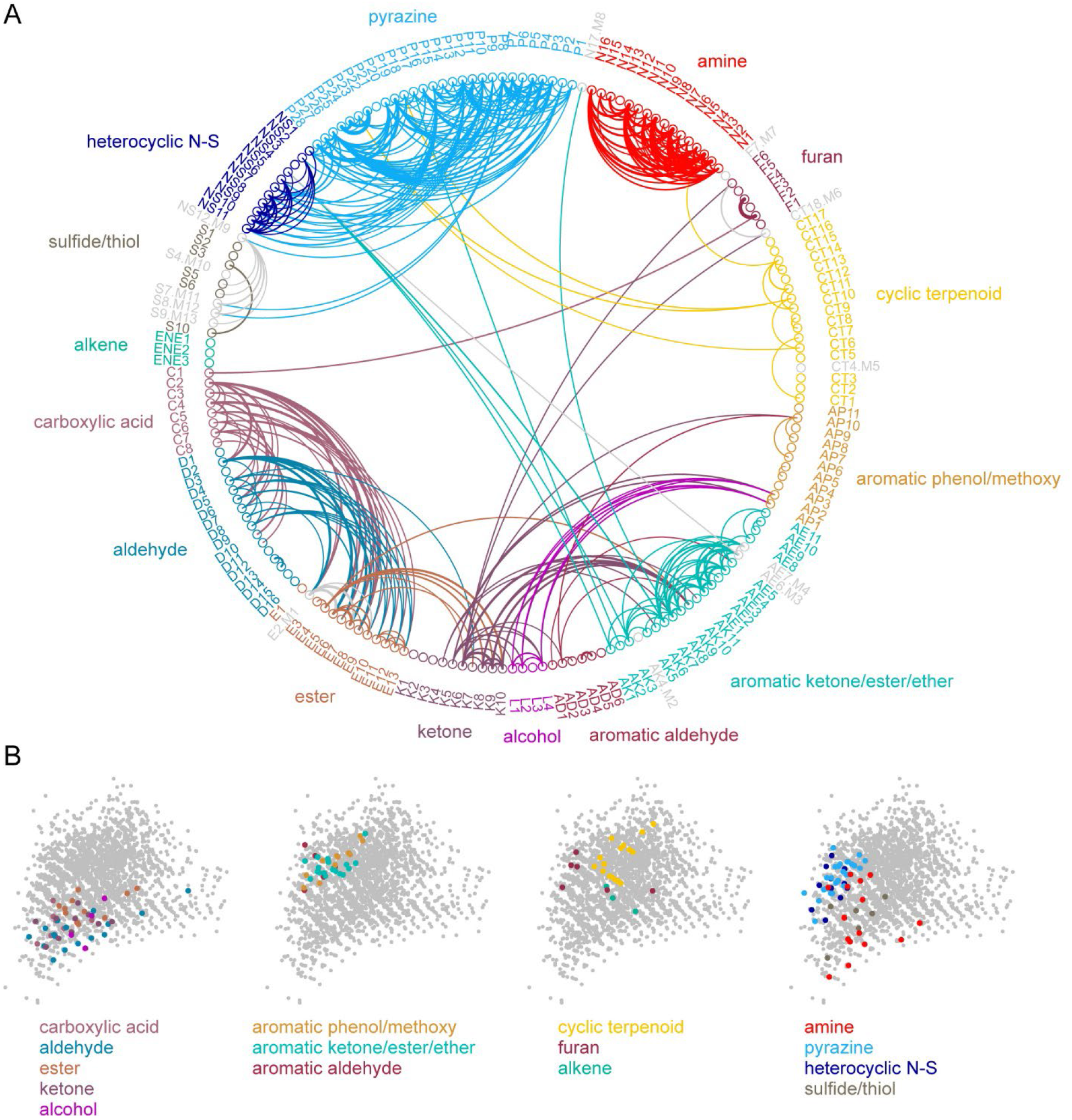
Odorant co-tuning relationships reflect basic chemical features of odorants. **A**. Circular network graph of odorant co-tuning relationships, using the co-tuning matrix from Fig. 4A. Lines connect odorant pairs with mean co-tuning values above 0.875 glomeruli per OB. Line thickness scales with co-tuning value. Colors indicate membership in odorant structural group. Letter-number codes indicate odorant identity (Table S1). Odorants with mixed group-defining structural features are shown in grey. Ordering is the same as in previous Figures. **B**. Odorant panel color-coded by structural group, plotted in the first two principal components of the 2587-odorant physicochemical descriptor space, as in Fig. 1A. Select odorant classes are highlighted in each replicate plot to facilitate visual comparison. Grey circles indicate all odorants in the database.

Finally, we analyzed the spatial organization of glomerular sensitivities with respect to odorant chemical features. Earlier studies have come to varying conclusions about the degree to which chemical features are reflected in the spatial organization of glomerular maps (i.e., ‘chemotopy’) (Takahashi et al., 2004; Johnson and Leon, 2007 Soelter, 2020 #4953; Soucy et al., 2009; Ma et al., 2012; Chae et al., 2019). Here, reasoning that any spatial organization in glomerulus tuning should reflect the features of its highest-sensitivity odorants, and building on the finding that glomerular co-tuning largely reflected gross structural features, we first classified odorants based on these features (i.e., functional group, carbon chain configuration, heteroatom substitution), and then assigned each glomerular position to the structural class of its primary odorant (**Fig. 6A**). We then used point-pattern analysis (Ripley’s K) to test for a nonrandom spatial distribution of glomeruli tuned to odorants within each class (see Methods). Of the 16 odorant classes considered, 9 included sufficient numbers of glomeruli in each OB to support a statistical analysis of spatial organization; of these, 7 classes showed significant spatial clustering of glomeruli in every OB; the two remaining classes – aromatic aldehydes and heterocyclic N-S compounds – showed significant clustering in most OBs (6/8 and 5/8, respectively) (**Fig. 6A, B**). Individual glomeruli within a cluster were narrowly tuned to only one or a few odorants within a class, indicating that spatial clustering did not arise from overlapping odorant sensitivities among nearby glomeruli. Spatial clustering was expected for the carboxylic acids and amines, which are preferred ligands for class I ORs and trace amine-associated receptors (TAARs), whose OSN projections selectively target domains in the anterior-medial and central-medial OB, respectively (Bozza et al., 2009; Pacifico et al., 2012; Cichy et al., 2019). Indeed, glomeruli with maximal sensitivities to carboxylic acids and amines showed spatial clustering of glomeruli within these regions (**Fig. 6A, C**). For the 5 remaining odorant classes, clustering was apparent on a spatial scale smaller than the domain defined by the remaining class II OSN projections. For example, pyrazine-sensitive glomeruli were preferentially located in the caudal-lateral most extent of the dorsal OB, while glomeruli sensitive to furans and aromatic compounds were clustered more centrally (**Fig. 6A, C**). However, these chemically-defined clusters were nevertheless highly overlapping in their spatial extent: glomeruli with distinct odorant sensitivities were spatially interspersed (**Fig. 6C**). These results are consistent with a ‘mosaic’ spatial organization of glomerular chemical sensitivities that has been noted previously for a few odorant classes (Soucy et al., 2009; Chae et al., 2019), and suggest that such an organization extends across much of olfactory chemical space.

**Figure 6.**
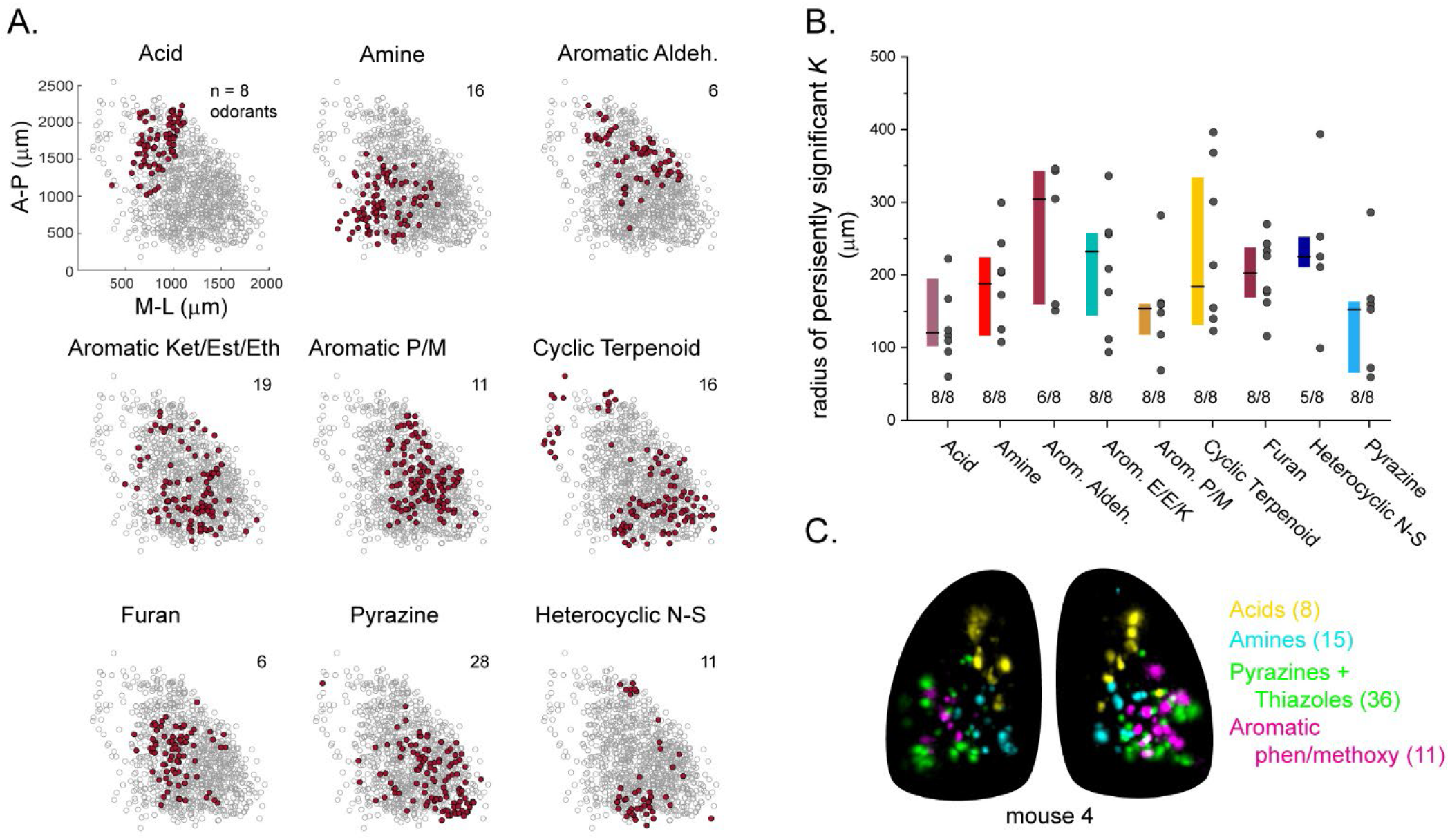
Glomerular sensitivity maps reveal spatial clustering of glomeruli tuned to odorant structural classes. **A**. Glomerular positions highlighted across all 8 OBs (grey), plotted separately and identified by the structural class of their primary odorant (red). Numbers indicate odorants in each functional class. **B**. Size of statistically significant spatial clusters for glomeruli in each odorant class. Size plotted as r, the minimal radius (in μm) at which Ripley’s K remained significant at p<0.01 as r was progressively increased. Numbers below each bar indicate number of OBs (out of 8) showing statistically significant clustering. Boxes show median and interquartile ranges. Colors match those from Fig. 4C. **C**. Maximal projection overlays of response maps evoked by odorants within four distinct structural classes: carboxylic acids, amines, pyrazines/thiazoles, and phenol/methoxy-containing aromatics. Numbers indicate number of odorants tested within each class. All data taken from the same mouse. Individual glomeruli show little to no co-tuning to odorants in different classes.

## DISCUSSION

By mapping high-sensitivity odorant responses to OB glomeruli using a large and diverse odorant panel, we found that individual glomeruli – and, by inference, their cognate ORs – are sensitively and selectively tuned to a very narrow portion of odorant chemical space. This narrow tuning leads to a sparse and apparently high-dimensional representation of individual odorants in the regime of low odorant concentrations. At the same time, the small amount of structure that was apparent in this coding regime – both in terms of glomerular tuning and glomerular location – was well-represented by relatively simple descriptions of chemical space derived from odorant structural features.

The prevalence of narrowly-tuned glomeruli, and the degree of selectivity in their tuning, was striking. While some earlier characterizations of OSN tuning have concluded that the majority of OSNs (and, thus, the ORs they express) are narrowly tuned to odorants with shared structural motifs (Araneda et al., 2000; Nara et al., 2011) - a result qualitatively similar to that seen here - most prior studies have reported a combination of broadly- and narrowly tuned OSNs (Hallem and Carlson, 2006; Grosmaitre et al., 2009; Saito et al., 2009; Nara et al., 2011), with even those OSNs classified as narrowly tuned appearing less selective than the majority of glomeruli in our dataset. Likewise, recordings from mitral/tufted cells in the mouse have indicated high odorant selectivity, but reported lifetime sparseness values across a range of odorants are substantially lower (indicating broader tuning) than those found here (Davison and Katz, 2007; Fantana et al., 2008; Tan et al., 2010).

A likely explanation for this difference is the much lower range of odorant concentrations used to characterize glomerular specificity here. In general, we presented odorants at concentrations several orders of magnitude lower than those in earlier in vivo studies (Davison and Katz, 2007; Soucy et al., 2009; Ma et al., 2012; Chae et al., 2019; Pashkovski et al., 2020), and 4-6 orders of magnitude lower than in vitro studies (Saito et al., 2009; Nara et al., 2011; Xu et al., 2020). Nonetheless, we found that glomeruli remained narrowly tuned even when challenged with tenfold higher concentrations, and that narrow tuning was not a function of poor signal-to-noise ratios nor of the particular selection of concentrations used for each odorant. Thus, highly selective tuning to odorants appears a robust feature of OSN input to OB glomeruli.

The high sensitivity of glomeruli to their primary odorant was also surprising: the median primary odorant concentration across all glomeruli was 2 × 10^−11^ M; threshold concentrations for characteristic odorant-glomerulus pairs are likely even lower, as we did not attempt to determine threshold sensitivities for any glomerulus. These concentrations are comparable to the sensitivity of the ‘ultrasensitive’ TAAR OSNs for their preferred ligands (Zhang et al., 2013), and substantially lower than those reported for other OSNs in vivo (Oka et al., 2006; Tan et al., 2010). This discrepancy could be explained by the smaller odorant panels used in those studies, resulting in a failure to find optimal odorants for a given OSN/OR, as well as a reliance on less-sensitive reporters in detecting responses. We note that the concentrations used here were comparable to or higher than psychophysical detection thresholds measured with rigorous behavioral assays in mice (Dewan et al., 2018; Cichy et al., 2019; Williams and Dewan, 2020), suggesting that the sparse responses at these concentrations can support odor perception.

The narrow tuning of OSNs has implications for how odorant identity is encoded in low-concentration regimes. Canonical models of olfactory information processing rely on a reduced dimensionality of the odorant coding space that arises from systematic overlap in the response spectra of OSN inputs, and predict that central circuits transform odorant representations with respect to this lower-dimensional space (Chae et al., 2019; Pashkovski et al., 2020). Given the low covariance in responses we observed across the OSN population for different odorants, such models may be less applicable in low-concentration regimes. Instead, activation of particular glomeruli may directly encode the presence of a particular odorant, or at least one among a small number of odorants with shared structural features.

Natural odors typically consist of mixtures of many components; in a sparse coding regime, natural odors may elicit combinatorial activity patterns that directly reflect their component composition, allowing for both a configural recognition of odor ‘objects’ based on co-active glomeruli as well as an analytic processing of individual mixture components (Coureaud et al., 2022). Previous studies using higher odorant concentrations have found a limited range of analytical abilities for odorant mixtures that decreases with increasing overlap in glomerular activity evoked by each component (Rokni et al., 2014; Qiu et al., 2021). Based on the present results, we suggest that odorant mixture analytical capacity may be more robust in low-concentration regimes. Narrowly-tuned, highly-sensitive glomeruli may also facilitate the recognition of a complex natural odor at trace concentrations based on detection of key components that are signatures of a given odor source (Dunkel et al., 2014).

Despite the overall sparse representation of odorants, we did find systematic relationships in the co-tuning of individual glomeruli to their highest-sensitivity odorants. Co-tuning relationships largely reflected straightforward features of odorant chemical structure: predictive models that incorporated a relatively small number of such structural features outperformed models generated from larger sets consisting of exhaustive lists of computationally-derived physicochemical properties, and performed similar to or better than reduced descriptor subsets previously optimized to explain OR-odorant interactions by fitting to response data (Haddad et al., 2008). Physicochemical descriptor sets – which are heavily used in computational drug discovery – have been moderately successful at predicting OR ligands and odorant perceptual features, but typically require careful tuning of feature selection to training datasets (Boyle et al., 2013; Ravia et al., 2020; Gerkin, 2021; Kowalewski et al., 2021) and introduce hazards such as overfitting a large parameter set to a much smaller response dataset (Chae et al., 2019). In addition, odorants can differ significantly from drug-like molecules in their size and other chemical features (Ruddigkeit et al., 2014). Thus, it is encouraging that the SMARTS and MACCS fingerprints used here were equally or more effective at predicting the highest-sensitivity odorants for a given OSN population, as they reflect clear substructural features of odorant molecules and required many fewer descriptors than odorants. These substructure-based models may generalize well across odorant chemical space or across different response measures. At the same time, emphasizing the highest-sensitivity odorants for a given OSN population revealed prominent co-tuning relationships among particular odorant groups, indicating a heterogeneous structure to odorant coding space at the level of OSNs that is not adequately captured by structure-based or computationally-derived chemical features. Taking into account the chemical relationships underlying this structure could inform predictive models of odorant-OR binding or odor perception, and be useful for more precisely understanding how central circuits transform odorant representations (Poivet et al., 2018; Licon et al., 2019; Kowalewski and Ray, 2020).

Structure in odorant representations was also apparent in the spatial organization of glomerular sensitivities with respect to odorant chemical features. We found that spatial clustering of glomeruli with high sensitivities to structurally similar odorants was common, being present in nearly all of the structurally-defined classes tested. These results differ from those of earlier studies mapping chemical features across the OB surface, which used higher odorant concentrations and metabolic measures of neural activity and reported relatively discrete spatial clusters of glomeruli responsive to odorants sharing particular chemical features (Takahashi et al., 2004; Johnson and Leon, 2007). In particular, spatial clustering in our dataset largely arose from the proximal positioning of individual glomeruli sensitively tuned to structurally similar but distinct odorants, rather than from correlated tuning to overlapping sets of odorants or a ‘tunotopic’ organization (Ma et al., 2012). Moreover, with the exception of carboxylic acid-sensitive glomeruli, which were tightly clustered within the presumed domain of class I OSN projections (Tsuboi et al., 2006; Bozza et al., 2009), spatial clustering of glomerular sensitivities was not discrete, but overlapping and interdigitated. This finding is consistent with the functional mosaic organization of glomerular responses noted previously for aldehydes and thiazoles (Soucy et al., 2009; Chae et al., 2019), and suggests that this organization is a general feature of glomerular maps with respect to odorant chemical space.

Mapping high-sensitivity odorants to individual glomeruli allowed for the straightforward functional identification of glomeruli across the dorsal OB using only a single diagnostic odorant at a low concentration. Previous studies have functionally identified glomeruli using a combination of location and response spectrum across multiple odorants (Wachowiak and Cohen, 2001; Soucy et al., 2009; Soelter et al., 2020), and have used these to link glomeruli to their cognate ORs (Oka et al., 2006; Shirasu et al., 2014). However, broader use of functionally-identified glomeruli has been limited thus far. Here, we identified at least 26 glomeruli that could be confidently identified - in both anesthetized and awake mice - using only a single diagnostic odorant at a low concentration; far more could likely be identified using additional criteria such as gross position or a second diagnostic odorant. The resulting lookup table of glomeruli and their diagnostic odorants provides an efficient means of accessing each of these glomeruli.

We used functionally-identified glomeruli to generate consensus response spectra and to assess variability across OBs and animals; variability was low and likely reflected slight inter-animal differences in physiological state and/or interhemispheric differences in sensitivity or nasal patency. Consensus response spectra were useful in comparing responses to predictions from odorant descriptor sets and in confidently identifying co-tuning relationships that deviated from these predictions. Future studies should be able to probe response spectra for these glomeruli over even larger odorant panels, across concentrations, and chronically in the awake animal. Paired with platforms for linking functionally-identified glomeruli to their cognate ORs, such as retrograde labeling of OSNs (Oka et al., 2006; Shirasu et al., 2014; Saito et al., 2017) or spatial transcriptomics (Wang et al., 2022), functionally identifying glomeruli could prove an efficient means of functionally deorphanizing and mapping mammalian ORs. Indeed, the number of functionally-identified glomeruli from our initial conservative survey exceeds the total number of OR-defined glomeruli whose functional properties have been characterized in prior studies to date (Peterlin et al., 2014; Shirasu et al., 2014; Saito et al., 2017). Functionally-identified glomeruli should also prove useful for investigating OB circuit transformations using multiplexed imaging from OSN inputs and OB outputs (Short and Wachowiak, 2019; Moran et al., 2021), as well as plasticity in glomerular responses (Kass et al., 2013; Dias and Ressler, 2014; Liu et al., 2016; Bhattarai et al., 2020).

An important question for interpreting the ethological significance of the present findings is the range of odorant concentrations encountered by the animal during odor-guided behaviors. Quantitative measurements of vapor-phase concentrations arising from natural odor sources under naturalistic conditions are rare, although it is likely that concentrations vary widely and have a long tail towards lower concentrations as odorants are dispersed by airflow with distance from their source. Even at the source, however, concentrations of many odorants appear to fall within the range of those tested here. For example, ambient concentrations of the most abundant species of volatile organic compounds measured in high-density plant environments range from 0.1 – 10 ppb (Petersson, 1988; Jansen et al., 2009; Bach et al., 2020). Likewise, reports of odorant flux rates from sources such as tomato leaf, forest floor, and freshly cut grass lead to concentration estimates within a similar range when the source is directly sampled via sniffing at its surface (Ruuskanen et al., 2011; Mäki et al., 2019; Dehimeche et al., 2021). Thus, while we have referred to the concentrations used in the present study as low relative to those used in most previous studies, this range likely reflects a common operating regime of the mouse olfactory system during natural behavior.

## MATERIALS AND METHODS

### Animals

Experiments were performed using both male and female compound heterozygous crosses of OMP-IRES-tTA (Jackson Laboratory stock #017754) (Yu et al., 2004) and tetO-GCaMP6s (Jackson Laboratory stock #024742) (Wekselblatt et al., 2016) mice aged 2-6 months. Mice were housed up to 5 per cage on a 12-hour light/dark cycle with food and water available ad libitum. All procedures were performed following the National Institutes of Health Guide for the Care and Use of Laboratory Animals and were approved by the University of Utah Institutional Animal Care and Use Committee.

### Olfactometry

Odorants were obtained from Sigma-Aldrich, TCI America, Bedoukian Research, or ICN Biomedicals. Liquid dilutions of odorants were prepared to achieve target delivery concentrations of approximately 0.1, 1, 10, 100, or 1000 pM (within an order-of-magnitude) using 1:10 and 1:100 serial dilutions. Non-amine odorants were diluted in caprylic/capric medium chain triglyceride oil (C3465, Spectrum Chemical Mfg. Corp.) within ~1 week of experiments; amine odorants were freshly diluted in water immediately prior to each experiment to minimize odorant oxidation. Trace quantities of Sudan Black B were included in all dilutions to facilitate visual confirmation of olfactometer loading. Diluted odorants were delivered in vapor phase using a custom-built olfactometer equipped with end-stage eductor and operating with 8-L/min charcoal-filtered carrier stream, 30-kPa delivery pressure, 5-cm olfactometer-to-mouse distance, and 2-s long delivery (Burton et al., 2019). A fan at the rear of the animal removed odorants after presentation. Odorants were delivered independent of inhalation timing in pseudorandom order, typically in sets of 12 odorants, with 3-5 trials per odorant and 8-10 s inter-trial interval. Eductors were washed with non-scented Alconox detergent and thoroughly rinsed with ethanol and water in between odorant sets to minimize possible odorant adsorption and inter-trial contamination.

### Imaging

Mice were initially anesthetized with intraperitoneal injection of pentobarbital (50 mg/kg) and subcutaneous injection of chlorprothixene (12.5 mg/kg). Subcutaneous injection of atropine (0.5 mg/kg) was further given to minimize mucus secretions and maintain nasal patency. A double tracheotomy was performed as described (Eiting and Wachowiak, 2018), and anesthesia was subsequently maintained by ~0.4-0.5% isoflurane delivered in pure O_2_ to the descending tracheal tube while artificial inhalation (150-ms duration, 300-mL/min flow rate) was continuously driven at 3 Hz through the ascending tracheal tube. Mice were then head-fixed and the bone over the dorsal OB thinned. For widefield imaging experiments, a large well surrounding the dorsal OB was constructed with dental cement and securely covered with a cut glass coverslip, forming a chamber with caudal opening. This chamber was filled with Ringer’s solution to render the thinned bone transparent, yielding a cranial window with stable optical plane throughout the experiment. Epifluorescence was collected through a 4×, 0.28 N.A. air objective (Olympus) at 256×256-pixel resolution and 25-Hz frame rate using a back-illuminated CCD camera (NeuroCCD-SM256; RedShirt Imaging) and Neuroplex software, with illumination provided by a 470 nm LED (M470L2, Thorlabs) and green fluorescent protein filter set (GFP-1828A-000, Semrock). For experiments involving both widefield and two-photon imaging trials, epifluorescence was collected similar to above using a 5×, 0.25 N.A. air objective (Olympus), while two-photon fluorescence was collected with 15.2-Hz frame rate using a resonant-scanning microscope (Sutter Instruments) coupled to a pulsed Ti-Sapphire laser (Mai Tai HP, Spectra Physics) tuned to 920 nm and equipped with a 16×, 0.8 N.A. water-dipping objective (Nikon) and GaAsP photomultiplier (Hamamatsu H10770B).

For epifluorescence imaging in awake, head-fixed mice (**Fig. S5**), the bone overlying both OBs was thinned and sealed with cyanoacrylate (Krazy Glue) to preserve transparency and a headbar was implanted caudal to the OBs (Wachowiak et al., 2013). Beginning 3 – 5 days after the surgical procedure, mice were acclimated to head fixation for periods increasing from 10 – 30 minutes over 3 – 4 days. A circular treadmill (design courtesy of D. Rinberg, New York University) allowed for locomotory movements and minimized torque on the headbar. Odorant-evoked responses were imaged on the same optical setup and using the same odorant delivery paradigm as in anesthetized mice. Data were collected over 2 – 3 consecutive daily sessions lasting 45 – 60 minutes.

### Odorant response maps

To generate odorant response maps, raw data from the ≥3 presentations of each odorant were first averaged, then ΔF images generated by, for each pixel, subtracting the mean of the fluorescence signal in the 1 second prior to odorant delivery onset from the mean signal in seconds 2-3 after odorant delivery onset. For presentation (e.g., **Document S2**), the resulting ΔF response maps were clipped at zero and a maximum equal to the mean of the highest 65 pixels, smoothed slightly by convolving with a 2D Gaussian kernel (sigma: 0.75 pixels), and the pixel resolution doubled (to 512 × 512 pixels) with bilinear interpolation. Response maps were displayed and analyzed in units of ΔF, rather than ΔF/F, due to the difficulty in precisely measuring baseline fluorescence for single glomeruli under epifluorescence, and to evaluate absolute response magnitudes rather than responses relative to pre-odorant spontaneous activity. However, repeating all analyses using ΔF/F measurements yielded near-identical results (not shown).

### Signal extraction and segmentation

For the main dataset (‘1x concentrations’, widefield epifluorescence), regions of interests (ROIs) representing single glomeruli were generated using an initial automated selection process followed by manual refinement. First, maximal ΔF projections were generated from the raw (unsmoothed) ΔF response maps across all odorants and initial ROI boundaries generated using ‘Find Circles’ in Matlab’s Image Segmenter App, followed by the ‘bwconncomp’ function. This initial ROI set was manually refined by addition or adjustment based on visual inspection of the maximal projections and, in some cases, individual odorant response maps. The mean ΔF signal of all pixels from each ROI was used to generate a response matrix (ROI × odorant) for each OB.

Response matrices derived from epifluorescence data required further segmentation to minimize effects from signal spread due to scattered or out-of-focus light arising from nearby glomeruli. Rather than attempt automated segmentation (Soelter et al., 2014), we verified that responses were accurately attributed to a given ROI by visual inspection. The inspection procedure involved sorting odorant response maps in descending order of their ΔF response for a given ROI, and setting to zero any entries in the response matrix that did not correspond to clear ΔF signal foci that were centered within the ROI of interest. Signals attributable to responses in adjacent ROIs were easily distinguished with this approach. Due to the sparse nature of odorant responses, the inspection algorithm required visual inspection of, typically, 5-20 response maps per ROI. Inspection was performed blind to odorant identity. In rare cases where odorants evoked clear activity that was not easily segmented into glomerular foci (e.g., octanal response maps, **Document S2**), no ROI was chosen and these signals were not included for analysis.

For the 10x concentration dataset acquired with two-photon imaging, ROIs were selected manually from maximal ΔF projections, and the resulting response matrices were thresholded using a modified z-score cutoff based on the variance in non-responsive odorant trials for each ROI. Variance was calculated as the standard deviation of ΔF response values, excluding the top 10th and bottom 5th percentiles of responses across the 185-odorant spectrum. Responses corresponding to a z-score <9 were set to zero, and the resulting thresholded ΔF response matrix was used for further analysis.

### Response matrix statistical measures

Lifetime sparseness (S_L_), a measure of tuning of each glomerulus across the odorant response panel, was calculated as previously (Davison and Katz, 2007; Schlief and Wilson, 2007; Pashkovski et al., 2020). For a set of responses to a single glomerulus across n odorants (R={r_1_, r_2_, … r_n_})

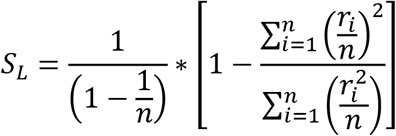

Similarly, population sparseness (S_P_), a measure of sparseness of n glomerular responses to each odorant (G={g_1_, g_2_, … g_n_}), was calculated as

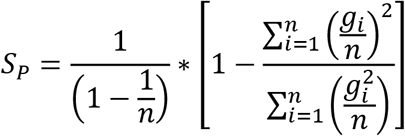

For both measures, a value of 1 indicates maximally selective tuning (a glomerular response to only one of the 185 odorants (S_L_), or a single glomerulus activated by an odorant (S_P_), while a value of 0 indicates completely nonselective tuning, i.e., equivalent responses to all odorants or equivalent activation of all glomeruli.

Principal component analysis (PCA) was performed separately on the response matrix of each OB after normalizing each glomerulus’s response spectrum to its maximal response. Effective dimensionality (ED; a measure of the number of principal components (PCs) required to explain a fraction of the variance in the response dataset) was calculated as described in (Litwin-Kumar et al., 2017; Pashkovski et al., 2020). To compare datasets with different numbers of glomeruli (and, thus, a different maximal number of PCs), we expressed variance explained as a fraction of the total number of PCs.

### Functional identification of glomeruli

To define odorants that could be reliably used to identify putatively cognate glomeruli across OBs, we first identified odorants that reliably activated singular or near-singular activation of glomeruli using two criteria: no more than two glomeruli activated above a 50% ΔF_max_ cutoff across all imaged OBs, and activation of at least one glomerulus in at least 6 of 8 OBs, and in each of the 4 mice. Next, for each potential diagnostic odorant, we identified the glomerulus maximally activated by that odorant in each imaged OB and compared the response spectra of each of these glomeruli across the full 185-odorant panel using Pearson’s correlation, taking the median of all pairwise correlations across the 6-8 OBs as a measure of consistency of response spectrum. We also measured the frequency with which the response spectrum of the maximally-activated glomerulus was the most the highly correlated with that of the maximally-activated glomerulus in other OBs; this fraction reflected the reliability with which the strongest-responding glomerulus to a given odorant also identified the glomerulus with the most similar response spectrum in another OB. We defined an error ratio as 1 minus this fraction. Odorants were considered diagnostic for functionally-identified glomeruli if they had a median correlation coefficient > 0.8 and an error ratio < 0.2.

### Odorant classification

To facilitate visual comparison of response maps across the large odorant panel, odorants were nominally grouped according to common structural classifiers including functional group, aliphatic, aromatic or cyclic structure, heteroatom substitution, etc. Odorants with mixed features are identified as such in **Table S1**. Within each group, odorants were ordered according to progressive changes in particular features such as carbon chain length. The ordering process was subjective and used only for data visualization. The structural classification was used only for analysis of spatial clustering (e.g., **Fig. 6**), as described below.

### Odorant concentrations

Concentrations are reported as estimated concentration delivered to the mouse nose based on the vapor pressure of each odorant, its dilution in liquid solvent, and prior calibrations of the olfactometer using a photoionization detector (Burton et al., 2019). We assumed that the odorant and solvent behave as an ideal mixture and that odorants reach saturated vapor concentrations in their delivery reservoir as predicted by Raoult’s Law; that is, the odorant and solvent are chemically nonreactive with each other and totally miscible at all proportions. This assumption will not hold across all odorants, but was necessary as data on activity coefficients that describe deviations from ideal behavior are extremely limited for these systems. Furthermore, calculations were based on estimated vapor pressures at 25 C calculated using the EpiSuite ‘mpvpbp’ module (US Environmental Protection Agency, https://www.epa.gov/tsca-screening-tools/epi-suitetm-estimation-program-interface) or as reported by The Good Scents company; reported vapor pressures can vary by 30 - 60% depending on the source and the estimation method. Given these considerations, all reported final concentrations should be considered estimates, and are reported in **Document S1** to the nearest order of magnitude precision.

For comparison of concentrations with prior studies (**Fig. S2**), we compared only odorants in common with our odorant panel. For comparison to (Pashkovski et al., 2020) and (Davison and Katz, 2007), we used their reported values of 100 ppm and 10 ppm, respectively. For (Soucy et al., 2009; Chae et al., 2019) we used their reported estimate of 1% of saturated vapor of a 1:100 dilution in mineral oil to arrive at a total dilution of 0.01% saturated vapor and calculated estimated delivered concentrations as described above. For (Ma et al., 2014) we used the lowest reported effective dilutions of saturated vapor from the ‘GIA0512’ dataset.

### Chemical descriptor sets

Glomerular response spectra were analyzed with respect to various odorant physicochemical descriptor sets, chosen from previous studies (see Text) and processed as follows: The ‘Dragon 1377’ set was identical to that used in (Haddad et al., 2008; Saito et al., 2009; Chae et al., 2019; Gerkin, 2021) and consisted of an initial set of 1664 descriptors taken from the Dragon database (Kode Systems), 1377 of which showed non-zero variance across our odorant panel. The ‘Dragon 32’ set consisted of the subset of 32 descriptors (a subset of the original 1664) found to optimally predict odorant-OR interactions (Haddad et al., 2008). The ‘FG’ descriptor set consisted of the subset of 154 descriptors defining odorant functional groups (Saito et al., 2009), 49 of which had non-zero variance across our odorant panel. The ‘Alvadesc 2982’ set was generated from an expanded list of 5666 descriptors added to the original Dragon database and accessed using the Alvadesc interface (Kode Systems), 2982 of which showed non-zero variance across our odorant set. This descriptor set included the same descriptors used in (Pashkovski et al., 2020) (2873 descriptors), with additional descriptors reflecting the larger number of odorants in our panel. For all of the above descriptor sets, descriptor values were extracted from the current Alvadesc database. Then, to normalize for differences in ranges and units across the descriptor set, values for each descriptor across the odorant panel were z-scored, as done previously (Saito et al., 2009; Chae et al., 2019; Pashkovski et al., 2020).

We also defined odorant distances using principal components of the chemical space defined by the 2982 descriptors across 2587 odorants, as performed previously (Pashkovski et al., 2020); 2548 odorants were taken from the Good Scents database and met molecular weight criteria for odorous compounds (m.w. between 50 and 300); the remaining 39 were in our odorant panel but not present in this original list and so were added. Parameter values were z-scored as above before performing PC analysis. The first two PCs, accounting for 39.5% of the total variance, were used to visualize odorants in chemical space (**Fig. 1A**). We used the weighting of each odorant across the first 20 PCs, which accounted for 66% of the total variance across the odorant set, to generate a 20×185 matrix describing the relative position of each odorant in this reduced chemical space.

Odorant distances were also defined using binary feature keysets representing the presence or absence of a particular chemical feature. ‘MACCS’ keys (Durant et al., 2002), 59 (of 166 total keys) of which were present in our odorant panel, were extracted using Alvadesc. The ‘SMARTS42’ key set was defined using the SMARTS chemical pattern matching language (Daylight Chemical Information Systems Inc.; https://www.daylight.com/dayhtml/doc/theory/theory.smarts.html), as described in the Text. The ‘SMARTS42’ set consisted of 42 features (**Table S4**) which were chosen to capture substructures that are present in common-use odorants, including those in our panel (e.g., carboxyl, aldehyde or ketone group; ester bonds; aromatic rings; pyridine substructure; ortho-, meta- and para-substituted rings; a number of terpenoid scaffolds; and the explicit inclusion of single-bond aliphatic chains of lengths from 6 to 11 carbon atoms). In all of the above cases, pairwise (185 × 185) odorant distance matrices were generated for each odorant-descriptor set, using cosine distance as the distance metric for the z-scored physicochemical descriptor sets and (1 minus dice similarity) for the binary feature keys.

A performance metric derived from receiver-operating characteristic (ROC) analysis, an approach commonly used to evaluate the quality of predictive models for receptor-ligand interactions (Lopes et al., 2017), was used to evaluate the quality of the different physicochemical descriptor sets as models for predicting glomerular odorant responses. For a given glomerulus, an initial ‘query’ odorant was defined as the odorant to which that glomerulus responded at the lowest concentration (i.e., primary odorant) or, in some cases (e.g., **Fig. 3C**, **Fig. S7A**) the odorant that elicited the strongest response. The remaining 184 odorants were sorted in order of increasing distance from the query odorant based on a given descriptor set, and the cumulative fraction of all odorant responses explained as a function of odorant number was plotted, generating a model performance curve, with odorant # as the false positive rate and fraction of responses explained as the true positive rate. This curve was compared to that derived from the measured responses, generated by sorting odorants in the order of their responsiveness. The performance metric, *P*, was defined as

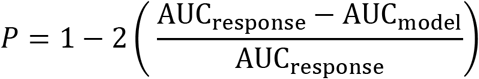

where AUC_response_ is the area under the cumulative response curve and AUC_model_ is the area under the cumulative model performance curve. A model perfectly predicting performance had a value of 1, and chance performance had a value of 0. For measuring model performance for the 19 identified glomeruli with non-singular tuning, median response spectra across all responsive OBs were used, and the query odorant was the primary odorant, with response spectra binarized before evaluation. For comparing performance using the strongest-activating odorant, odorants were sorted in descending order of their response amplitudes and odorant number was plotted against the cumulative fraction of the summed response amplitudes explained. For measuring performance for individual glomeruli across the full dataset, responses were thresholded at 10% of the maximal response for each glomerulus, binarized, and analysis performed only on glomeruli responding above threshold to more than one odorant (694 total glomeruli). For each glomerulus, the performance metric was calculated using each effective odorant as the query odorant (rather than the primary odorant) and the median of these values was reported.

### Response matrix correlation and co-tuning analysis

Odorant response correlation matrices were generated by calculating rank correlation (Spearman’s ρ) across all glomeruli in a response matrix for all odorant pairs; the resulting odorant correlation matrices were averaged across the 8 imaged OBs. Because odorant response vectors typically included only a few responsive glomeruli, we also calculated pairwise odorant co-tuning matrices using a binary measure of responsiveness, where the co-tuning index was equal to the number of glomeruli responsive to a given odorant pair, at any magnitude. Pairwise co-tuning matrices were then averaged across the 8 OBs and thresholded at 0.875 (corresponding to an average of at least one co-tuned glomerulus in 7 of the 8 OBs) to highlight the most consistent odorant co-tuning relationships. Co-tuning relationships were then visualized as a circular network graph using the mean, thresholded co-tuning matrix as the weighted adjacency matrix input. The network graph was rendered using the ‘circulargraph’ function in Matlab (https://www.mathworks.com/matlabcentral/fileexchange/48576-circulargraph/).

### Spatial clustering analysis

To test for nonrandom spatial distribution of glomerular odorant sensitivities, we used the spatial point pattern measure Ripley’s K, which reports whether points within a given radius, r, are dispersed, clustered, or randomly distributed (Ripley, 1977). Within each preparation, each glomerulus was assigned to an odorant group based on our structural classification of its primary odorant and Ripley’s K was calculated using the set of glomeruli in a given group. Glomeruli with primary odorants having mixed classification (16 of the 185 odorants; see **Table S1**) were excluded from analysis. Each imaged OB was analyzed separately and treated as a replicate for statistical analysis. A minimum of 3 glomeruli per structural group in each OB were required to support statistical tests; 9 of the 16 groups met these criteria. For these groups, K was calculated for increasing radius (r) values from 0 to ~400 μm (depending on the largest spread of glomeruli in each dataset), in 512 steps of ~ 0.7 - 0.8 μm, using the R package ‘spatstat’ (http://spatstat.org/). Statistical significance of K at each r was calculated by comparing to a Monte Carlo distribution of K values calculated from the same number of glomeruli randomly chosen from the full glomerular array (10,000 iterations). P-values were taken directly from the probability distribution of the Monte Carlo simulation. We used an alpha level of p<0.01 to identify the radius at which nonrandom spatial distributions of glomeruli existed in each dataset; significant Ripley’s K values were always greater than the random distribution, indicating spatial clustering. To prevent over-interpreting spurious significant values of r, we further required that Ripley’s K remain significant at all increasing radii to consider spatial clustering to be meaningful at a given radius.

## Supporting information

Supplemental Document S2, Complete Odor Map Atlas

Supplemental Document S3, Atlas of Functionally Identified Glomeruli

## DATA AVAILABILITY

Data underlying all analyses, including response matrices for all imaged OBs, are available on request at https://www.wachowiaklab.org/data-request.

## FUNDING

National Institute of Mental Health F32MH115448 (SDB)

National Institute of Neurological Disorders and Stroke R01NS109979 (MW)

National Science Foundation 1555919 (MW)

National Science Foundation/DFG Next Generation Networks for Neuroscience Award 2014217 (MW, MS)

## ACKNOWLEDGEMENTS

We thank Rebecca Kummer, Gustavo Opazo-Vasquez and Alex Price for technical support, Alex Holtmann (University of Utah) for advice on point pattern statistics, Hiro Matsunami (Duke University) for advice on physicochemical descriptors, and Alla Borisyuk and Daniel Zavitz (University of Utah) for help implementing dimensionality measures. We thank C. Ron Yu (Stowers Institute, Univ. of Kansas) for providing OMP-IRES-tTA mice and Stan Pashkovski (Harvard Medical School) for providing the list of Good Scents odorants.

## AUTHOR CONTRIBUTIONS

SDB designed and performed experiments, analyzed data, wrote the manuscript; AB, IA, TPE analyzed data; TCR wrote software for analysis and visualization; MS performed data analysis and developed the SMARTS42 keys; MW designed experiments, analyzed data, wrote the manuscript.

