## Supplemental Document S1, Suppl. Figures and Tables for "Mapping odorant sensitivities reveals a sparse but structured representation of olfactory chemical space by sensory input to the mouse olfactory bulb"

### Supplementary Document S3, Burton et al.

**Supplementary Figures S1 – S7**

**Supplementary Tables S1 – S4**

**Supplementary References**

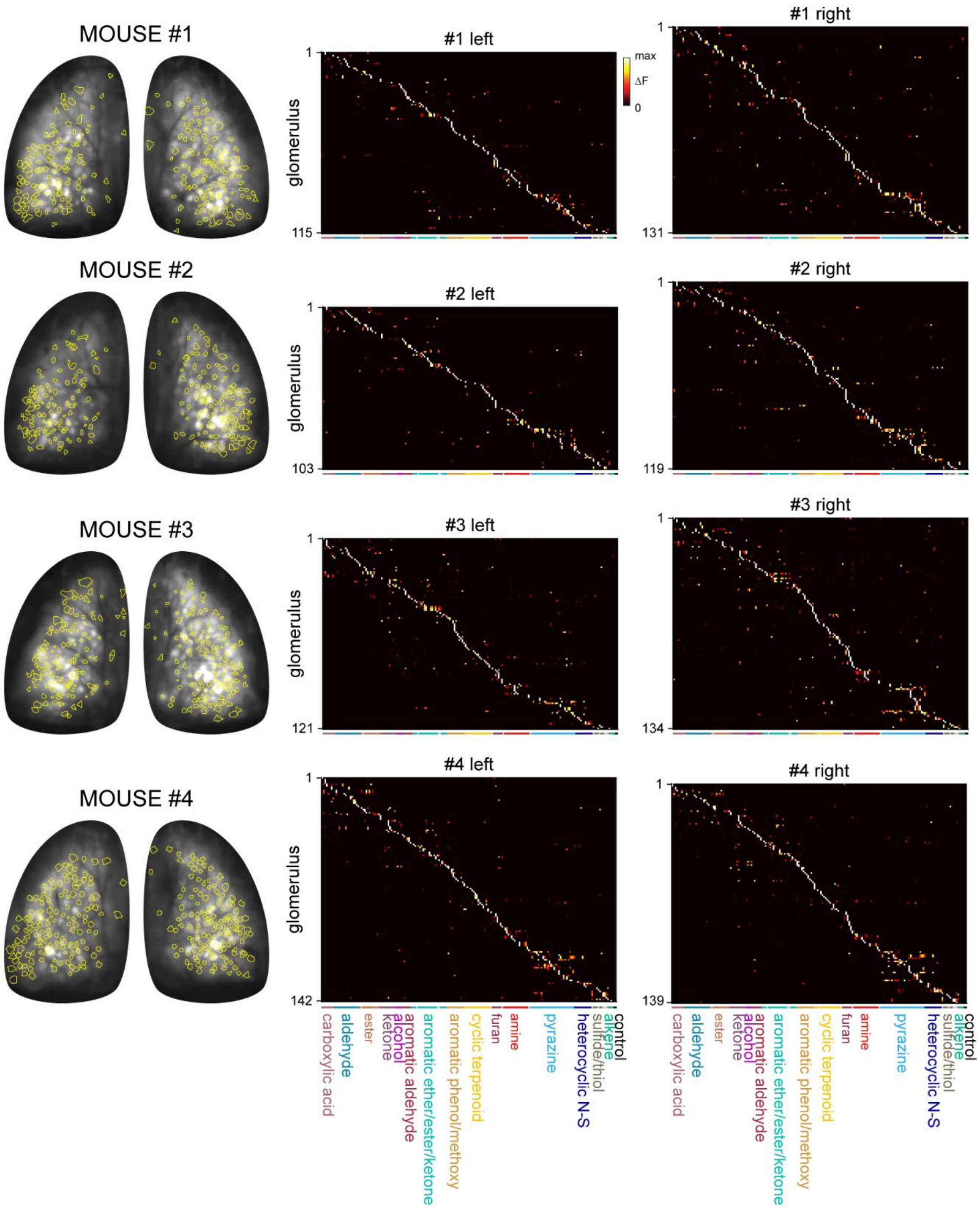


**Figure S1. Summary of responsive glomeruli and their response spectra.**

**Left:** Overlay of all ROIs indicating odorant-responsive glomeruli (yellow) and baseline fluorescence for the four mice imaged under widefield epifluorescence.
**Right:** Matrices of responses across all responsive glomeruli and odorants in each OB after manual segmentation as described (see Methods). Glomeruli and odorants are sorted as in Fig. 1E and occur in the same order as in Table S1. Naming and color-coding of structural classes matches those in Fig. 5A.

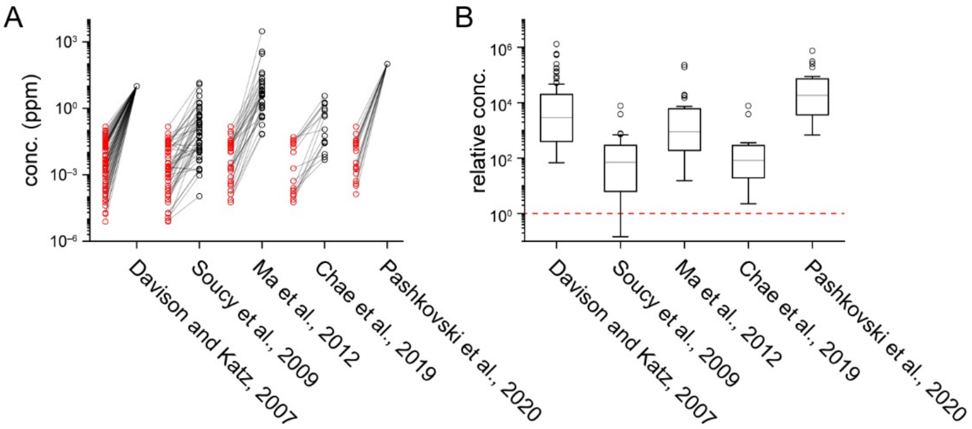


**Figure S2. Comparison of delivered odorant concentrations across studies.**

**A.** Pairwise comparison of delivered odorant concentrations in the current study (red circles) vs. select previous studies (Davison and Katz, 2007; Soucy et al., 2009; Ma et al., 2012; Chae et al., 2019; Pashkovski et al., 2020). Only odorants common across each pair of studies are plotted.
**B.** Distribution of the delivered concentration of each odorant in previous studies relative to the concentration of the same odorant used in the current study. Dashed red line marks equal concentrations across the previous and current study. Both absolute (A) and relative (B) odorant concentrations are shown on a log-scale.

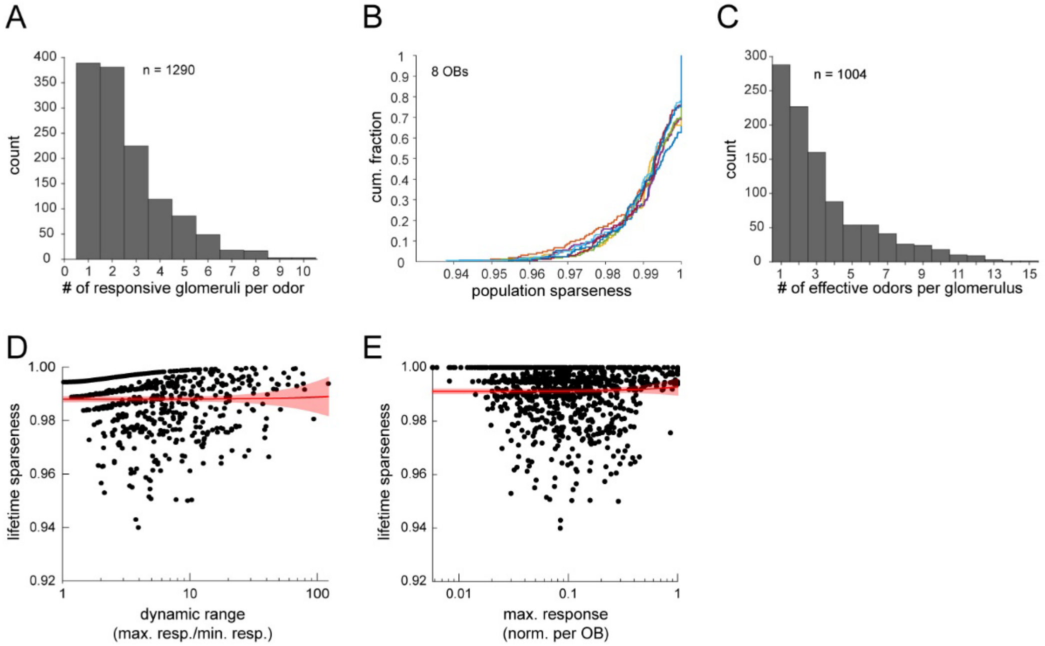


**Supplementary Figure S3. Sparse glomerular responses evoked across the 185-odorant panel.**

**A.** Histogram of number of glomeruli activated by a given odorant presentation, excluding non-responsive presentations (n = 1290, 8 OBs).
**B.** Cumulative distribution plots of population sparseness for each odorant, plotted separately for each of 8 OBs. Population sparseness of 1 indicates a single repsonsive glomerulus.
**C.** Histogram of number of number of effective odorants for each of 1004 glomeruli across the 8 OBs.
**D.** Lifetime sparseness of glomerular tuning across the 185-odorant panel, plotted as a function of dynamic range for all glomeruli responding to more than one odorant (n = 716 glomeruli). Dynamic range is conservatively defined as the ratio between the minimal non-zero odorant response and the maximal response. Red plot shows linear regression to data, with 95% confidence interval. Slope is not significantly different from 0 (linear regression t-test: p=0.89, $t_{714} = 0.14$) .
**E.** Lifetime sparseness as a function of maximal response amplitude for all imaged glomeruli (n = 1004). Responses normalized to the maximal response in a given OB. Red plot shows linear regression to data, as in (D). Slope is not significantly different from 0 (linear regression t-test: p=0.42, $t_{1002}=0.81$) .

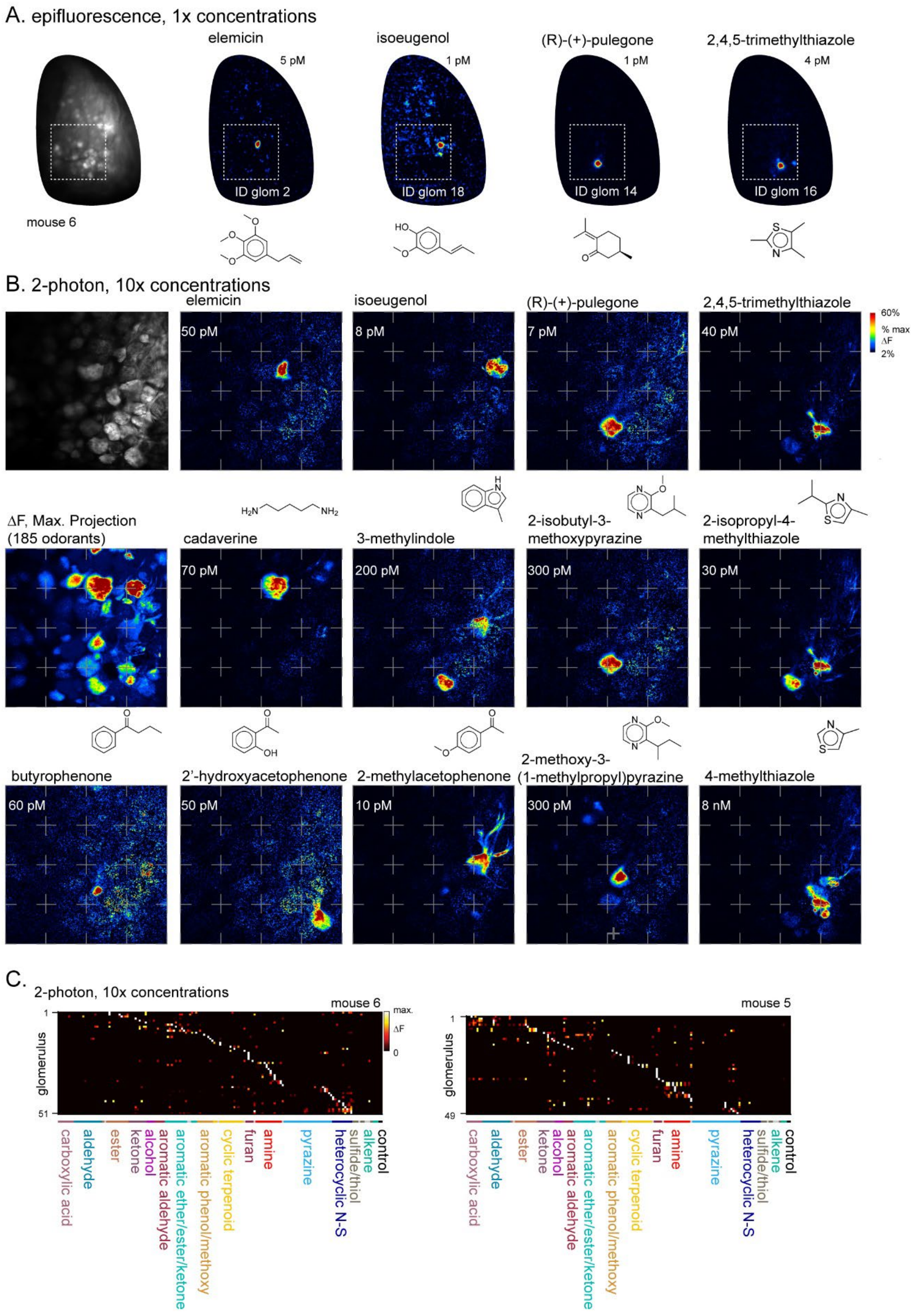

A. epifluorescence, 1x concentrations
elemicin
5 pM
ID glom 2
isoeugenol
1 pM
ID glom 18
(R)-(+)-pulegone
1 pM
ID glom 14
2,4,5-trimethylthiazole
4 pM
ID glom 16
mouse 6
B. 2-photon, 10x concentrations
elemicin
50 pM
isoeugenol
8 pM
(R)-(+)-pulegone
7 pM
2,4,5-trimethylthiazole
40 pM
60%
% max
ΔF
2%
ΔF, Max. Projection
(185 odorants)
cadaverine
70 pM
3-methylindole
200 pM
2-isobutyl-3-
methoxypyrazine
300 pM
2-isopropyl-4-
methylthiazole
30 pM
butyrophenone
60 pM
2'-hydroxyacetophenone
50 pM
2-methylacetophenone
10 pM
2-methoxy-3-
(1-methylpropyl)pyrazine
300 pM
4-methylthiazole
8 nM
C.
2-photon, 10x concentrations
mouse 6
mouse 5
glomerulus
max.
ΔF
0
1
51
49
carboxylic acid
aldehyde
ester
ketone
alcohol
aromatic aldehyde
aromatic ether/ester/ketone
aromatic phenol/methoxy
cyclic terpenoid
furan
amine
pyrazine
heterocyclic N-S
sulfide/thiol
alkene
control

**Figure S4. Two-photon imaging of glomerular odorant responses to ten-fold higher odorant concentrations.**

**A.** Epifluorescence response maps to '1x' concentrations of diagnostic odorants for functionally identified glomeruli, imaged across one OB (new mouse, ‘mouse #6’).
**B.** Two-photon imaging of inset area shown in (A) using '10x' odorant concentrations. Upper left, baseline fluorescence of glomerular imaging field. Response maps ($\Delta F$) show activation of specific glomeruli by select odorants. Dashed grid lines added to facilitate visual inspection across maps.
**C.** Left: Response matrix for glomeruli imaged in (B), showing responses to the '10x' concentration odorant panel for mouse #6. Only responses to odorants tested at 10x concentration (plus blank and solvent controls) are shown. Right: Response matrix for same odorant panel imaged in a second mouse (mouse #5).

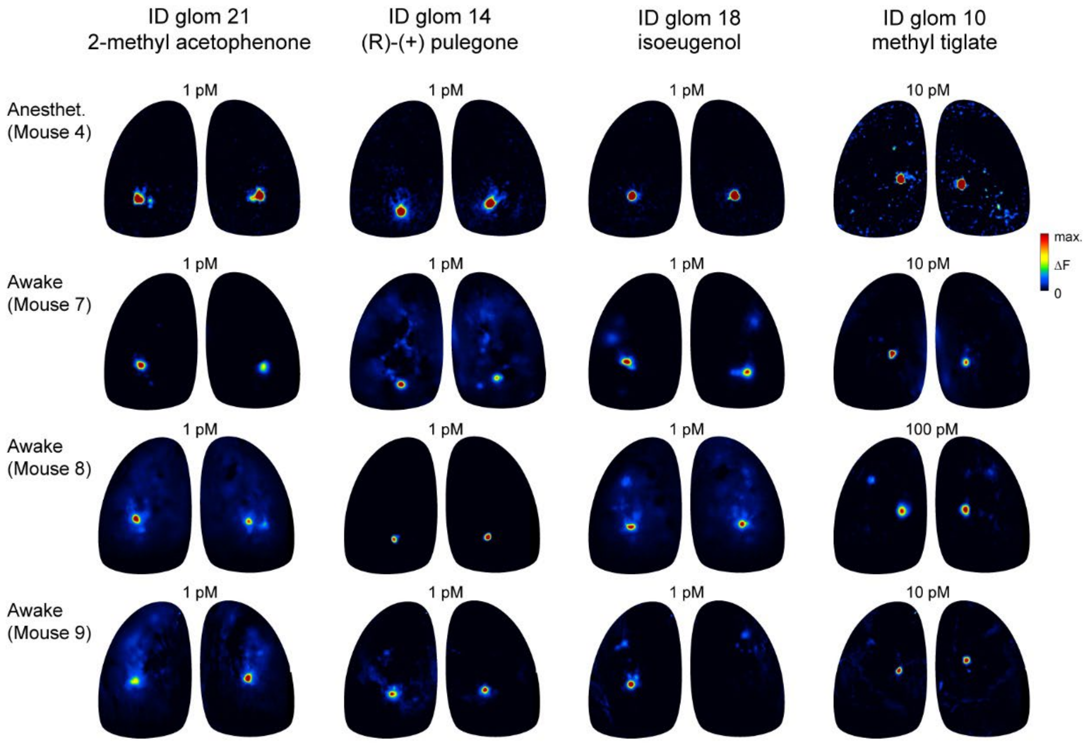


**Figure S5. Diagnostic odorants functionally identify glomeruli with similar sensitivity in awake mice.**

Comparison of response maps evoked by diagnostic odorants for four functionally-identified glomeruli in an anesthetized mouse (Mouse #4, top row) and in three awake, head-fixed mice (Mice #7-9). Odorants were prepared and presented identically. Awake response maps are averages of 3 - 5 presentations and are scaled as for anesthetized responses. Concentrations indicate estimated delivered concentration, as in previous Figures.

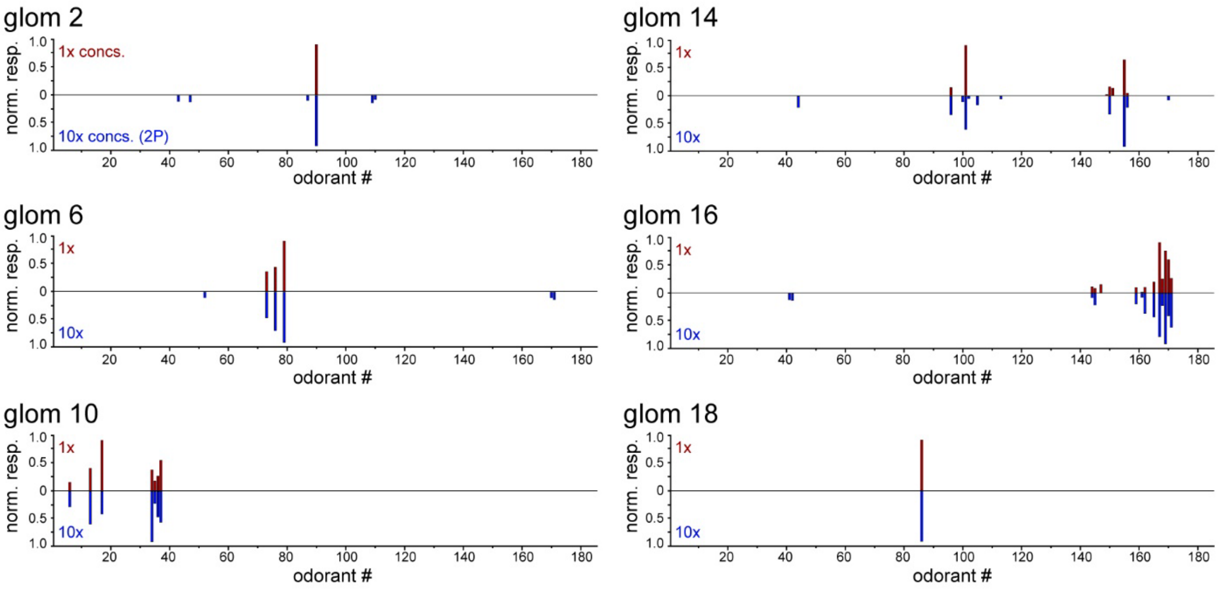


**Figure S6. Identified glomeruli maintain narrow tuning across a ten-fold increase in odorant concentrations.**

Response spectra of 6 functionally-identified glomeruli to the original 185-odorant panel ('1x',top, grey bars) and to the same odorants presented at ten-fold higher concentrations ('10x', bottom, blue bars), imaged in separate preparations with two-photon imaging. '1x' plots show median response spectrum across the 8 imaged OBs. '10x' plots show mean responses from two imaged OBs (two mice) (for glomeruli 2,14,18); or from a single OB (for glomeruli 6,10, 16), each normalized to the maximum response across the odorant panel.

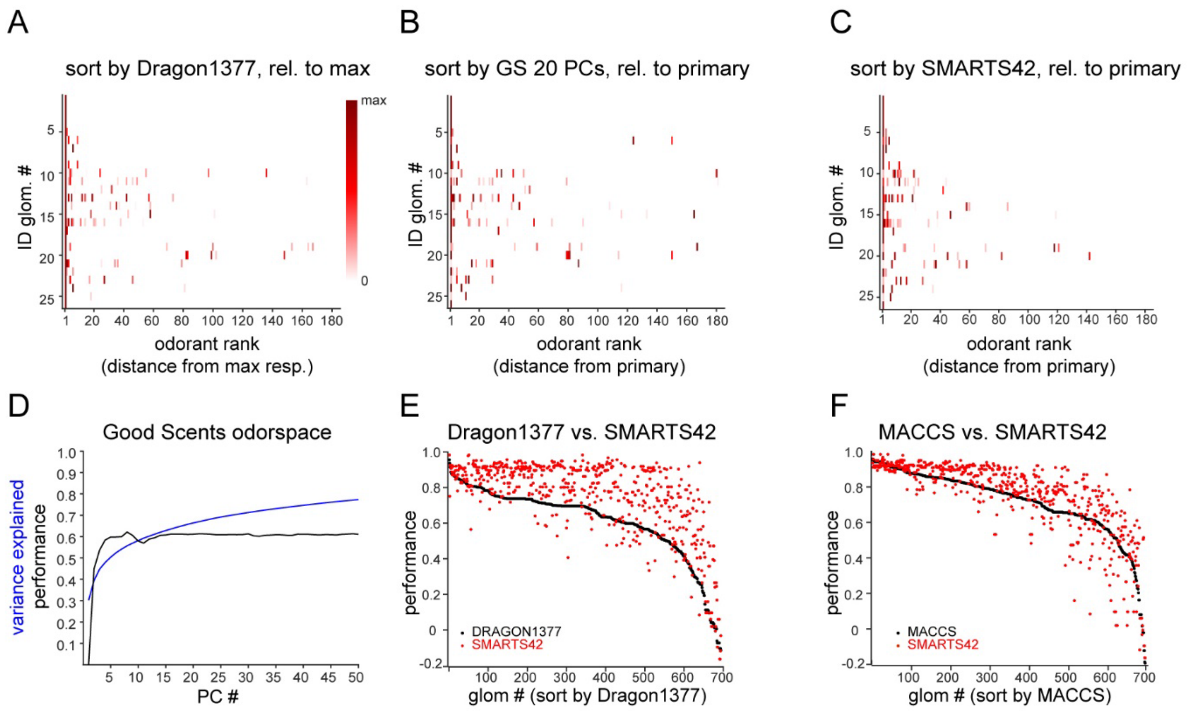


**Figure S7. Additional comparisons of glomerular response predictions by physicochemical descriptor sets.**

**A.** Median response spectra of functionally-identified glomeruli, with odorants ordered by physicochemical descriptor distance from the strongest-activating odorant for each glomerulus, using the Dragon1377 descriptor set (compare with ordering relative to 'primary' odorant, Fig. 3B).
**B.** Median response spectra ordered relative to distance in the space defined by the first 20 principal components of the space defined by the 2982-element Alvadesc descriptors applied to 2624 odorant compounds in the GoodScents odorant database (see Text).
**C.** Median response spectra ordered relative to distance (dice similarity) from the primary odorant using the SMARTS42 fingerprint.
**D.** Variance in the 2982-descriptor x 2624-odorant space explained, and mean prediction performance score for the 694 co-tuned glomeruli, as a function of principal component number. The first 20 PCs explain 66% of the variance. Performance asymptotes at ~0.6 after 5 – 10 PCs.
**E.** Comparison of prediction performance scores for each of the co-tuned glomeruli for the Dragon1377 descriptor set (black points) and the SMARTS42 set (red points), with glomeruli ordered by performance in the Dragon1377 descriptor space.
**F.** Comparison of MACCS (black points) and SMARTS42 performance scores (red points) for the same 694 co-tuned glomeruli.

| **Odorant** | **Code** | **CAS#** | **Epifl. est. conc. (M)** | **Dilution** | **Two-photon rel. conc.** | **Class** |
|---|---|---|---|---|---|---|
| propionic acid | C1 | 79-09-4 | 2E-11 | 1E-05 | 10× | acid |
| butyric acid | C2 | 107-92-6 | 2E-11 | 1E-05 | 10× | acid |
| 2-methylbutyric acid | C3 | 116-53-0 | 6E-12 | 1E-05 | 10× | acid |
| valeric acid | C4 | 109-52-4 | 6E-12 | 1E-05 | 10× | acid |
| isovaleric acid | C5 | 503-74-2 | 9E-12 | 1E-05 | 10× | acid |
| 2-methyl-2-pentenoic acid | C6 | 3142-72-1 | 4E-14 | 1E-06 | 10× | acid |
| hexanoic acid | C7 | 142-62-1 | 2E-11 | 1E-04 | 10× | acid |
| heptanoic acid | C8 | 111-14-8 | 9E-11 | 1E-03 | 10× | acid |
| *methacrolein* | *D1* | *78-85-3* | *1E-9* | *1E-05* | *10×* | *aldehyde* |
| butyraldehyde | D2 | 123-72-8 | 6E-09 | 1E-04 | 10× | aldehyde |
| isobutyraldehyde | D3 | 78-84-2 | 1E-09 | 1E-05 | 10× | aldehyde |
| 2-methylbutyraldehyde | D4 | 96-17-3 | 8E-11 | 1E-05 | 10× | aldehyde |
| trans-2-methyl-2-butenal | D5 | 497-03-0 | 1E-11 | 1E-06 | 10× | aldehyde |
| valeraldehyde | D6 | 110-62-3 | 2E-09 | 1E-04 | 10× | aldehyde |
| isovaleraldehyde | D7 | 590-86-3 | 4E-09 | 1E-04 | 1× | aldehyde |
| 2-methylvaleraldehyde | D8 | 123-15-9 | 1E-10 | 1E-05 | 10× | aldehyde |
| 2-methyl-2-pentenal | D9 | 623-36-9 | 6E-11 | 1E-05 | 1× | aldehyde |
| hexanal | D10 | 66-25-1 | 7E-10 | 1E-04 | 10× | aldehyde |
| heptanal | D11 | 111-71-7 | 3E-09 | 1E-03 | 10× | aldehyde |
| octanal | D12 | 124-13-0 | 1E-09 | 1E-03 | 10× | aldehyde |
| trans-2-nonenal | D13 | 18829-56-6 | 2E-09 | 1E-02 | 10× | aldehyde |
| trans-2,cis-6-nonadienal | D14 | 557-48-2 | 2E-09 | 1E-02 | 10× | aldehyde |
| trans-2-dodecenal | D15 | 20407-84-5 | 8E-10 | 1E-01 | 10× | aldehyde |
| 2-hexyl-2-decenal | D16 | 13893-39-5 | 5E-09 | 1E-01 | 10× | aldehyde |
| butyl acetate | E1 | 123-86-4 | 9E-10 | 1E-04 | 10× | ester |
| S-methyl thiobutanoate | E2.M1 | 2432-51-1 | 2E-10 | 1E-04 | 10× | mixed |
| isoamyl acetate | E3 | 123-92-2 | 4E-10 | 1E-04 | 10× | ester |
| hexyl acetate | E4 | 142-92-7 | 1E-09 | 1E-03 | 10× | ester |
| ethyl butyrate | E5 | 105-54-4 | 1E-09 | 1E-04 | 10× | ester |
| methyl 2-methylbutyrate | E6 | 868-57-5 | 2E-10 | 1E-05 | 10× | ester |
| vinyl butyrate | E7 | 123-20-6 | 9E-11 | 1E-05 | 10× | ester |
| methyl valerate | E8 | 624-24-8 | 8E-10 | 1E-04 | 10× | ester |
| 1-octen-3-yl butyrate | E9 | 16491-54-6 | 5E-09 | 1E-01 | 10× | ester |
| methyl tiglate | E10 | 6622-76-0 | 1E-11 | 1E-06 | 10× | ester |
| ethyl tiglate | E11 | 5837-78-5 | 2E-12 | 1E-06 | 10× | ester |
| isopropyl tiglate | E12 | 1733-25-1 | 2E-11 | 1E-05 | 10× | ester |
| hexyl tiglate | E13 | 16930-96-4 | 4E-10 | 1E-02 | 10× | ester |
| diacetyl | K1 | 431-03-8 | 4E-09 | 1E-04 | 10× | ketone |
| *2-butanone* | *K2* | *78-93-3* | *7E-10* | *1E-05* | *10×* | *ketone* |
| *2-pentanone* | *K3* | *107-87-9* | *3E-09* | *1E-04* | *10×* | *ketone* |
| 4-methyl-3-penten-2-one | K4 | 141-79-7 | 9.E-10 | 1E-04 | 10× | ketone |

| | | | | | | |
|---|---|---|---|---|---|---|
| 2-hexanone | K5 | 591-78-6 | 1E-09 | 1E-04 | 10× | ketone |
| 3-hepten-2-one | K6 | 1119-44-4 | 2E-09 | 1E-03 | 10× | ketone |
| 5-methyl-2-hepten-4-one | K7 | 81925-81-7 | 1E-09 | 1E-03 | 10× | ketone |
| 2-octanone | K8 | 111-13-7 | 1E-10 | 1E-04 | 10× | ketone |
| 3-octen-2-one | K9 | 1669-44-9 | 2E-10 | 1E-04 | 1× | ketone |
| 2-nonanone | K10 | 821-55-6 | 5E-10 | 1E-03 | 10× | ketone |
| 1-hexanol | L1 | 111-27-3 | 7E-10 | 1E-03 | 10× | alcohol |
| cis-3-hexenol | L2 | 928-96-1 | 8E-10 | 1E-03 | 10× | alcohol |
| geraniol | L3 | 106-24-1 | 1E-09 | 1E-02 | 10× | alcohol |
| R(-)1-octen-3-ol | L4 | 3687-48-7 | 4E-10 | 1E-03 | 10× | alcohol |
| benzaldehyde | AD1 | 100-52-7 | 8E-11 | 1E-04 | 10× | aromatic ald. |
| trans-cinnamaldehyde | AD2 | 14371-10-9 | 2E-11 | 1E-03 | 10× | aromatic ald. |
| cuminaldehyde | AD3 | 122-03-2 | 5E-11 | 1E-03 | 10× | aromatic ald. |
| p-anisaldehyde | AD4 | 123-11-5 | 8E-12 | 1E-05 | 10× | aromatic ald. |
| piperonal | AD5 | 120-57-0 | 2E-12 | 1E-03 | 1× | aromatic ald. |
| vanillin | AD6 | 121-33-5 | 3E-11 | 1E-01 | 10× | aromatic ald. |
| acetophenone | AK1 | 98-86-2 | 3E-12 | 1E-05 | 10× | aromatic ket./est. |
| 2'-hydroxyacetophenone | AK2 | 118-93-4 | 5E-12 | 1E-04 | 1× | aromatic ket./est. |
| 2,4-dimethylacetophenone | AK3 | 89-74-7 | 6E-13 | 1E-05 | 10× | aromatic ket./est. |
| 4-aminoacetophenone | AK4.M2 | 99-92-3 | 3E-10 | 1E+00 | 1× | mixed |
| 4-methylacetophenone | AK5 | 122-00-9 | 7E-12 | 1E-04 | 10× | aromaticket./est. |
| 4-methoxyacetophenone | AK6 | 100-06-1 | 5E-13 | 1E-04 | 1× | aromatic ket./est. |
| 2-methylacetophenone | AK7 | 577-16-2 | 1E-12 | 1E-05 | 10× | aromatic ket./est. |
| propiophenone | AK8 | 93-55-0 | 1E-11 | 1E-04 | 1× | aromatic ket./est. |
| butyrophenone | AK9 | 495-40-9 | 6E-12 | 1E-04 | 10× | aromatic ket./est. |
| 4-phenyl-2-butanone | AK10 | 2550-26-7 | 5E-11 | 1E-03 | 10× | aromatic ket./est. |
| 4-(4-hydroxyphenyl)-2-butanone | AK11 | 5471-51-2 | 8E-11 | 1E-01 | 10× | aromatic ket./est. |
| methyl salicylate | AE1 | 119-36-8 | 4E-11 | 1E-03 | 1× | aromatic ket./est. |
| methyl benzoate | AE2 | 93-58-3 | 3E-11 | 1E-04 | 10× | aromatic ket./est. |
| benzyl acetate | AE3 | 140-11-4 | 1E-10 | 1E-03 | 10× | aromatic ket./est. |
| ethyl benzoate | AE4 | 93-89-0 | 2E-11 | 1E-04 | 10× | aromatic ket./est. |
| benzyl benzoate | AE5 | 120-51-4 | 4E-11 | 1E-01 | 10× | aromatic ket./est. |
| methyl anthranilate | AE6.M3 | 134-20-3 | 1E-13 | 1E-05 | 10× | mixed |
| dimethyl anthranilate | AE7.M4 | 85-91-6 | 2E-11 | 1E-03 | 10× | mixed |
| phenylacetate | AE8 | 122-79-2 | 3E-12 | 1E-05 | 10× | aromatic ket./est. |
| ethyl phenylacetate | AE9 | 101-97-3 | 7E-11 | 1E-03 | 1× | aromatic ket./est. |
| allyl phenylacetate | AE10 | 1797-74-6 | 2E-11 | 1E-03 | 10× | aromatic ket./est. |
| phenyl propionate | AE11 | 637-27-4 | 1E-11 | 1E-04 | 10× | aromatic ket./est. |
| m-cresol | AP1 | 108-39-4 | 1E-10 | 1E-03 | 1× | aromatic phen./moxy. |
| carvacrol | AP2 | 499-75-2 | 2E-09 | 1E-01 | 10× | aromatic phen./moxy. |
| 4-methylanisole | AP3 | 104-93-8 | 9E-11 | 1E-04 | 10× | aromatic phen./moxy. |
| guaiacol | AP4 | 90-05-1 | 9E-13 | 1E-05 | 1× | aromatic phen./moxy. |
| 2,6-dimethoxyphenol | AP5 | 91-10-1 | 2E-12 | 1E-03 | 1× | aromatic phen./moxy. |

| | | | | | | |
|---|---|---|---|---|---|---|
| eugenol | AP6 | 97-53-0 | 7E-13 | 1E-04 | 10× | aromatic phen./moxy. |
| isoeugenol | AP7 | 97-54-1 | 8E-13 | 1E-04 | 10× | aromatic phen./moxy. |
| methyl eugenol | AP8 | 93-15-2 | 2E-12 | 1E-04 | 10× | aromatic phen./moxy. |
| methyl isoeugenol | AP9 | 93-16-3 | 8E-13 | 1E-04 | 10× | aromatic phen./moxy. |
| eugenyl acetate | AP10 | 93-28-7 | 4E-11 | 1E-02 | 10× | aromatic phen./moxy. |
| elemicin | AP11 | 487-11-6 | 5E-12 | 1E-03 | 10× | aromatic phen./moxy. |
| fenchol | CT1 | 1632-73-1 | 5E-10 | 1E-02 | 1× | cyclic terpenoid |
| 2-ethyl fenchol | CT2 | 18368-91-7 | 6E-11 | 1E-02 | 10× | cyclic terpenoid |
| alpha-pinene | CT3 | 7785-70-8 | 3E-09 | 1E-03 | 10× | cyclic terpenoid |
| isobornyl isovalerate | CT4.M5 | 7779-73-9 | 1E-09 | 1E-01 | 10× | mixed |
| 1,8-cineole | CT5 | 470-82-6 | 1E-09 | 1E-03 | 10× | cyclic terpenoid |
| L-carvone | CT6 | 6485-40-1 | 1E-10 | 1E-03 | 10× | cyclic terpenoid |
| beta-ionone | CT7 | 14901-07-6 | 1E-11 | 1E-03 | 10× | cyclic terpenoid |
| beta-damascone | CT8 | 23726-91-2 | 1E-11 | 1E-03 | 10× | cyclic terpenoid |
| damascenone | CT9 | 23696-85-7 | 2E-10 | 1E-02 | 10× | cyclic terpenoid |
| menthone | CT10 | 10458-14-7 | 3E-10 | 1E-03 | 10× | cyclic terpenoid |
| (R)-(+)-pulegone | CT11 | 89-82-7 | 7E-13 | 1E-05 | 10× | cyclic terpenoid |
| (+)-isomenthone | CT12 | 1196-31-2 | 2E-10 | 1E-03 | 10× | cyclic terpenoid |
| nootkatone | CT13 | 4674-50-4 | 9E-10 | 1E+00 | 1× | cyclic terpenoid |
| (+-)-menthol | CT14 | 89-78-1 | 6E-11 | 1E-02 | 10× | cyclic terpenoid |
| (+)-neomenthol | CT15 | 2216-52-6 | 2E-10 | 1E-02 | 10× | cyclic terpenoid |
| (+-)-geosmin | CT16 | 16423-19-1 | 4E-11 | 1E-01 | 10× | cyclic terpenoid |
| (-)-ambroxide | CT17 | 6790-58-5 | 7E-10 | 1E-01 | 10× | cyclic terpenoid |
| (+)-menthofuran | CT18.M6 | 17957-94-7 | 2E-10 | 1E-03 | 10× | mixed |
| 2-pentylfuran | F1 | 3777-69-3 | 2E-09 | 1E-03 | 10× | furan/pyrone/lactone |
| 5-ethyl-4-hydroxy-2-methyl-3(2H)-furanone | F2 | 27538-09-6 | 9E-13 | 1E-04 | 10× | furan/pyrone/lactone |
| 5-methylfurfural | F3 | 620-02-0 | 5E-10 | 1E-03 | 10× | furan/pyrone/lactone |
| ethyl maltol | F4 | 4940-11-8 | 1E-12 | 1E-02 | 1× | furan/pyrone/lactone |
| gamma-undecalactone | F5 | 104-67-6 | 3E-09 | 1E+00 | 1× | furan/pyrone/lactone |
| coumarin | F6 | 91-64-5 | 7E-10 | 1E-02 | 10× | furan/pyrone/lactone |
| 1-furfurylpyrrole | F7.M7 | 1438-94-4 | 1E-10 | 1E-02 | 10× | mixed |
| ethylenediamine | N1 | 107-15-3 | 1E-10 | 1E-05 | 10× | amine |
| butylamine | N2 | 109-73-9 | 6E-10 | 1E-05 | 10× | amine |
| 2-methylbutylamine | N3 | 96-15-1 | 4E-10 | 1E-05 | 10× | amine |
| N-butyldimethylamine | N4 | 927-62-8 | 3E-10 | 1E-05 | 10× | amine |
| isopentylamine | N5 | 107-85-7 | 3E-11 | 1E-06 | 10× | amine |
| cadaverine | N6 | 462-94-2 | 7E-13 | 1E-06 | 10× | amine |
| octylamine | N7 | 111-86-4 | 7E-11 | 1E-04 | 10× | amine |
| N,N-dimethyloctylamine | N8 | 7378-99-6 | 6E-12 | 1E-05 | 10× | amine |
| cyclohexylamine | N9 | 108-91-8 | 6E-11 | 1E-05 | 10× | amine |
| N,N-dimethylcyclohexylamine | N10 | 98-94-2 | 2E-11 | 1E-05 | 10× | amine |
| phenethylamine | N11 | 64-04-0 | 3E-13 | 1E-06 | 1× | amine |
| benzylamine | N12 | 100-46-9 | 5E-11 | 1E-04 | 10× | amine |

| *tyramine* | *N13* | *51-67-2* | *2E-09* | *1E+00* | 1× | *amine* |
|---|---|---|---|---|---|---|
| N,N-dimethyl-2-phenethylamine | N14 | 1126-71-2 | 2E-13 | 1E-06 | 10× | amine |
| 3-phenylpropylamine | N15 | 2038-57-5 | 1E-11 | 1E-04 | 10× | amine |
| N-methyl piperidine | N16 | 626-67-5 | 2E-10 | 1E-05 | 1× | amine |
| 5-methyl heptan-3-one oxime | N17.M8 | 22457-23-4 | 3E-10 | 1E-02 | 10× | mixed |
| pyrazine | P1 | 290-37-9 | 2E-09 | 1E-04 | 10× | pyrazine |
| 2-acetylpyrazine | P2 | 22047-25-2 | 1E-09 | 1E-02 | 10× | pyrazine |
| 2-methylpyrazine | P3 | 109-08-0 | 7E-11 | 1E-05 | 10× | pyrazine |
| 2-methoxypyrazine | P4 | 3149-28-8 | 3E-11 | 1E-05 | 10× | pyrazine |
| 2-ethylpyrazine | P5 | 13925-00-3 | 3E-11 | 1E-05 | 10× | pyrazine |
| 2,3-diethylpyrazine | P6 | 15707-24-1 | 6E-10 | 1E-03 | 1× | pyrazine |
| 2-chloropyrazine | P7 | 14508-49-7 | 2E-09 | 1E-03 | 10× | pyrazine |
| pyrazineethanethiol | P8 | 35250-53-4 | 4E-11 | 1E-03 | 10× | pyrazine |
| 2-methoxy-3-methylpyrazine | P9 | 2847-30-5 | 5E-11 | 1E-04 | 10× | pyrazine |
| 2-ethyl-3-methylpyrazine | P10 | 15707-23-0 | 5E-11 | 1E-04 | 10× | pyrazine |
| 2,3-dimethylpyrazine | P11 | 5910-89-4 | 3E-10 | 1E-04 | 10× | pyrazine |
| 2,5-dimethylpyrazine | P12 | 123-32-0 | 3E-10 | 1E-04 | 10× | pyrazine |
| 2,6-dimethylpyrazine | P13 | 108-50-9 | 3E-09 | 1E-03 | 10× | pyrazine |
| 2,3,5-trimethylpyrazine | P14 | 14667-55-1 | 1E-11 | 1E-05 | 10× | pyrazine |
| 2,3,5,6-tetramethylpyrazine | P15 | 1124-11-4 | 5E-10 | 1E-03 | 10× | pyrazine |
| 2-ethyl-5-methylpyrazine | P16 | 13360-64-0 | 2E-11 | 1E-05 | 10× | pyrazine |
| 2-acetyl-3,(5 or 6)-dimethylpyrazine | P17 | 54300-08-2 | 2E-09 | 1E-03 | 1× | pyrazine |
| 2-isobutyl-3-methylpyrazine | P18 | 13925-06-9 | 2E-10 | 1E-03 | 10× | pyrazine |
| 2-acetyl-3-ethylpyrazine | P19 | 32974-92-8 | 2E-10 | 1E-02 | 1× | pyrazine |
| 2-acetyl-3-methylpyrazine | P20 | 23787-80-6 | 8E-11 | 1E-03 | 10× | pyrazine |
| 2-ethyl-3-methoxypyrazine | P21 | 25680-58-4 | 1E-11 | 1E-05 | 10× | pyrazine |
| 2-methoxy-3(5 or 6)-isopropylpyrazine | P22 | 93905-03-4 | 3E-11 | 1E-04 | 10× | pyrazine |
| 2-isobutyl-3-methoxypyrazine | P23 | 24683-00-9 | 3E-11 | 1E-03 | 10× | pyrazine |
| 2-methoxy-3-(1-methylpropyl)pyrazine | P24 | 24168-70-5 | 3E-11 | 1E-03 | 10× | pyrazine |
| 2-isopropyl-3-methoxypyrazine | P25 | 25773-40-4 | 9E-12 | 1E-04 | 10× | pyrazine |
| 5H-5-methyl-6,7-dihydrocyclopenta[b]pyrazine | P26 | 23747-48-0 | 2E-10 | 1E-03 | 1× | pyrazine |
| 5,6,7,8-tetrahydroquinoxaline | P27 | 34413-35-9 | 4E-11 | 1E-03 | 10× | pyrazine |
| 5-methylquinoxaline | P28 | 13708-12-8 | 4E-10 | 1E-02 | 10× | pyrazine |
| 2-acetylpyridine | NS1 | 1122-62-9 | 4E-11 | 1E-04 | 1× | heterocyclic N-S |
| 4-tert-butylpyridine | NS2 | 3978-81-2 | 4E-12 | 1E-05 | 10× | heterocyclic N-S |
| indole | NS3 | 120-72-9 | 9E-11 | 1E-02 | 10× | heterocyclic N-S |
| 3-methylindole | NS4 | 83-34-1 | 2E-11 | 1E-02 | 10× | heterocyclic N-S |
| 2-isobutylthiazole | NS5 | 18640-74-9 | 3E-12 | 1E-05 | 10× | heterocyclic N-S |
| 2-acetylthiazole | NS6 | 24295-03-2 | 1E-10 | 1E-03 | 10× | heterocyclic N-S |
| 2-isopropyl-4-methylthiazole | NS7 | 15679-13-7 | 3E-12 | 1E-05 | 10× | heterocyclic N-S |
| 2-methyl-2-thiazoline | NS8 | 2346-00-1 | 4E-10 | 1E-04 | 10× | heterocyclic N-S |
| 2,4,5-trimethylthiazole | NS9 | 13623-11-5 | 5E-12 | 1E-05 | 10× | heterocyclic N-S |

| | | | | | | |
|---|---|---|---|---|---|---|
| ethyl-2,5-dihydro-4-methylthiazole | NS10 | 41803-21-8 | 6E-11 | 1E-04 | 10× | heterocyclic N-S |
| 4-methylthiazole | NS11 | 693-95-8 | 8E-10 | 1E-04 | 10× | heterocyclic N-S |
| 2-methyl-4-propyl-1,3-oxathiane | NS12.M9 | 67715-80-4 | 9E-11 | 1E-03 | 1× | mixed |
| dimethyl trisulfide | S1 | 3658-80-8 | 8E-10 | 1E-03 | 10× | sulfide-thiol |
| *allyl disulfide* | *S2* | *2179-57-9* | *7E-10* | *1E-03* | *10×* | *sulfide-thiol* |
| 4-methoxy-2-methyl-2-butanethiol | S3 | 94087-83-9 | 3E-12 | 1E-06 | 10× | sulfide-thiol |
| methional | S4.M10 | 3268-49-3 | 1E-11 | 1E-05 | 10× | mixed |
| 3-mercaptohexyl acetate | S5 | 136954-20-6 | 4E-12 | 1E-04 | 10× | sulfide-thiol |
| 3-(methylthio)-1-hexanol | S6 | 51755-66-9 | 6E-09 | 1E-02 | 10× | sulfide-thiol |
| furfuryl mercaptan | S7.M11 | 98-02-2 | 3E-12 | 1E-06 | 10× | mixed |
| difurfuryl disulfide | S8.M12 | 4437-20-1 | 4E-13 | 1E-03 | 10× | mixed |
| furfuryl methyl sulfide | S9.M13 | 1438-91-1 | 1E-09 | 1E-03 | 10× | mixed |
| 2-methyl-3-tetrahydrofuranthiol | S10 | 57124-87-5 | 2E-10 | 1E-04 | 10× | sulfide-thiol |
| myrcene | ENE1 | 123-35-3 | 2E-09 | 1E-03 | 10× | alkene |
| 1,3,5-undecatriene | ENE2 | 16356-11-9 | 5E-09 | 1E-02 | 10× | alkene |
| (R)-(+)-limonene | ENE3 | 5989-27-5 | 1E-09 | 1E-03 | 10× | alkene |
| empty | - | - | - | - | - | control |
| triglyceride | - | - | - | - | - | control |

**Table S1. Odorants and estimated concentrations used in the study.**

**Code:** abbreviated identifier code, used in Fig. 4C and Document S1. **Epifl. est. conc.:** estimated delivered concentration of odorant vapor, in mols/L (M), for '1x' dataset. Most commonly-presented values (of 4 preparations) are shown, reported to one significant digit precision. **Dilution:** liquid dilution of odorant used to generate delivered concentration. **Two-photon rel. conc.:** concentration relative to '1x' dataset used for the two-photon imaging dataset. **Class:** nominal classification based on structural features, used in Fig. 5 and Fig. 6. Odorants in italics gave no response at the given concentration in any of the 8 OBs.

| # | Odorant | Est. conc. (M) | Error ratio | Median ORS corr. | Mediolateral (µm) | Anteroposterior (µm) |
|---|---|---|---|---|---|---|
| 1 | benzaldehyde | 8E-11 | 0.00 | 1.00 | 1448.3 ± 76.9 | 1298.4 ± 82.4 |
| 2 | elemicin | 5E-12 | 0.00 | 1.00 | 1011.6 ± 80.9 | 753.2 ± 126.7 |
| 3 | vanillin | 3E-11 | 0.07 | 1.00 | 1249.6 ± 54.8 | 1651 ± 75.9 |
| 4 | trans-2-dodecenal | 8E-10 | 0.00 | 1.00 | 514.9 ± 54.2 | 2117.5 ± 179.5 |
| 5 | ethyl phenylacetate | 7E-11 | 0.00 | 1.00 | 884 ± 50.5 | 1911.9 ± 118.9 |
| | allyl phenylacetate | 1E-10 | 0.00 | 1.00 | | |
| 6 | phenylacetate | 3E-12 | 0.00 | 0.99 | 1375.6 ± 93.3 | 1103.4 ± 111.8 |
| | phenyl propionate | 1E-11 | 0.00 | 0.99 | | |
| 7 | heptanoic acid | 7E-11 | 0.00 | 0.97 | 1026.6 ± 60.5 | 2132.8 ± 54.7 |
| | heptanal | 2E-9 | 0.00 | 0.95 | | |
| 8 | methional | 1E-11 | 0.00 | 1.00 | 1680.2 ± 99.6 | 1144.1 ± 91.3 |
| 9 | 3-mercaptohexyl acetate | 4E-12 | 0.00 | 0.97 | 1316.2 ± 170.4 | 1244.5 ± 67.3 |
| 10 | trans-2-methyl-2-butenal | 1E-11 | 0.00 | 0.95 | 700.8 ± 79.3 | 1103.7 ± 113.9 |
| | 2-methyl-2-pentenal | 6E-11 | 0.00 | 0.95 | | |
| | methyl tiglate | 1E-11 | 0.00 | 0.95 | | |
| | ethyl tiglate | 2E-12 | 0.00 | 0.95 | | |
| | isopropyl tiglate | 2E-11 | 0.00 | 0.95 | | |
| | hexyl tiglate | 4E-10 | 0.00 | 0.95 | | |
| 11 | isovaleric acid | 9E-12 | 0.00 | 0.92 | 973.5 ± 32.4 | 1696.9 ± 138 |
| | isovaleraldehyde | 4E-9 | 0.00 | 0.92 | | |
| 12 | 2'-hydroxyacetophenone | 5E-12 | 0.00 | 0.86 | 1258.5 ± 66.8 | 444 ± 68.9 |
| 13 | pyrazine | 2E-9 | 0.03 | 0.91 | 1618 ± 54.2 | 1154.8 ± 117.4 |
| 14 | 2-isobutyl-3-methoxypyrazine | 3E-11 | 0.04 | 0.91 | 1007.4 ± 93.9 | 503.6 ± 80.8 |
| | (R)-(+)-pulegone | 7E-13 | 0.04 | 0.91 | | |
| 15 | 4-(4-hydroxyphenyl)-2-butanone | 4E-10 | 0.02 | 0.84 | 1202.5 ± 92.4 | 1771.8 ± 66.1 |
| 16 | 2,4,5-trimethylthiazole | 5E-12 | 0.04 | 0.91 | 1242.6 ± 128.3 | 460.8 ± 106.6 |
| | ethyl-2,5-dihydro-4-methylthiazole | 3E-10 | 0.04 | 0.91 | | |
| 17 | 4-methoxy-2-methyl-2-butanethiol | 3E-12 | 0.07 | 0.98 | 1633.1 ± 116.8 | 853.6 ± 107.1 |
| | 2-methyl-3-tetrahydrofuranthiol | 2E-10 | 0.07 | 0.98 | | |
| 18 | isoeugenol | 8E-13 | 0.05 | 1.00 | 1162.9 ± 86.7 | 746.2 ± 105.4 |
| 19 | menthone | 3E-10 | 0.05 | 0.85 | 1058.4 ± 76.9 | 701.8 ± 93.8 |
| 20 | 2-hexanone | 1E-9 | 0.11 | 0.94 | 1372 ± 98.3 | 945.6 ± 105.8 |
| 21 | acetophenone | 1E-11 | 0.02 | 0.83 | 1317.2 ± 81.1 | 776.9 ± 61.6 |
| | 2-methylacetophenone | 1E-12 | 0.02 | 0.83 | | |
| 22 | methyl eugenol | 2E-12 | 0.13 | 0.90 | 1491.6 ± 107 | 771.2 ± 103.5 |
| 23 | 2-methylbutyraldehyde | 8E-11 | 0.16 | 0.87 | 928.6 ± 38.7 | 1497.3 ± 126.9 |
| | 2-methylvaleraldehyde | 1E-10 | 0.16 | 0.87 | | |
| | methyl 2-methylbutyrate | 1E-10 | 0.16 | 0.87 | | |
| 24 | hexanal | 7E-10 | 0.18 | 0.97 | 1047.7 ± 63.7 | 1977.8 ± 66 |
| 25 | fenchol | 5E-10 | 0.16 | 0.98 | 1251 ± 109.5 | 795.9 ± 76.9 |
| 26 | 5-methylfurfural | 5E-10 | 0.19 | 1.00 | 1465 ± 67.1 | 1153.3 ± 380.5 |

**Table S2. Diagnostic odorants and concentrations for functionally-identified glomeruli.**

**Error ratio:** Incidence of mismatch between strongest-activated glomeruli and glomeruli with most correlated odorant response spectra (ORS) across 2 OBs, divided by all potential 2-OB comparisons. **Median ORS corr.:** Median ORS correlation coefficient (Pearson's r) across all pairwise comparisons of response spectra for the maximally-activated glomerulus in each responsive OB.

| # | Odorant | Est. conc. (M) | Error ratio | Median ORS corr. |
|---|---|---|---|---|
| - | 2,3,5-trimethylpyrazine | 1E-11 | 0.12 | 0.74 |
| - | p-anisaldehyde | 8E-12 | 0.21 | 0.93 |
| - | piperonal | 2E-12 | 0.21 | 0.93 |
| - | cadaverine | 2E-12 | 0.23 | 0.98 |
| - | furfuryl mercaptan | 3E-12 | 0.24 | 0.92 |
| - | difurfuryl disulfide | 4E-13 | 0.24 | 0.92 |
| - | 2-methyl-2-pentenoic acid | 4E-14 | 0.25 | 0.93 |
| - | 2-methylpyrazine | 7E-11 | 0.26 | 0.85 |
| - | 2-chloropyrazine | 2E-9 | 0.26 | 0.85 |
| - | damascenone | 2E-10 | 0.29 | 0.76 |
| - | N,N-dimethyloctylamine | 3E-11 | 0.32 | 1.00 |
| - | beta-damascone | 1E-11 | 0.32 | 0.81 |
| - | 2-ethyl-5-methylpyrazine | 1E-11 | 0.36 | 0.74 |
| - | 3-(methylthio)-1-hexanol | 6E-9 | 0.39 | 0.89 |
| - | benzyl benzoate | 4E-11 | 0.39 | 0.82 |
| - | 4-methylthiazole | 8E-10 | 0.39 | 0.87 |
| - | eugenol | 7E-13 | 0.40 | 0.59 |
| - | 2-methylbutyric acid | 5E-12 | 0.43 | 0.95 |
| - | beta-ionone | 7E-11 | 0.43 | 0.73 |
| - | L-carvone | 1E-10 | 0.43 | 0.66 |
| - | 2-methoxy-3-methylpyrazine | 5E-11 | 0.45 | 0.88 |
| - | 4-methylacetophenone | 7E-12 | 0.48 | 0.81 |
| - | N,N-dimethyl-2-phenethylamine | 5E-13 | 0.50 | 0.75 |
| - | N-methyl piperidine | 2E-10 | 0.50 | 0.99 |
| - | 1,3,5-undecatriene | 5E-9 | 0.50 | 0.55 |
| - | 2-methyl-2-thiazoline | 1E-9 | 0.52 | 0.83 |
| - | 4-methoxyacetophenone | 5E-13 | 0.54 | 0.83 |
| - | 2,6-dimethoxyphenol | 2E-12 | 0.54 | 1.00 |
| - | isopentylamine | 3E-11 | 0.55 | 0.67 |
| - | nootkatone | 9E-10 | 0.57 | 0.97 |
| - | champignol | 4E-10 | 0.59 | 0.50 |
| - | 2-acetyl-3,(5 or 6)-dimethylpyrazine | 2E-9 | 0.59 | 0.42 |
| - | (+)-menthofuran | 2E-10 | 0.59 | 0.61 |
| - | 2-ethyl-3-methoxypyrazine | 4E-11 | 0.61 | 0.51 |
| - | 2-octanone | 1E-10 | 0.66 | 0.62 |
| - | furfuryl methyl sulfide | 1E-9 | 0.67 | 0.20 |
| - | geraniol | 1E-9 | 0.67 | 0.29 |
| - | butyrophenone | 2E-11 | 0.70 | 0.50 |
| - | 4-methylanisole | 9E-11 | 0.75 | 0.00 |
| - | benzyl acetate | 1E-10 | 0.86 | 0.15 |

**Table S3. Additional odorants and concentrations eliciting consistently sparse activation but failing conservative requirements for functional identification.**

| # | SMARTS pattern | description |
|---|---|---|
| 1 | *-C(=O)-[OH1] | carboxylic acid |
| 2 | [CH1]=O | aldehyde |
| 3 | C-C(=O)-[O]-C | ester |
| 4 | C-C(=O)-[S]-C | thioester |
| 5 | [!O&!S]-C(=O)-[!O&!S] | ketone |
| 6 | [OX2H][CX4&!$(C([OX2H])[O,S,#7,#15]),c] | alcohol |
| 7 | c1ccccc1 | benzyl |
| 8 | C~C(~C)~C~C~C~C(~C)~C | monoterpene |
| 9 | [#8]1~[#6]~[#6]~[#6]~[#6]1 | furanoid |
| 10 | o1cccc1 | furan |
| 11 | [NH2][C] | primary amine |
| 12 | [NH](C)C | secondary amine |
| 13 | [NH0](C)(C)C | tertiary amine |
| 14 | [N,n]1~[C,c]~[C,c]~[C,c]~[C,c]~[C,c]1 | pyridine |
| 15 | [n,N]1~[C,c]~[C,c]~[C,c]~[C,c]1 | pyrrole |
| 16 | [N,n]1~[C,c]~[C,c]~[N,n]~[C,c]~[C,c]1 | pyrazine |
| 17 | [#16]1~[#6]~[#7]~[#6]~[#6]1 | thiazoline |
| 18 | [!#8]~C-S-C~[!#8] | thioether |
| 19 | [$(C-S-S-C),$(C-S-S-S-C)] | sulfide |
| 20 | [#6]-[SH] | thiol |
| 21 | [#6]=[#6] | alkene |
| 22 | [#16] | sulfur |
| 23 | [#7] | nitrogen |
| 24 | [#8] | oxygen |
| 25 | [R] | ring |
| 26 | [CH3]-*-[CH2]-* | 4-bond chain with C at 1 and 3 |
| 27 | *!@*@*!@* | ortho-substituted rings |
| 28 | *!@*@*@*!@* | meta-substituted rings |
| 29 | *1(!@*)@*@*@*(!@*)@*@*@1 | para substituted 6-ring but not fused ring |
| 30 | C~C(~C)~[R1]1~[R1]~[R1]~[R1](~C)~[R1]~[R1]~1 | menthane scaffold |
| 31 | C~C(~C)~2~[R2]1~[R2]~2~[R1]~[R1](~C)~[R1]~[R1]~1 | carene scaffold |
| 32 | C~C(~C)~[R2]12~[R1]~[R2]~2~[R1](~C)~[R1]~[R1]~1 | thujane scaffold |
| 33 | C~C2(~C)~[R]1~[R]~[R]~2~[R](~C)~[R]~[R]~1 | pinane scaffold |
| 34 | [!H]~[!H]2(~[!H])~[R]1~[R]~[R]~[R](~[!H])~2~[R]~[R]~1 | camphane scaffold |
| 35 | [!H]~[!H]2(~[!H])~[R]~[R](~[!H])1~[R]~[R]~2~[R]~[R]~1 | fenchane scaffold |
| 36 | C(-C)(-C)(-C)-C | quaternary carbon |
| 37 | C-C-C-C-C-C | six carbon single bond |
| 38 | C-C-C-C-C-C-C | seven carbon single bond chain |
| 39 | C-C-C-C-C-C-C-C | eight carbon single bond chain |
| 40 | C-C-C-C-C-C-C-C-C | nine carbon single bond chain |
| 41 | C-C-C-C-C-C-C-C-C-C | ten carbon single bond chain |
| 42 | C-C-C-C-C-C-C-C-C-C-C | eleven carbon single bond chain |

**Table S4. ‘SMARTS 42’ feature set.**

SMARTS42 fingerprints consisted of binary keys indicating the presence or absence of each feature. See Methods for additional explanation.
