## Supplemental Document S2, Complete Odor Map Atlas for "Mapping odorant sensitivities reveals a sparse but structured representation of olfactory chemical space by sensory input to the mouse olfactory bulb"

**Supplementary Material, Burton et al., 2022**

**Document S2. Functional Atlas of Sensory Inputs to the Dorsal Mouse Olfactory Bulb.**

Compilation of olfactory sensory neuron response maps to the 185-odorant panel imaged in each of four mice (OMP-tTA x TetO-GCaMP6s). See Methods for details of odorant presentation, imaging and map generation. Each map is scaled to its own maximum (mean of highest 65 pixels) and clipped at $\Delta F=0$. Concentrations are given as estimated delivered concentration to the mouse nose, rounded to the nearest order of magnitude to allow for deviations from ideal behavior of the odorant in its solvent. See Table S1 for more precise estimates of delivered concentration. Grid overlay is for facilitating visual comparison across maps. Odorants are grouped by major structural features and are numbered in order of presentation in Main figures and Table S1. Letter-number identifier matches that in Fig. 5A and in Table S1. Maps with no measured response in any glomerulus of a given mouse are shown at 50% opacity. Odorants failing to evoke a response in all 4 mice are indicated with text.

### Carboxylic acid

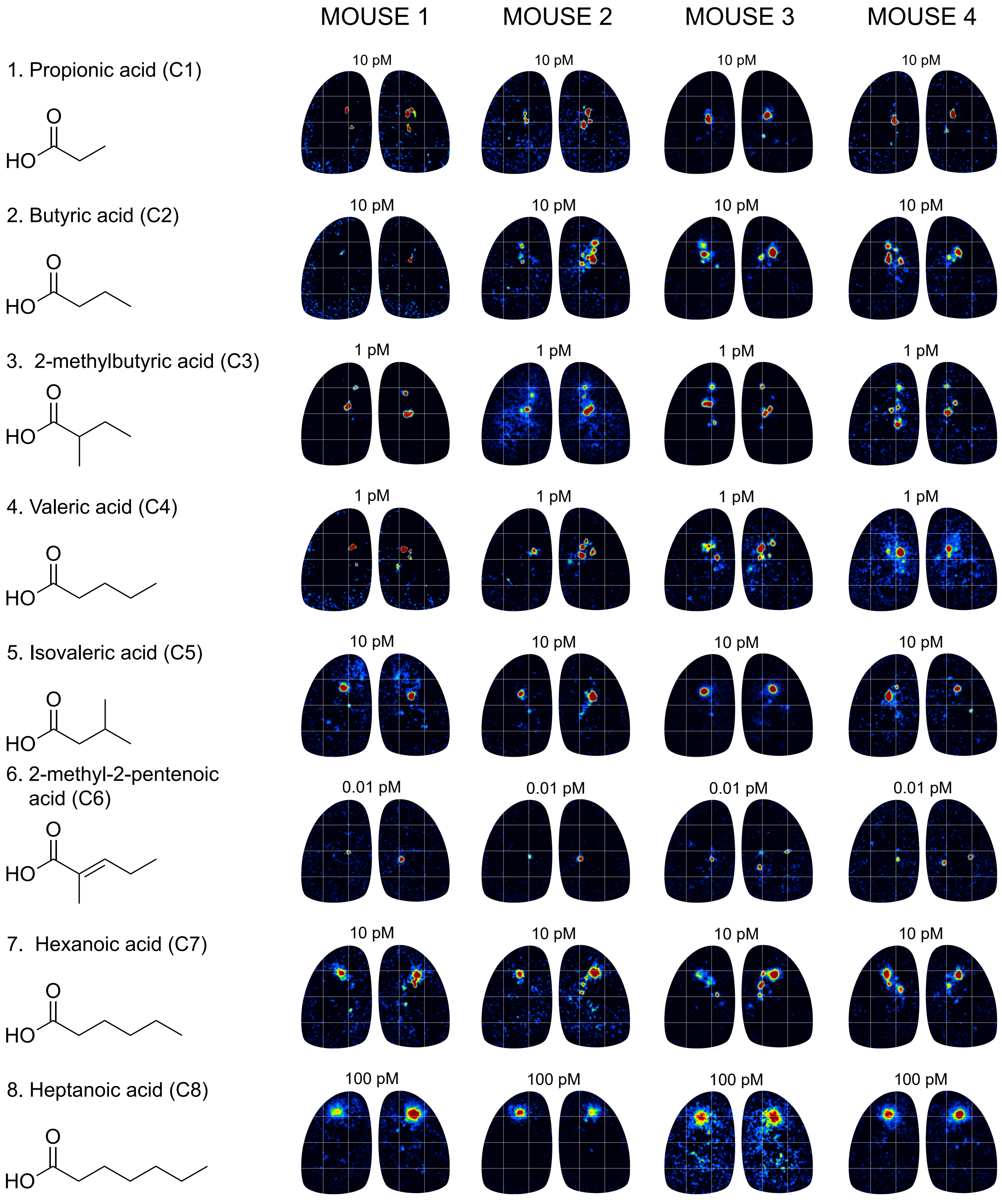

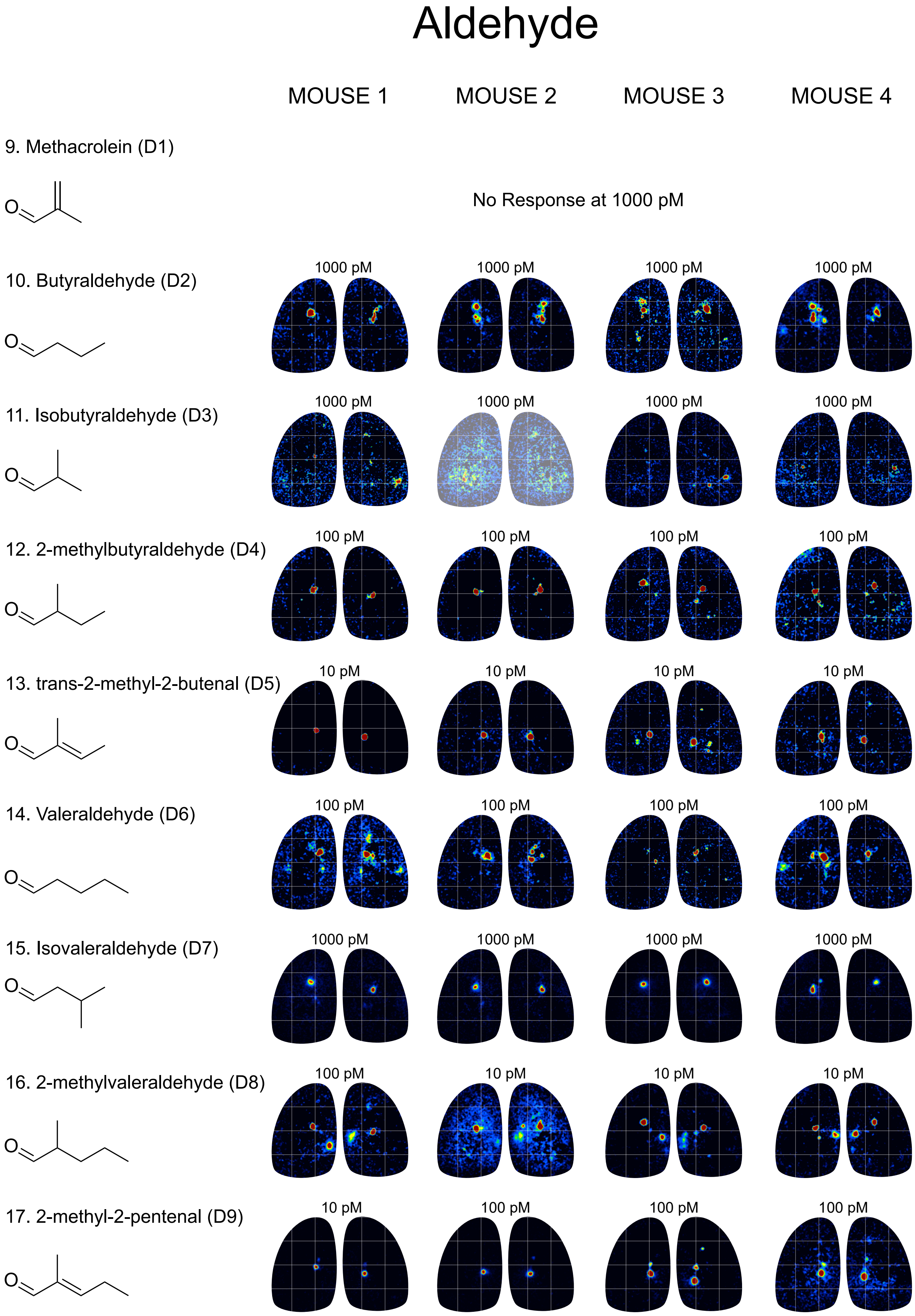
Aldehyde
MOUSE 1
MOUSE 2
MOUSE 3
MOUSE 4
9. Methacrolein (D1)
O
No Response at 1000 pM
10. Butyraldehyde (D2)
O
1000 pM
1000 pM
1000 pM
1000 pM
11. Isobutyraldehyde (D3)
O
1000 pM
1000 pM
1000 pM
1000 pM
12. 2-methylbutyraldehyde (D4)
O
100 pM
100 pM
100 pM
100 pM
13. trans-2-methyl-2-butenal (D5)
O
10 pM
10 pM
10 pM
10 pM
14. Valeraldehyde (D6)
O
100 pM
100 pM
100 pM
100 pM
15. Isovaleraldehyde (D7)
O
1000 pM
1000 pM
1000 pM
1000 pM
16. 2-methylvaleraldehyde (D8)
O
100 pM
10 pM
10 pM
10 pM
17. 2-methyl-2-pentenal (D9)
O
10 pM
100 pM
100 pM
100 pM

### Aldehyde II

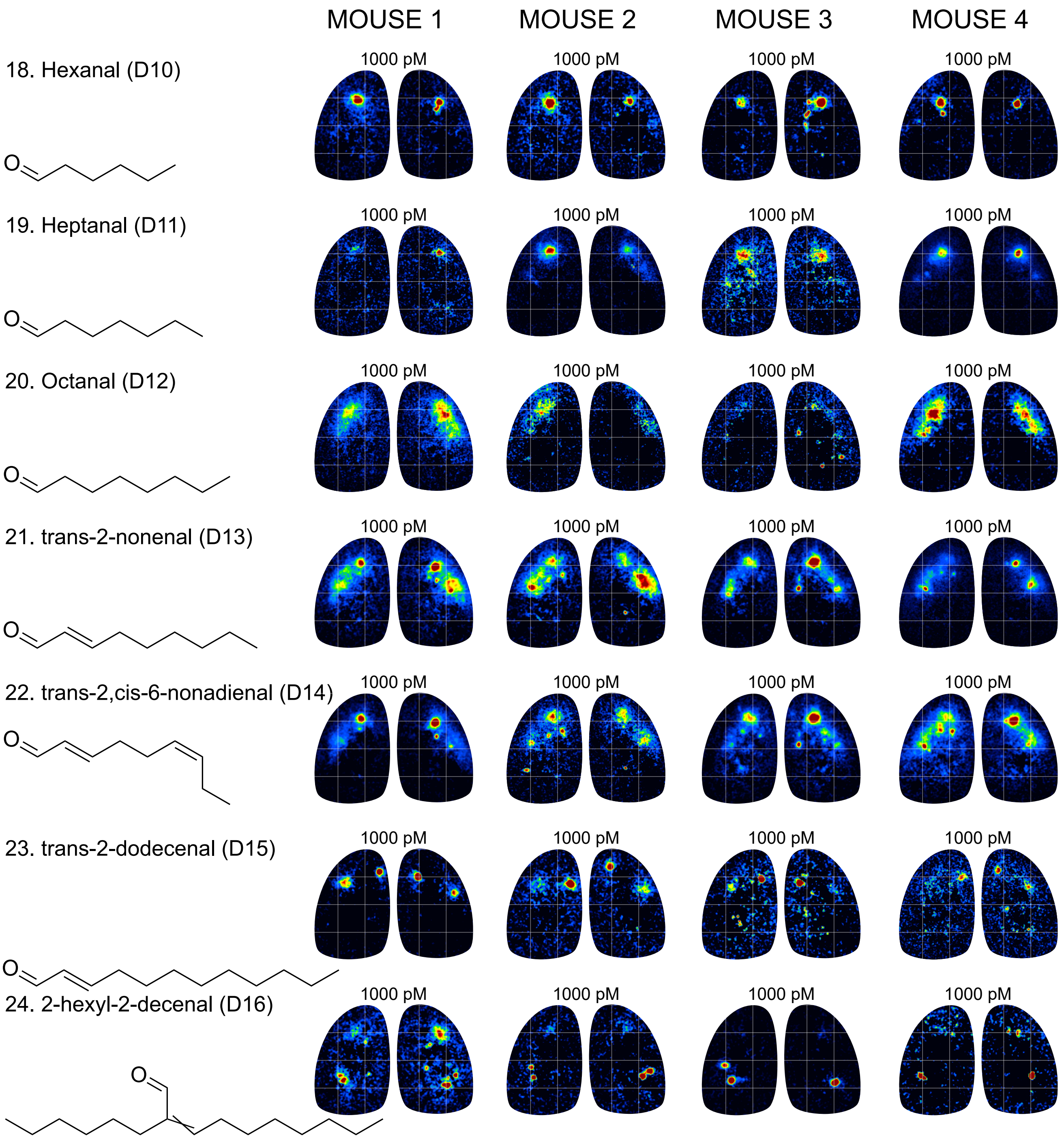

MOUSE 1
MOUSE 2
MOUSE 3
MOUSE 4
18. Hexanal (D10)
19. Heptanal (D11)
20. Octanal (D12)
21. trans-2-nonenal (D13)
22. trans-2,cis-6-nonadienal (D14)
23. trans-2-dodecenal (D15)
24. 2-hexyl-2-decenal (D16)
1000 pM
O

### Ester

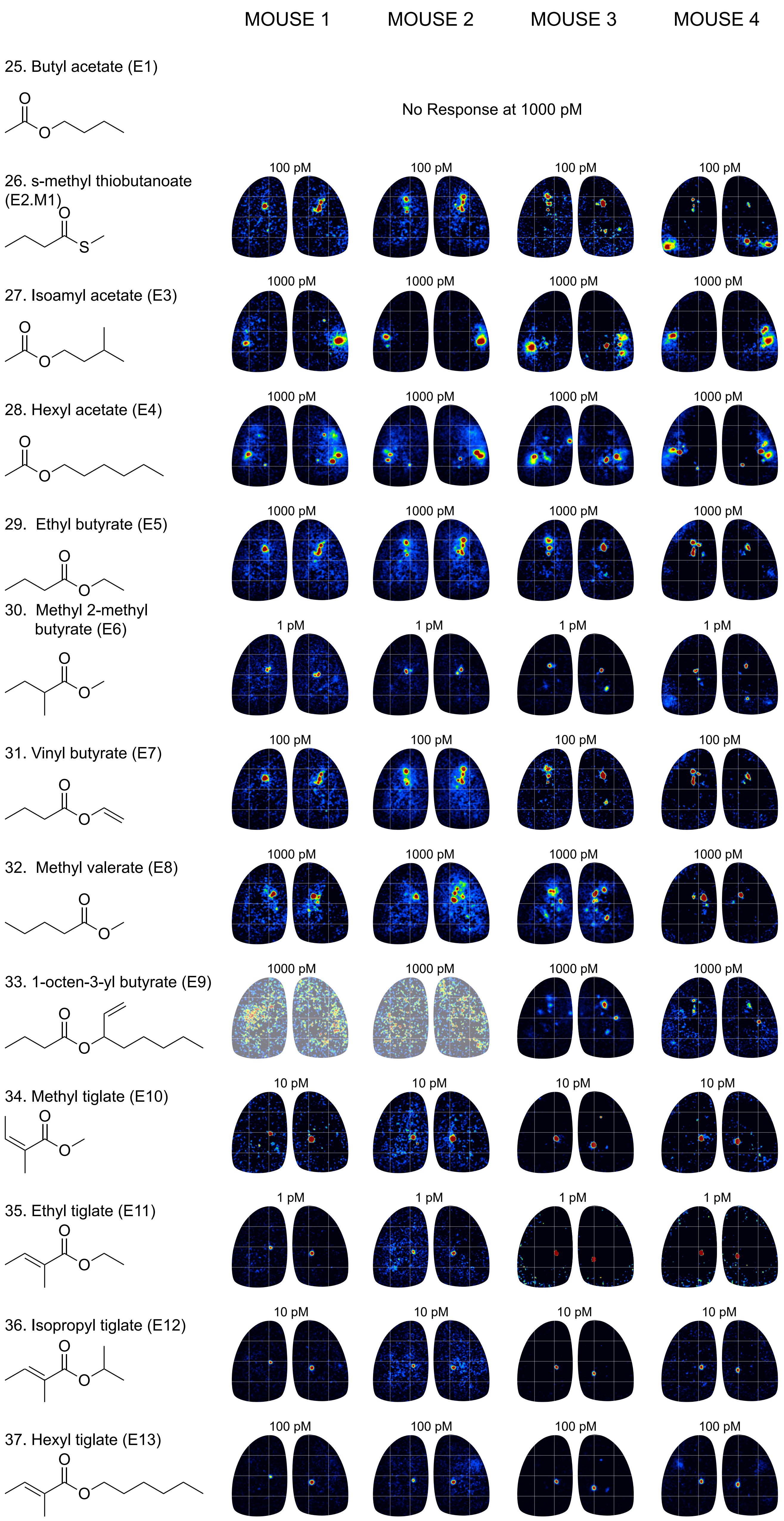

MOUSE 1
MOUSE 2
MOUSE 3
MOUSE 4
25. Butyl acetate (E1)
No Response at 1000 pM
26. s-methyl thiobutanoate (E2.M1)
100 pM
27. Isoamyl acetate (E3)
1000 pM
28. Hexyl acetate (E4)
1000 pM
29. Ethyl butyrate (E5)
1000 pM
30. Methyl 2-methyl butyrate (E6)
1 pM
31. Vinyl butyrate (E7)
100 pM
32. Methyl valerate (E8)
1000 pM
33. 1-octen-3-yl butyrate (E9)
1000 pM
34. Methyl tiglate (E10)
10 pM
35. Ethyl tiglate (E11)
1 pM
36. Isopropyl tiglate (E12)
10 pM
37. Hexyl tiglate (E13)
100 pM

### Ketone

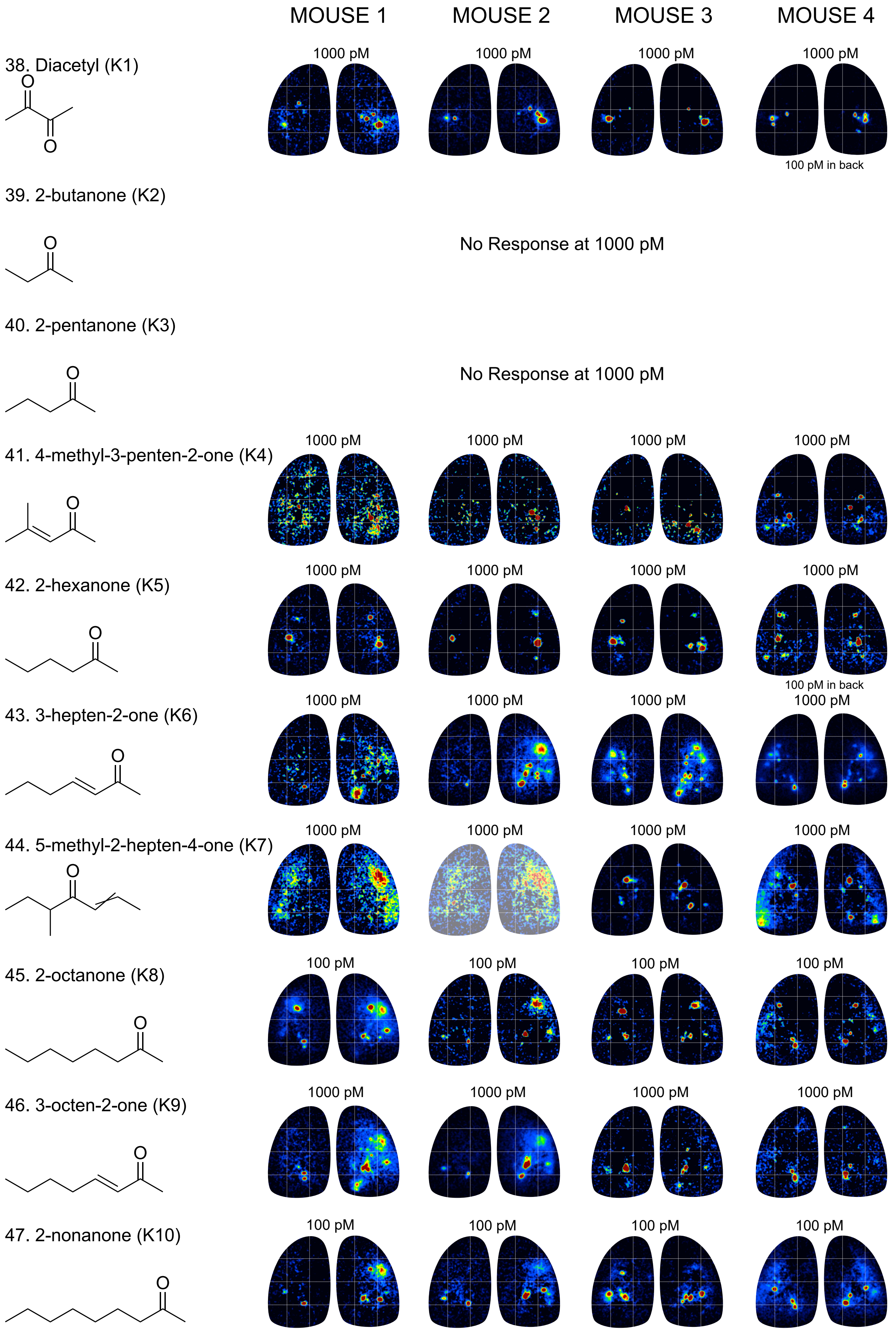
MOUSE 1
MOUSE 2
MOUSE 3
MOUSE 4
38. Diacetyl (K1)
1000 pM
1000 pM
1000 pM
1000 pM
100 pM in back
39. 2-butanone (K2)
No Response at 1000 pM
40. 2-pentanone (K3)
No Response at 1000 pM
41. 4-methyl-3-penten-2-one (K4)
1000 pM
1000 pM
1000 pM
1000 pM
42. 2-hexanone (K5)
1000 pM
1000 pM
1000 pM
1000 pM
100 pM in back
43. 3-hepten-2-one (K6)
1000 pM
1000 pM
1000 pM
1000 pM
44. 5-methyl-2-hepten-4-one (K7)
1000 pM
1000 pM
1000 pM
1000 pM
45. 2-octanone (K8)
100 pM
100 pM
100 pM
100 pM
46. 3-octen-2-one (K9)
1000 pM
1000 pM
1000 pM
1000 pM
47. 2-nonanone (K10)
100 pM
100 pM
100 pM
100 pM

### Alcohol

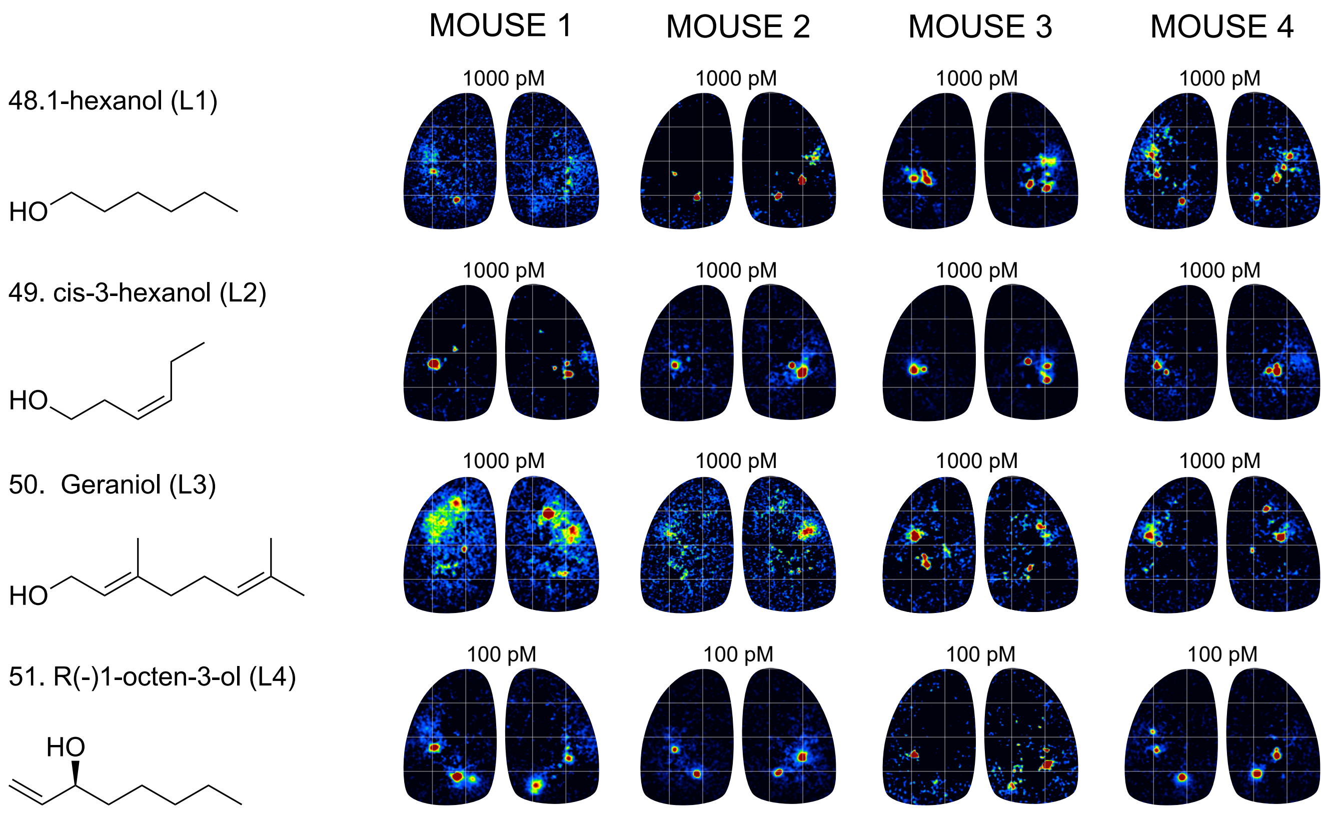


### Aromatic Aldehyde

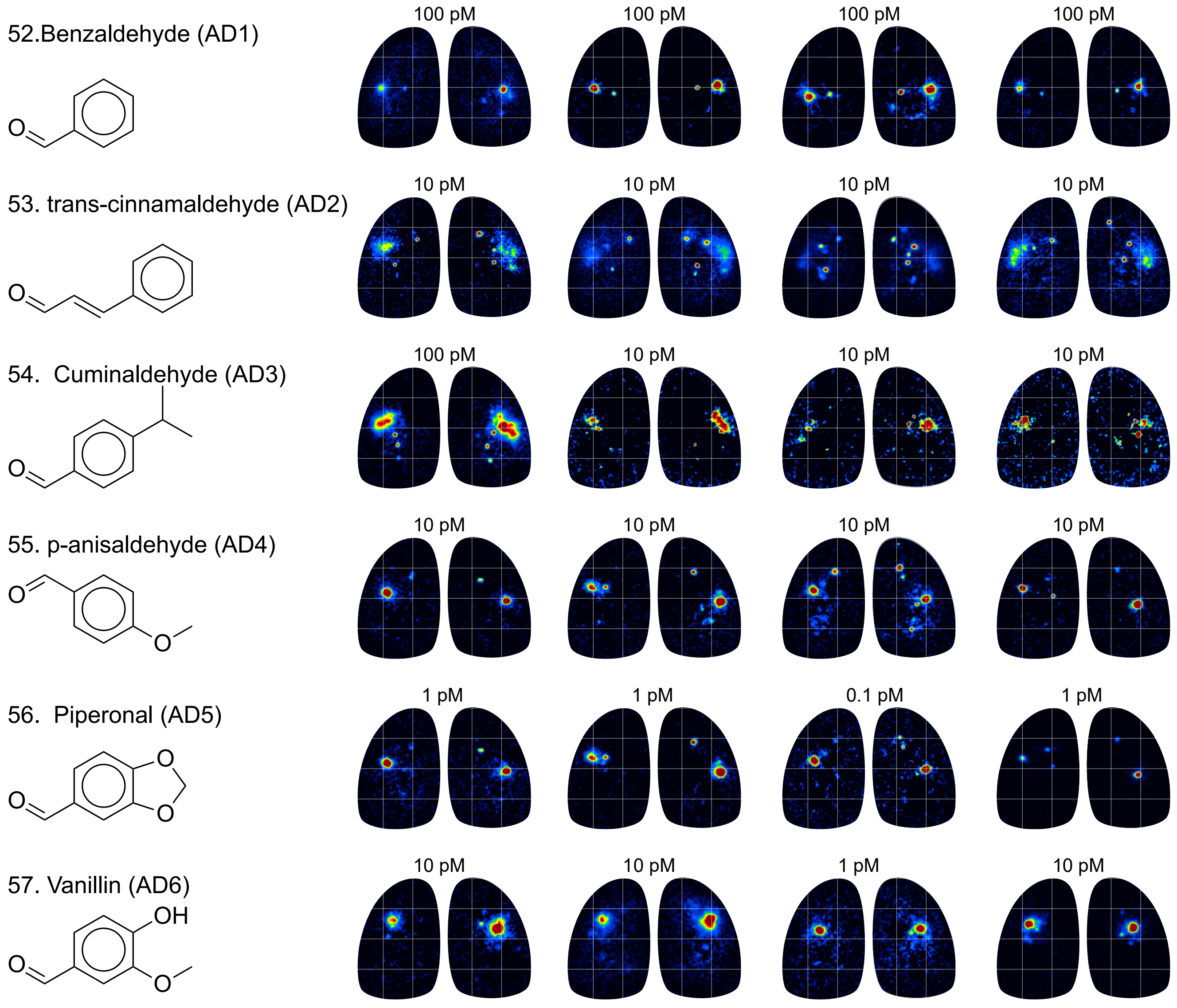

### Aromatic Ketone

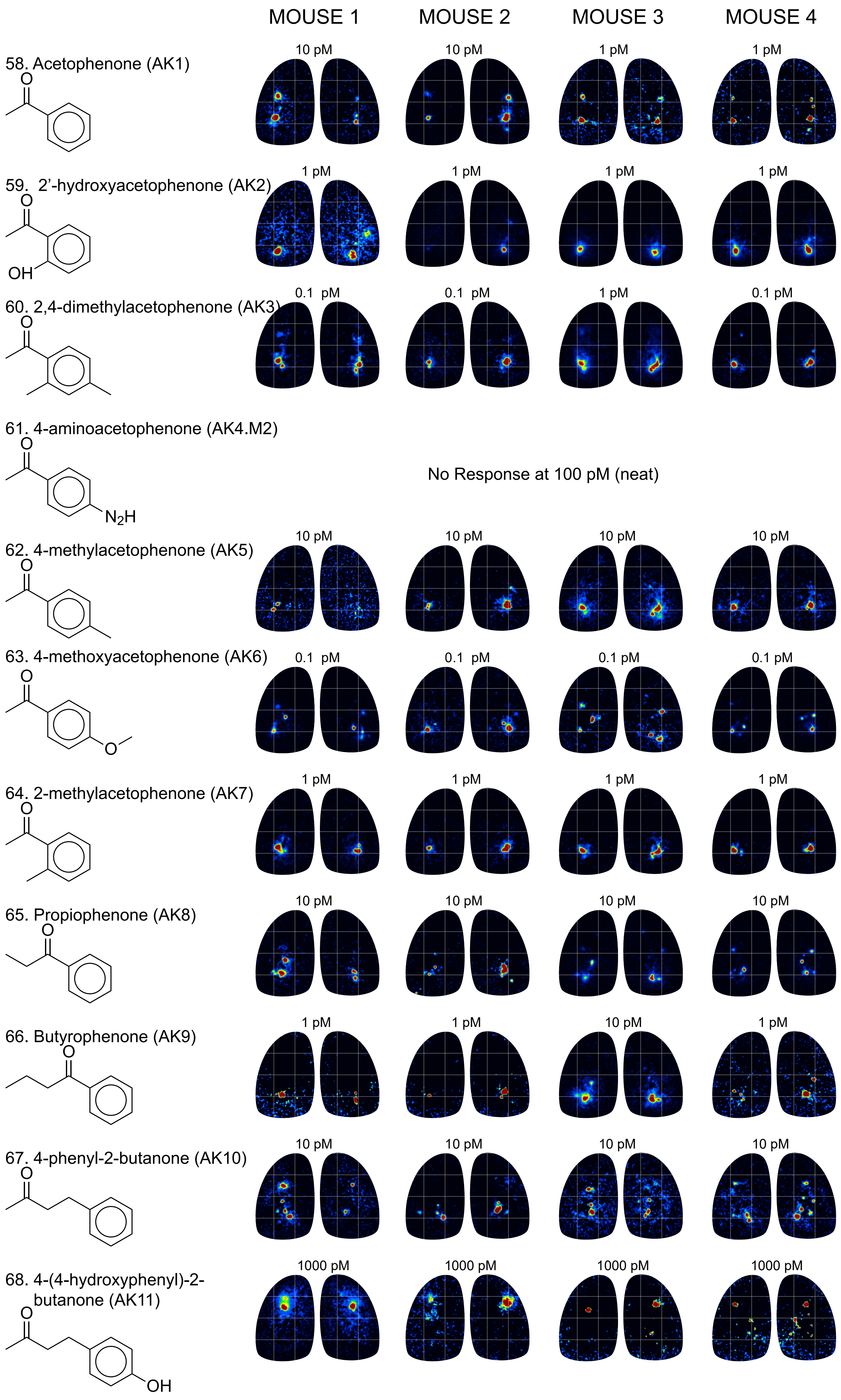

### Aromatic Ester

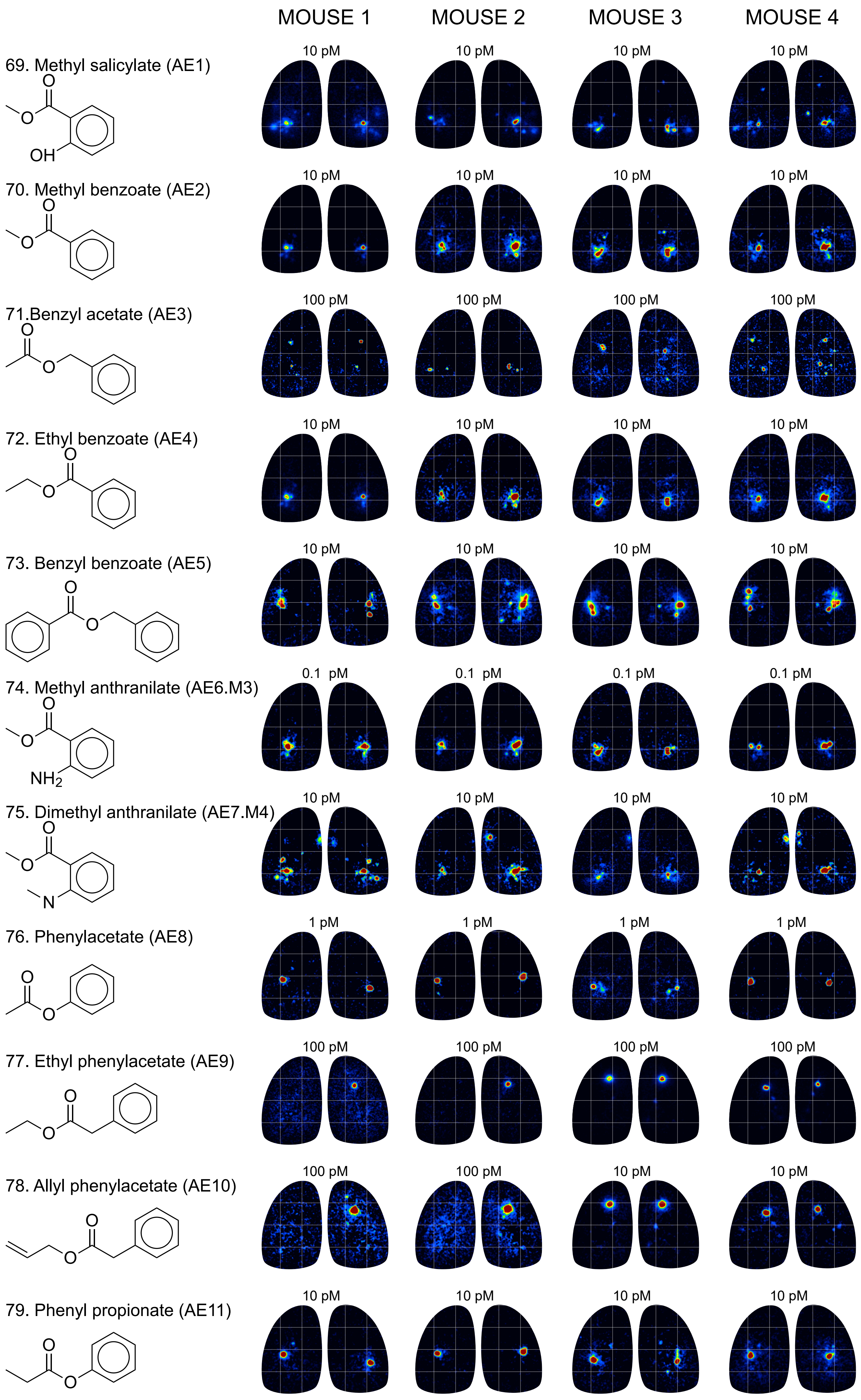

### Aromatic Phenol/Methoxy

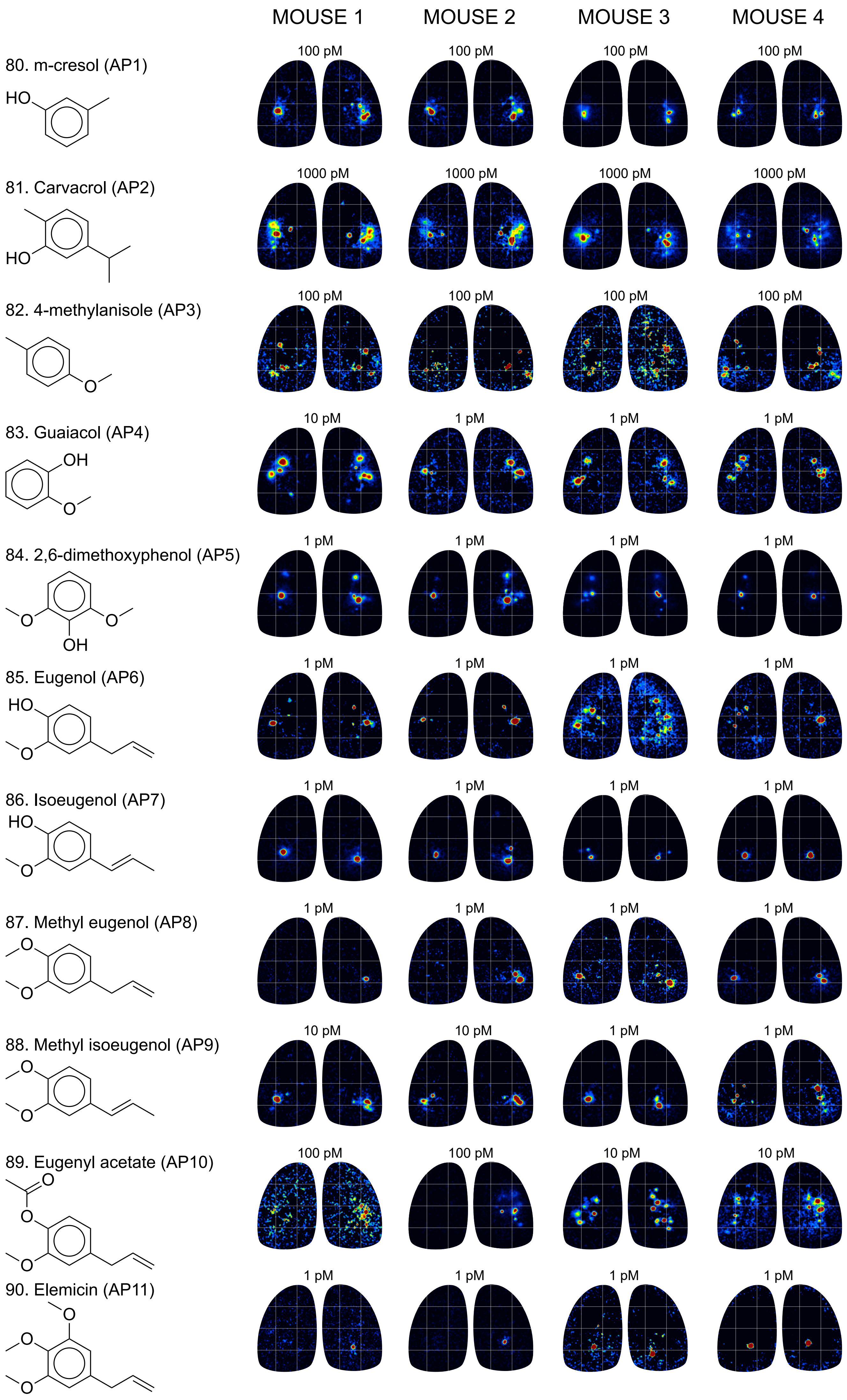

### Cyclic Terpenoid I

91. Fenchol (CT1)

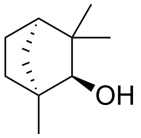


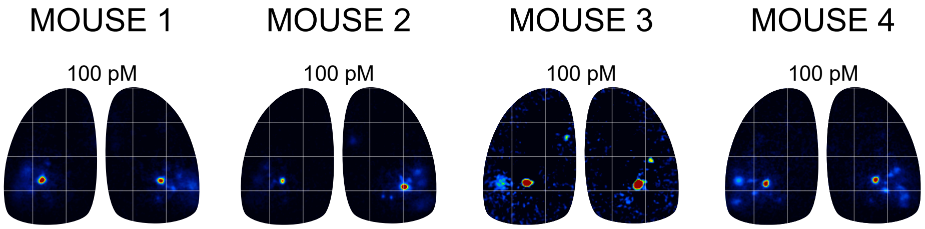


92. 2-ethyl fenchol (CT2)

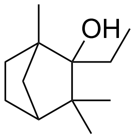


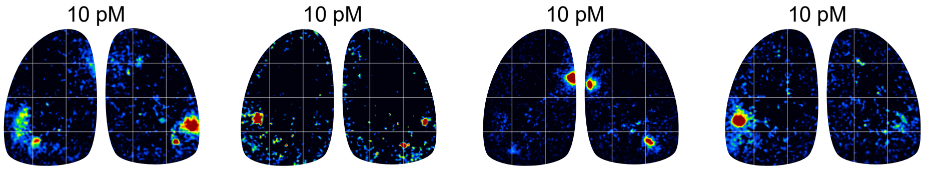


93. α-pinene (CT3)

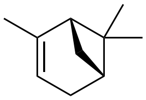

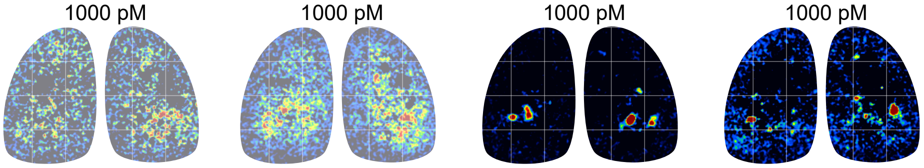


94. Isobornyl isovalerate (CT4.M5)

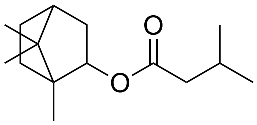


No Response at 1000 pM

95. 1,8-cineole (CT5)

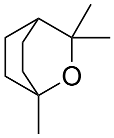


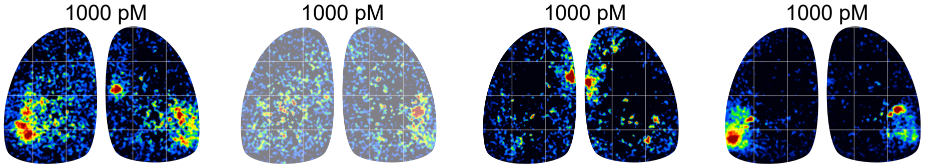


96. L-carvone (CT6)

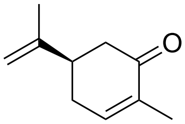


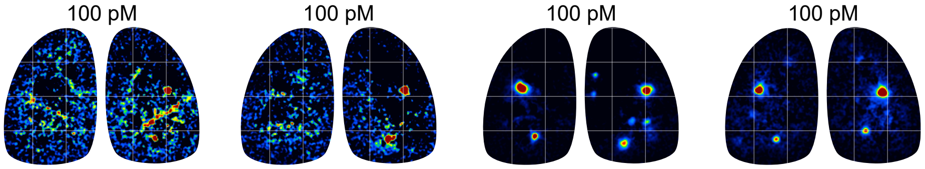


97. β-Ionone (CT7)

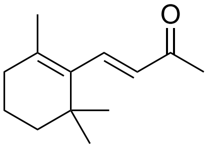


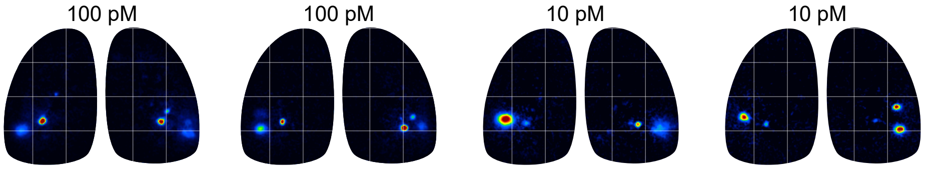


98. β-damascone (CT8)

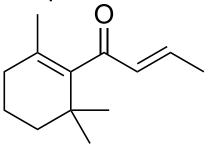


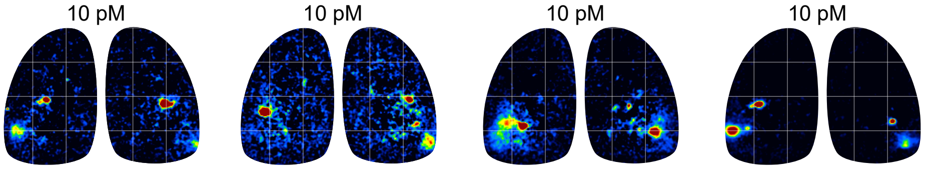


99. Damascenone (CT9)

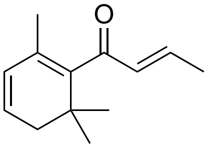


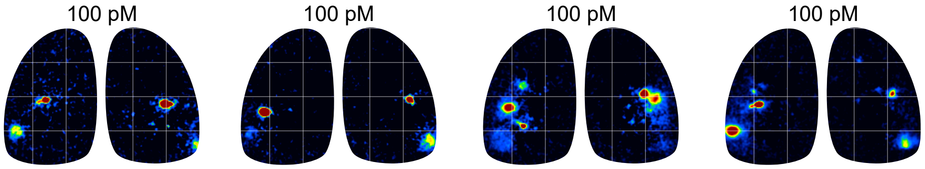

### Cyclic Terpenoid II

MOUSE 1 | MOUSE 2 | MOUSE 3 | MOUSE 4

100. Menthone (CT10)

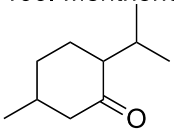

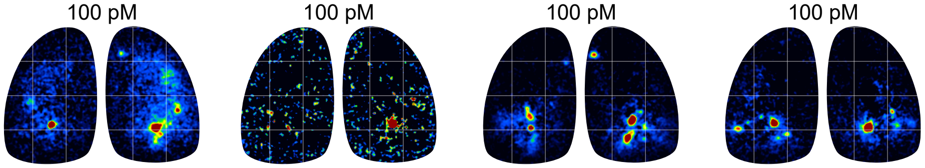


101. (R)-(+)-pulegone (CT11)

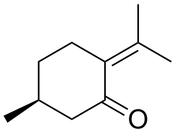

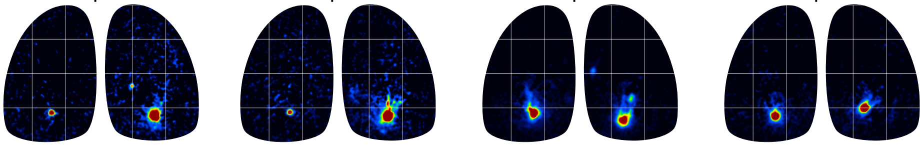


102. (+)-isomethone (CT12)

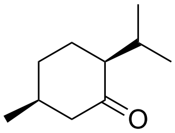

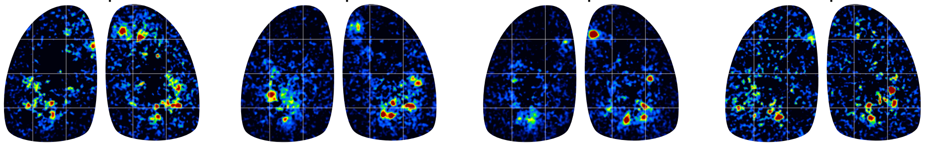


103. Nootkatone (CT13)

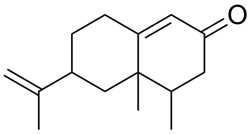

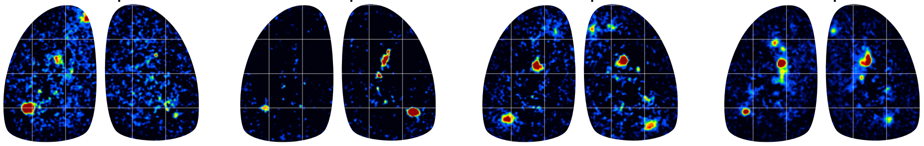


104. (+/-)-menthol (CT14)

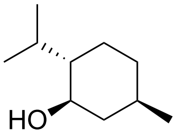

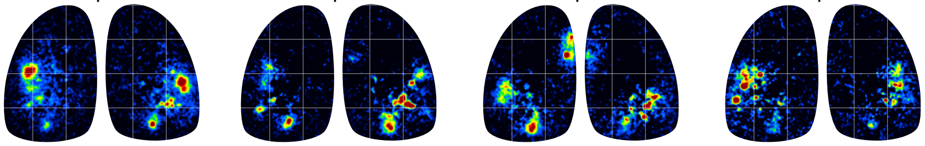


105. (+)-neomenthol (CT15)

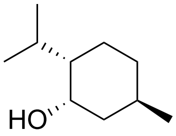

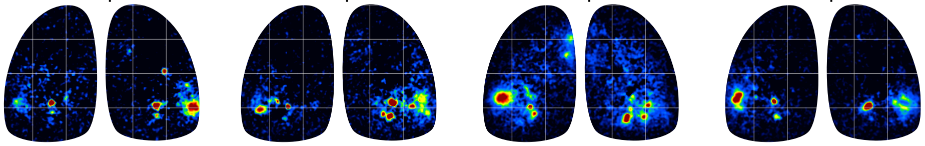


106. (+/-)-geosmin (CT16)

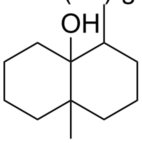

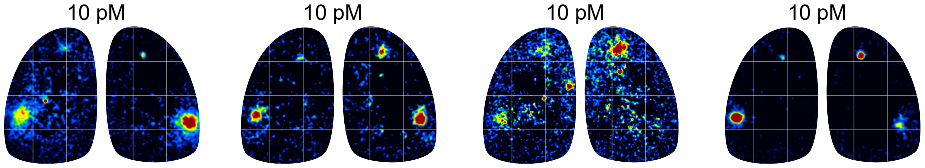


107. (-)-ambroxide (CT17)

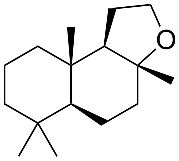

No response at 1000 pM

108. (+)-menthofuran (CT18)

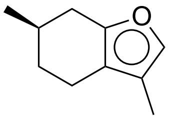

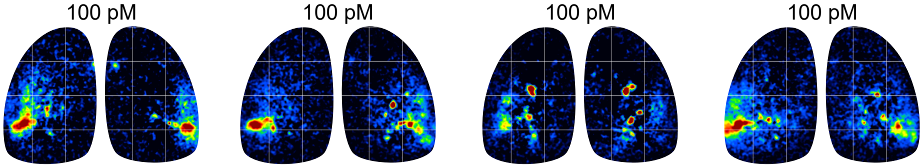

### Furan/Pyrone/Lactone

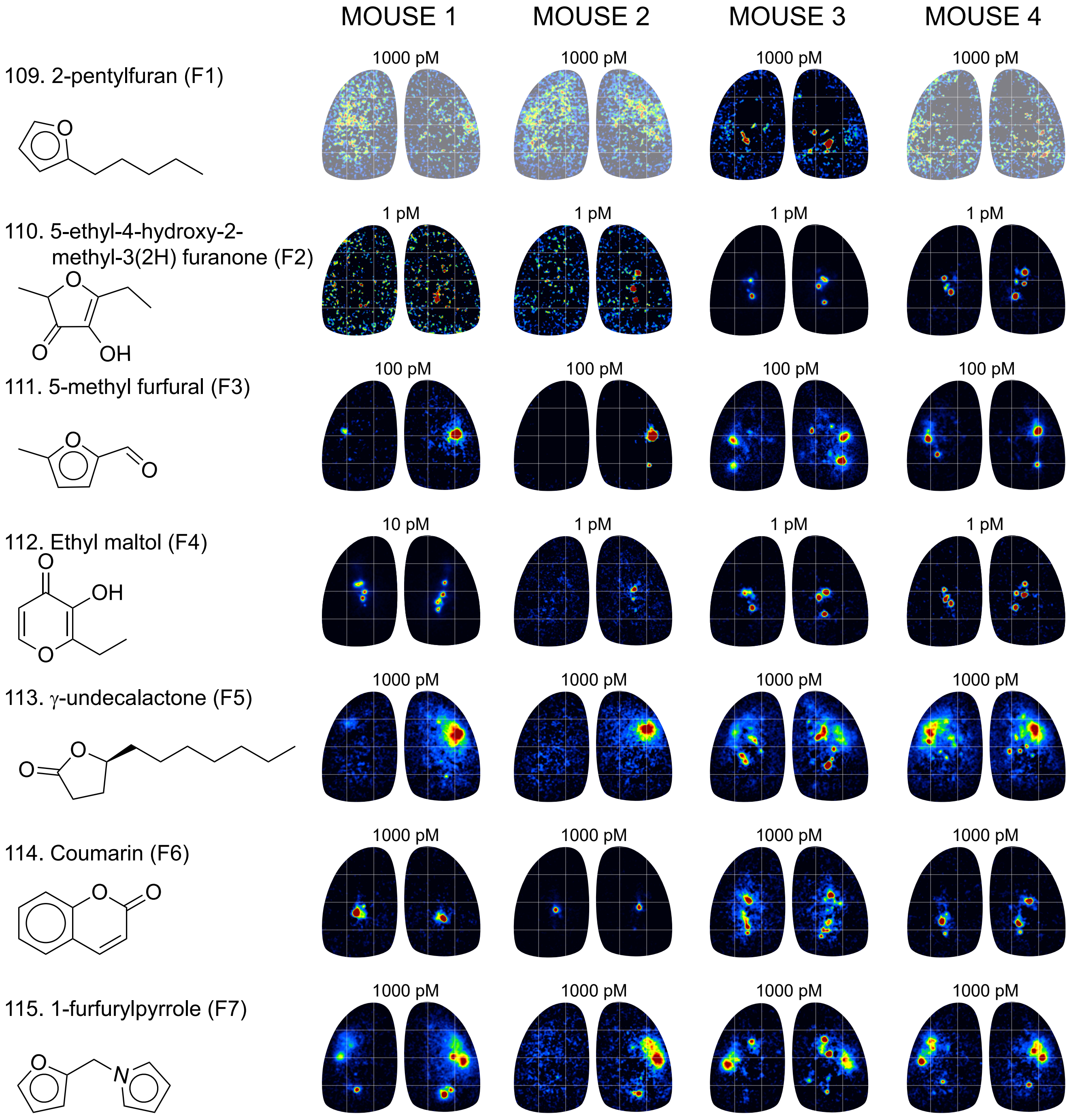

### Amine I

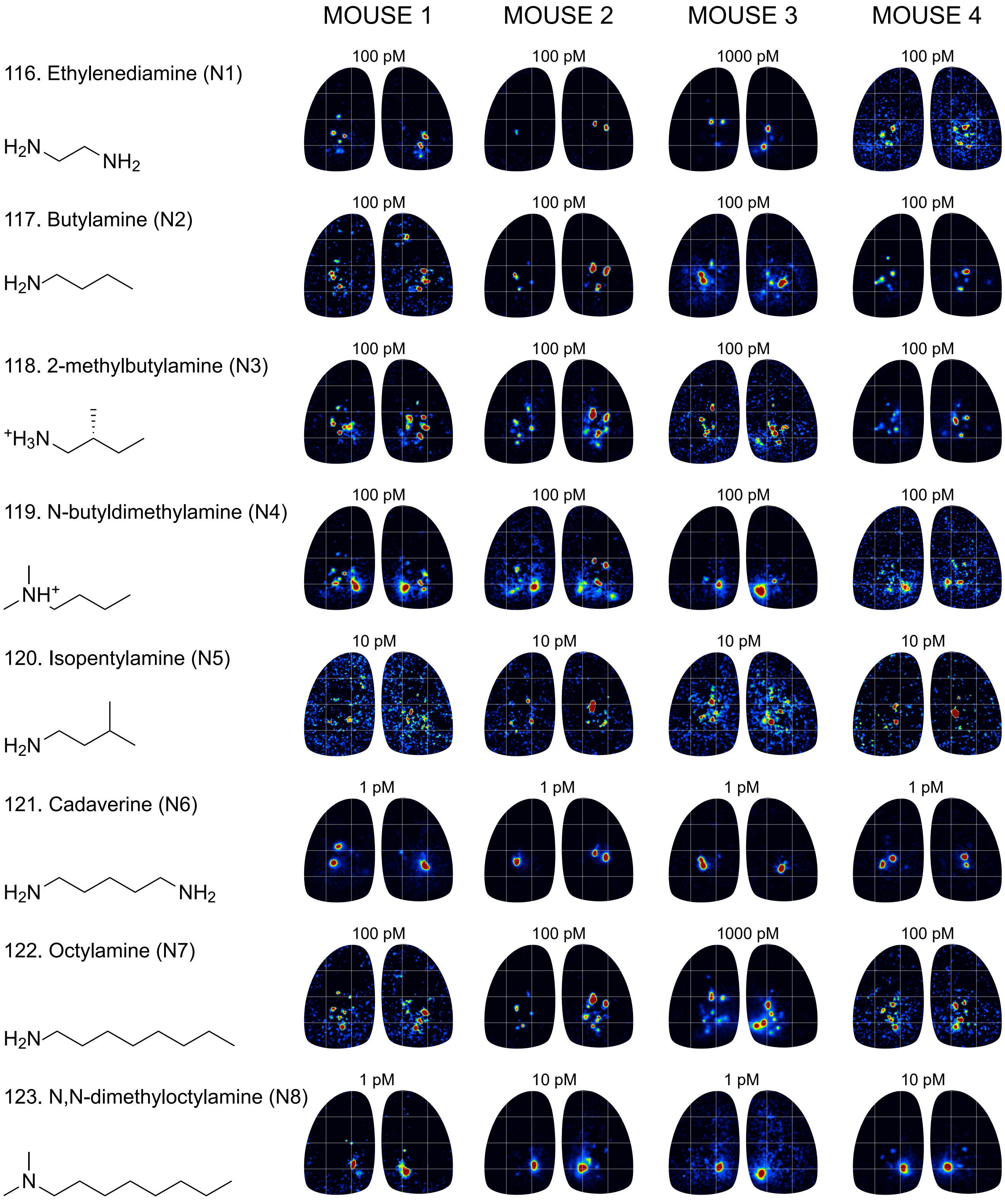
MOUSE 1
MOUSE 2
MOUSE 3
MOUSE 4
116. Ethylenediamine (N1)
100 pM
100 pM
1000 pM
100 pM
H2N
NH2
117. Butylamine (N2)
100 pM
100 pM
100 pM
100 pM
H2N
118. 2-methylbutylamine (N3)
100 pM
100 pM
100 pM
100 pM
+H3N
119. N-butyldimethylamine (N4)
100 pM
100 pM
100 pM
100 pM
NH+
120. Isopentylamine (N5)
10 pM
10 pM
10 pM
10 pM
H2N
121. Cadaverine (N6)
1 pM
1 pM
1 pM
1 pM
H2N
NH2
122. Octylamine (N7)
100 pM
100 pM
1000 pM
100 pM
H2N
123. N,N-dimethyloctylamine (N8)
1 pM
10 pM
1 pM
10 pM
N

### Amine II

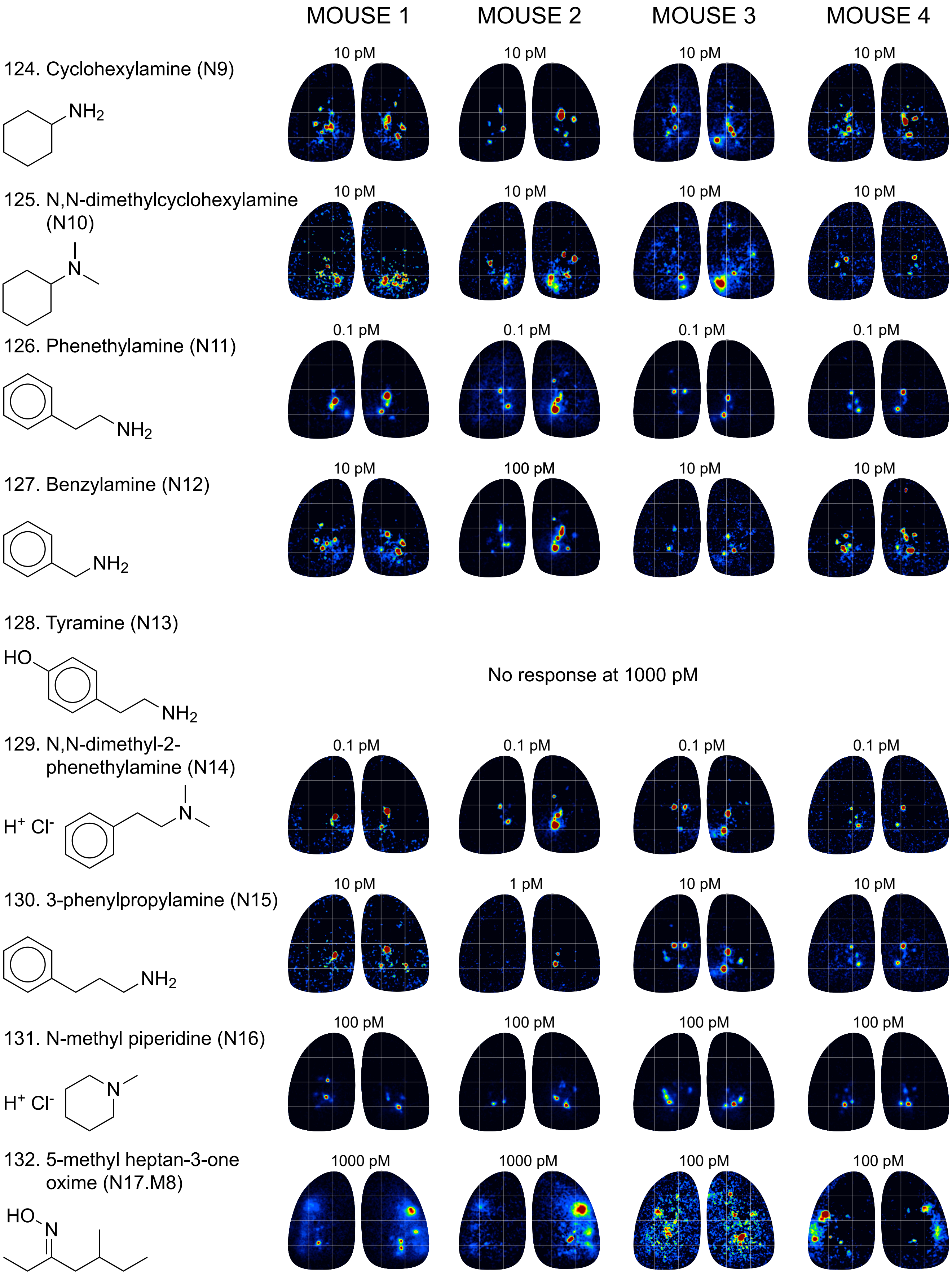

MOUSE 1
MOUSE 2
MOUSE 3
MOUSE 4
124. Cyclohexylamine (N9)
NH2
10 pM
10 pM
10 pM
10 pM
125. N,N-dimethylcyclohexylamine (N10)
N
10 pM
10 pM
10 pM
10 pM
126. Phenethylamine (N11)
NH2
0.1 pM
0.1 pM
0.1 pM
0.1 pM
127. Benzylamine (N12)
NH2
10 pM
100 pM
10 pM
10 pM
128. Tyramine (N13)
HO
NH2
No response at 1000 pM
129. N,N-dimethyl-2-phenethylamine (N14)
H+ Cl-
N
0.1 pM
0.1 pM
0.1 pM
0.1 pM
130. 3-phenylpropylamine (N15)
NH2
10 pM
1 pM
10 pM
10 pM
131. N-methyl piperidine (N16)
H+ Cl-
N
100 pM
100 pM
100 pM
100 pM
132. 5-methyl heptan-3-one oxime (N17.M8)
HO
N
1000 pM
1000 pM
100 pM
100 pM

### Pyrazine I

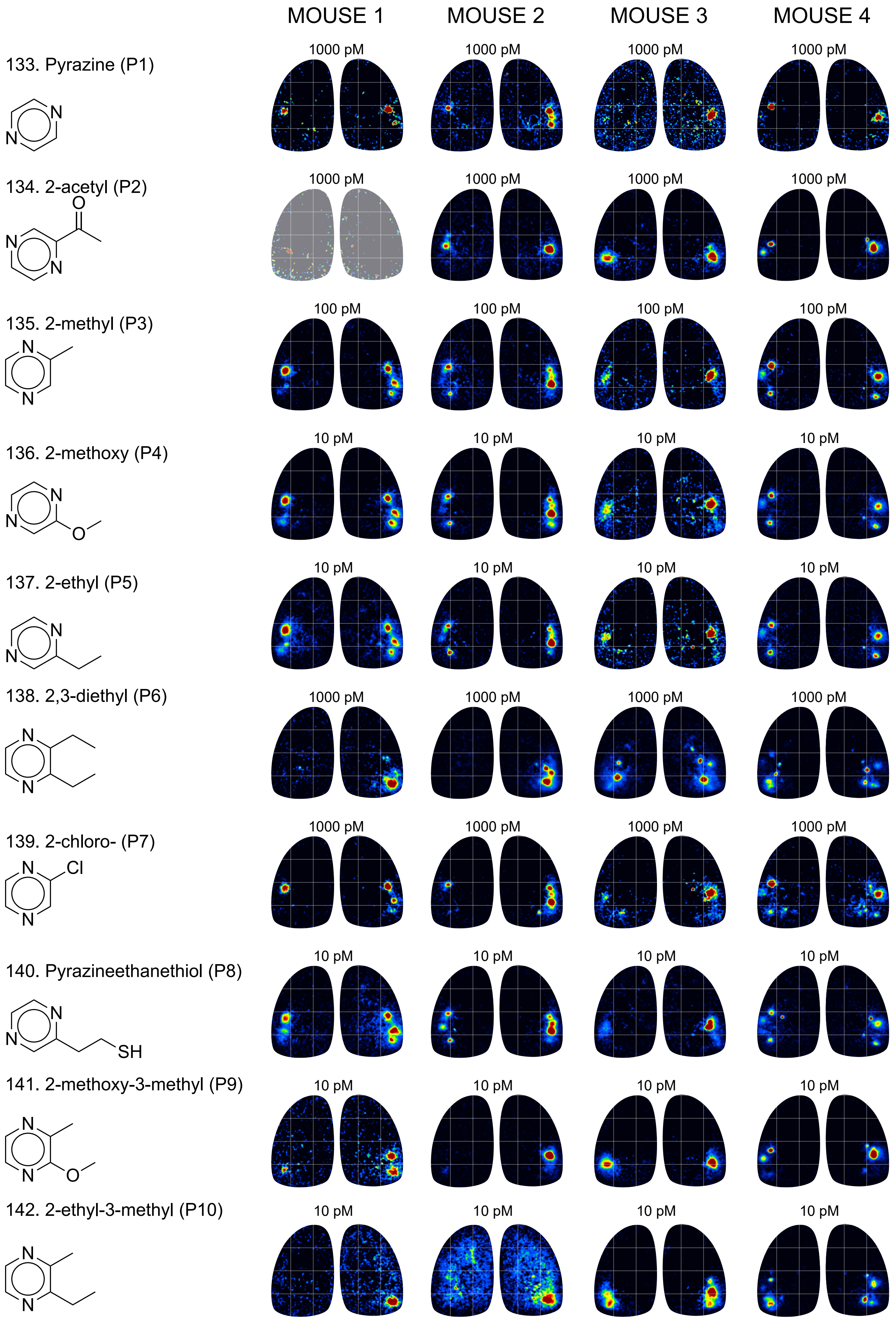

MOUSE 1
MOUSE 2
MOUSE 3
MOUSE 4
133. Pyrazine (P1)
1000 pM
1000 pM
1000 pM
1000 pM
134. 2-acetyl (P2)
1000 pM
1000 pM
1000 pM
1000 pM
135. 2-methyl (P3)
100 pM
100 pM
100 pM
100 pM
136. 2-methoxy (P4)
10 pM
10 pM
10 pM
10 pM
137. 2-ethyl (P5)
10 pM
10 pM
10 pM
10 pM
138. 2,3-diethyl (P6)
1000 pM
1000 pM
1000 pM
1000 pM
139. 2-chloro- (P7)
1000 pM
1000 pM
1000 pM
1000 pM
140. Pyrazineethanethiol (P8)
10 pM
10 pM
10 pM
10 pM
141. 2-methoxy-3-methyl (P9)
10 pM
10 pM
10 pM
10 pM
142. 2-ethyl-3-methyl (P10)
10 pM
10 pM
10 pM
10 pM

### Pyrazine II

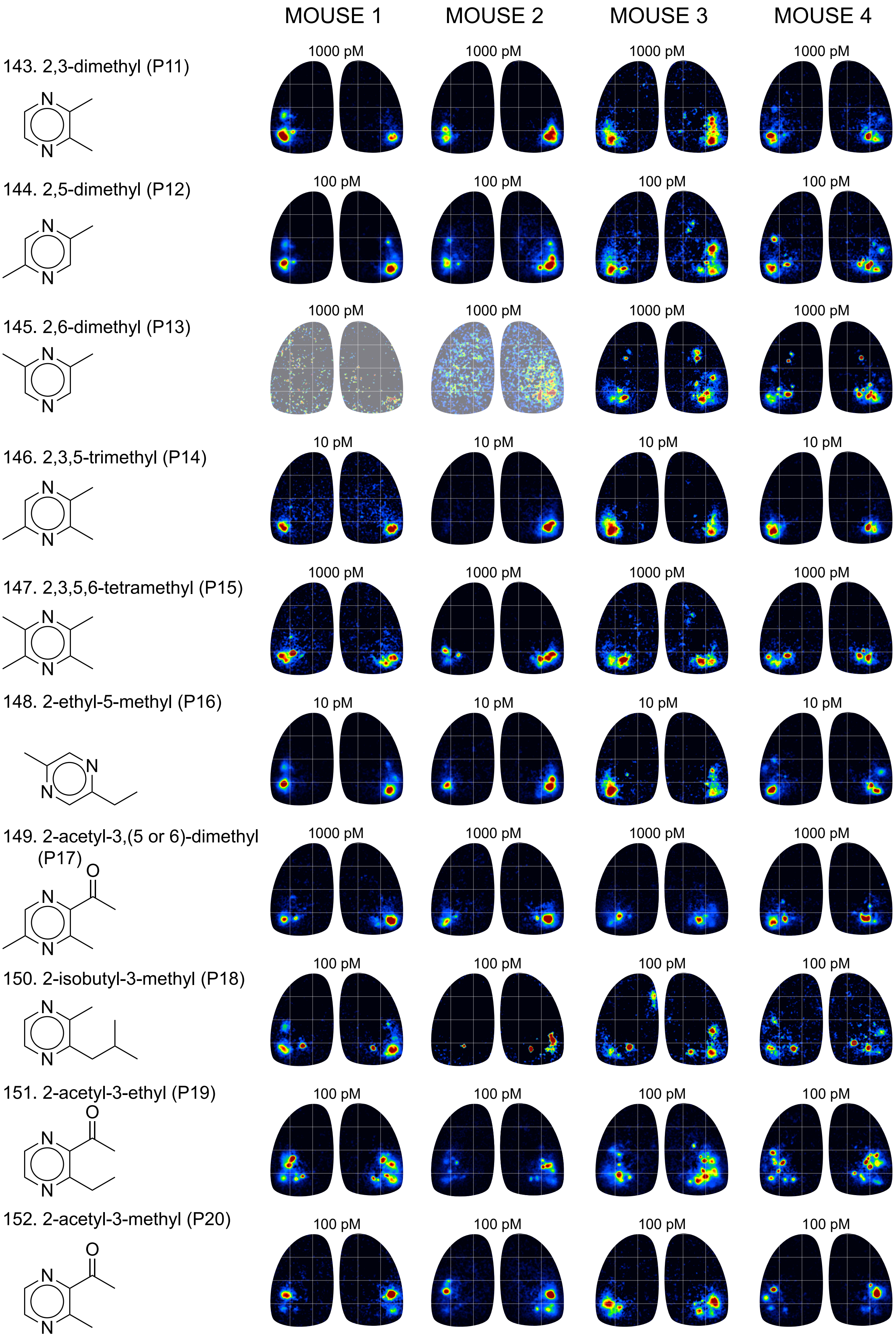
MOUSE 1
MOUSE 2
MOUSE 3
MOUSE 4
143. 2,3-dimethyl (P11)
1000 pM
1000 pM
1000 pM
1000 pM
144. 2,5-dimethyl (P12)
100 pM
100 pM
100 pM
100 pM
145. 2,6-dimethyl (P13)
1000 pM
1000 pM
1000 pM
1000 pM
146. 2,3,5-trimethyl (P14)
10 pM
10 pM
10 pM
10 pM
147. 2,3,5,6-tetramethyl (P15)
1000 pM
1000 pM
1000 pM
1000 pM
148. 2-ethyl-5-methyl (P16)
10 pM
10 pM
10 pM
10 pM
149. 2-acetyl-3,(5 or 6)-dimethyl (P17)
1000 pM
1000 pM
1000 pM
1000 pM
150. 2-isobutyl-3-methyl (P18)
100 pM
100 pM
100 pM
100 pM
151. 2-acetyl-3-ethyl (P19)
100 pM
100 pM
100 pM
100 pM
152. 2-acetyl-3-methyl (P20)
100 pM
100 pM
100 pM
100 pM

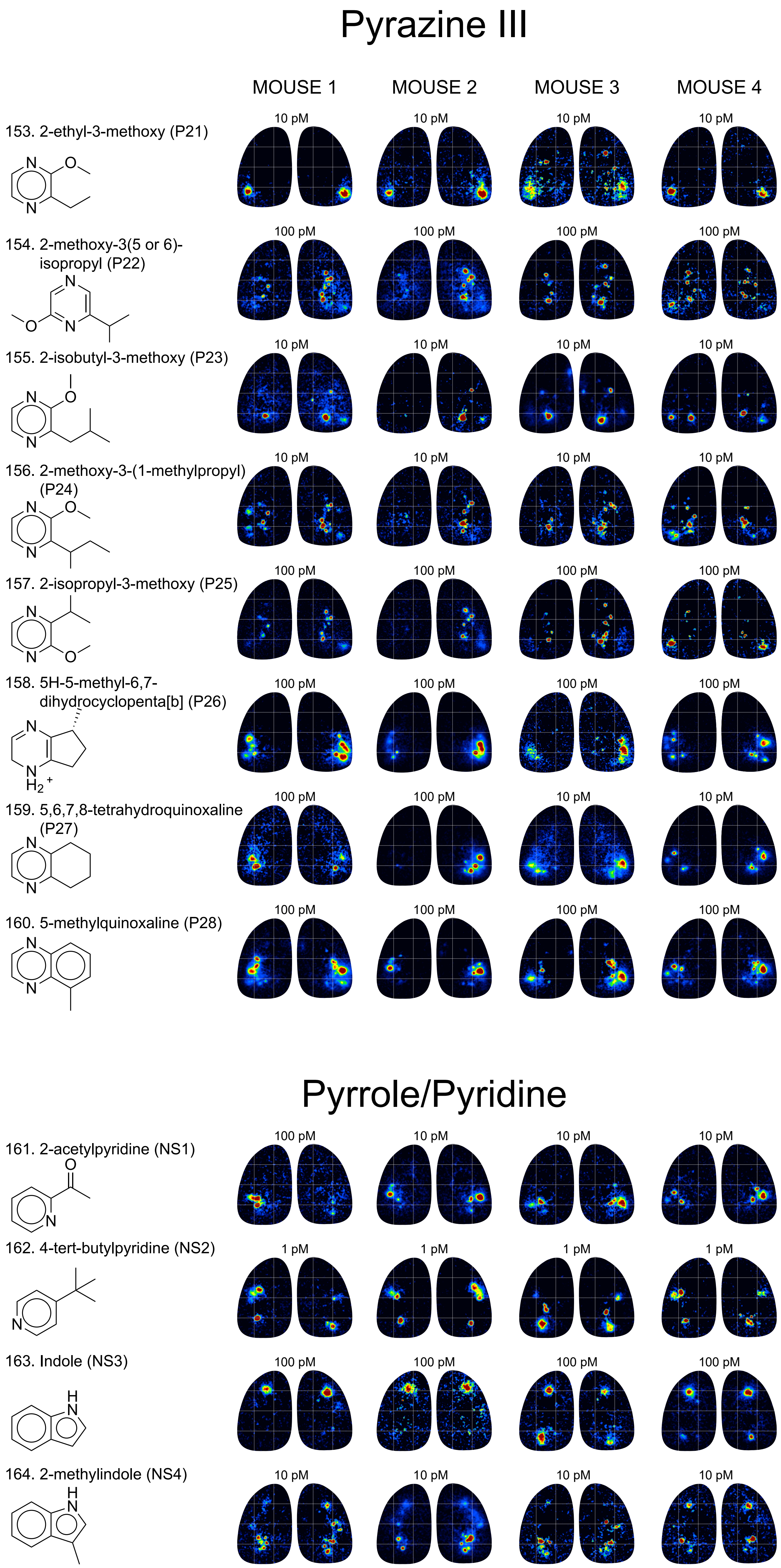

Pyrazine III
MOUSE 1
MOUSE 2
MOUSE 3
MOUSE 4
153. 2-ethyl-3-methoxy (P21)
10 pM
154. 2-methoxy-3(5 or 6)-isopropyl (P22)
100 pM
155. 2-isobutyl-3-methoxy (P23)
10 pM
156. 2-methoxy-3-(1-methylpropyl) (P24)
10 pM
157. 2-isopropyl-3-methoxy (P25)
100 pM
158. 5H-5-methyl-6,7-dihydrocyclopenta[b] (P26)
100 pM
159. 5,6,7,8-tetrahydroquinoxaline (P27)
100 pM
10 pM
160. 5-methylquinoxaline (P28)
100 pM
Pyrrole/Pyridine
161. 2-acetylpyridine (NS1)
100 pM
10 pM
162. 4-tert-butylpyridine (NS2)
1 pM
163. Indole (NS3)
100 pM
164. 2-methylindole (NS4)
10 pM

### Thiazole/Thiane

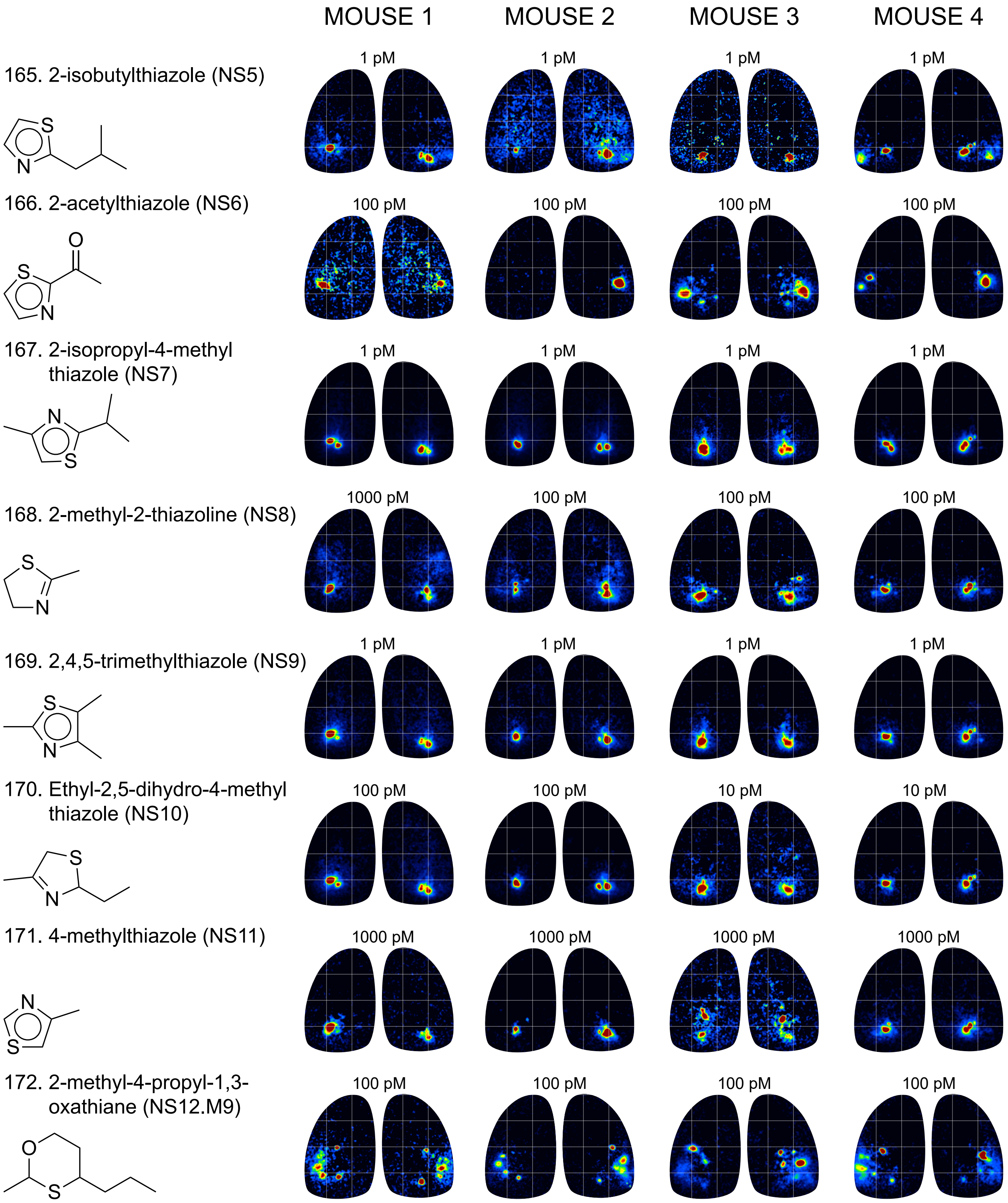

### Sulfide/Thiol

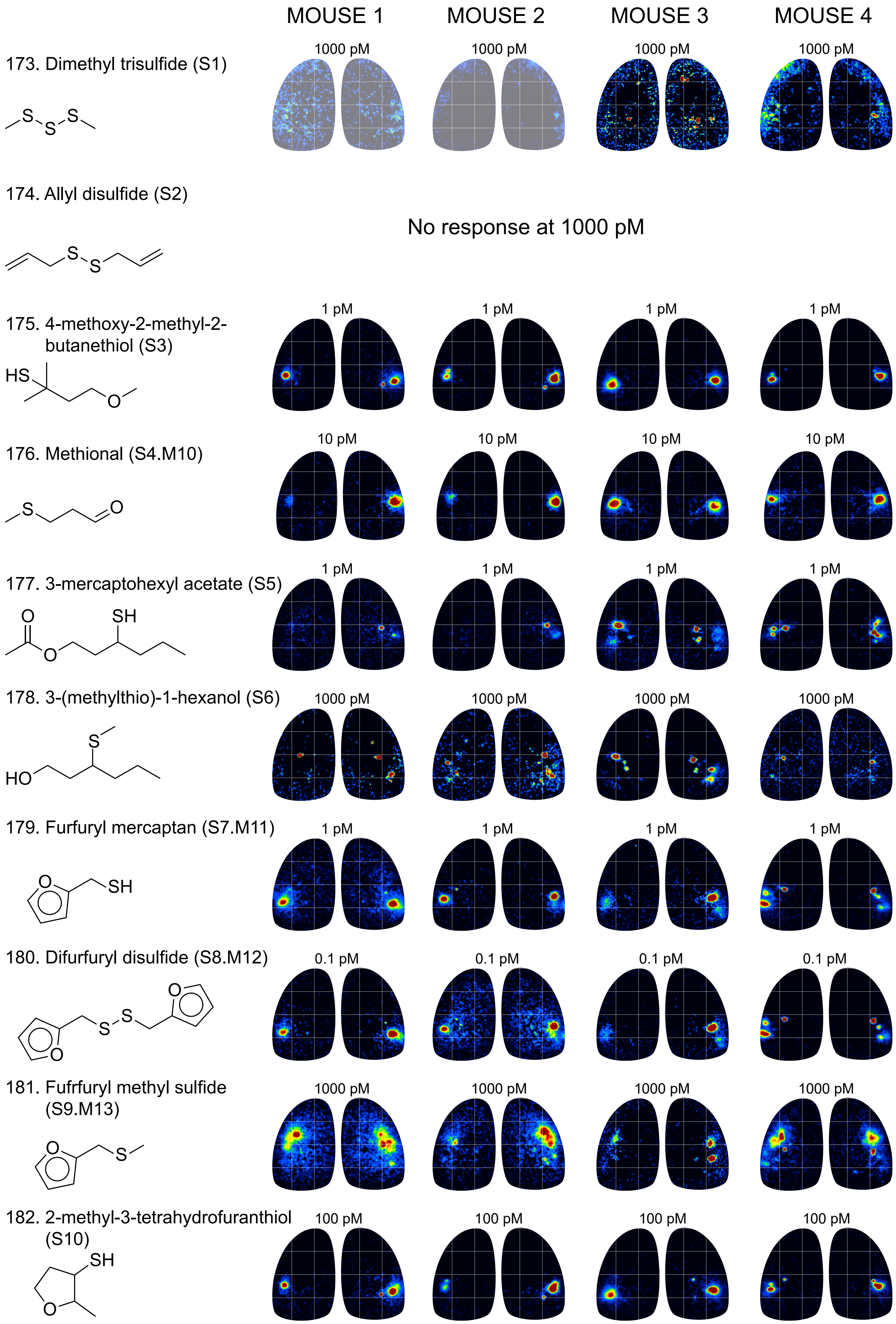

### Alkene

| | MOUSE 1 | MOUSE 2 | MOUSE 3 | MOUSE 4 |
|---|---|---|---|---|

183. Myrcene (ENE1)

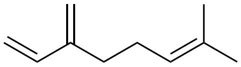

No response at 1000 pM

184. 1,3,5-undecatriene (ENE2)

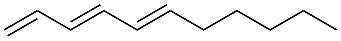

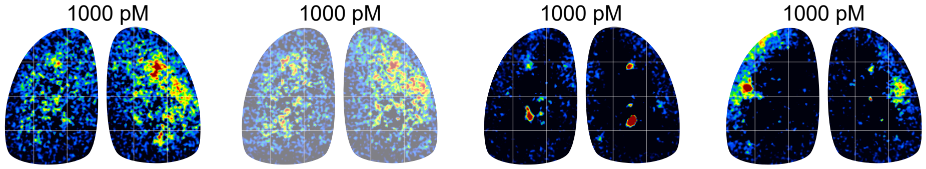


185. (R)-+-limonene (ENE3)

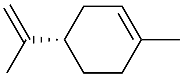

No response at 1000 pM
