## Supplemental Document S3, Atlas of Functionally Identified Glomeruli for "Mapping odorant sensitivities reveals a sparse but structured representation of olfactory chemical space by sensory input to the mouse olfactory bulb"

**Supplementary Material, Burton et al., 2022**

**Document S3. Atlas of Functionally Identified Glomeruli.**

Compilation of diagnostic odorants and example response maps for each of the 26 functionally-identified glomeruli (see Text and Table S2). Concentrations are given as estimated delivered concentration, rounded to the nearest order of magnitude, as in Document S2. See Table S2 for more precise estimates of delivered concentration. Bar plots indicate relative response magnitudes evoked by all effective odorants for each glomerulus, taken from the median response spectrum across the 8 OBs. Positions of each glomerulus are shown in lower right of p. 1, reproduced from Figure 2, main Text.

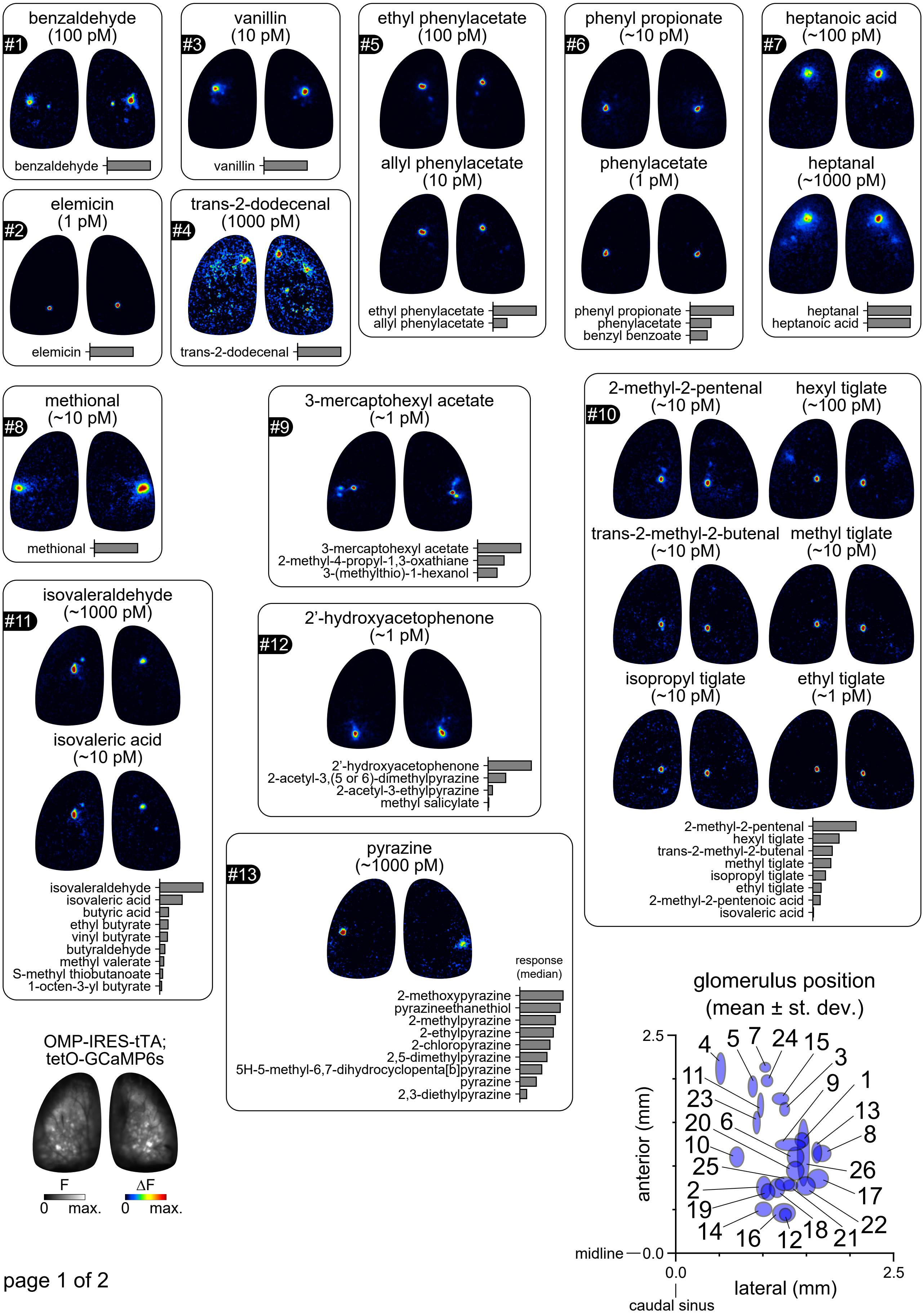

#1 benzaldehyde (100 pM)
benzaldehyde
#3 vanillin (10 pM)
vanillin
#5 ethyl phenylacetate (100 pM)
allyl phenylacetate (10 pM)
ethyl phenylacetate
allyl phenylacetate
#6 phenyl propionate (~10 pM)
phenylacetate (1 pM)
phenyl propionate
phenylacetate
benzyl benzoate
#7 heptanoic acid (~100 pM)
heptanal (~1000 pM)
heptanal
heptanoic acid
#2 elemicin (1 pM)
elemicin
#4 trans-2-dodecenal (1000 pM)
trans-2-dodecenal
#8 methional (~10 pM)
methional
#9 3-mercaptohexyl acetate (~1 pM)
3-mercaptohexyl acetate
2-methyl-4-propyl-1,3-oxathiane
3-(methylthio)-1-hexanol
#10 2-methyl-2-pentenal (~10 pM)
hexyl tiglate (~100 pM)
trans-2-methyl-2-butenal (~10 pM)
methyl tiglate (~10 pM)
isopropyl tiglate (~10 pM)
ethyl tiglate (~1 pM)
2-methyl-2-pentenal
hexyl tiglate
trans-2-methyl-2-butenal
methyl tiglate
isopropyl tiglate
ethyl tiglate
2-methyl-2-pentenoic acid
isovaleric acid
#11 isovaleraldehyde (~1000 pM)
isovaleric acid (~10 pM)
isovaleraldehyde
isovaleric acid
butyric acid
ethyl butyrate
vinyl butyrate
butyraldehyde
methyl valerate
S-methyl thiobutanoate
1-octen-3-yl butyrate
#12 2'-hydroxyacetophenone (~1 pM)
2'-hydroxyacetophenone
2-acetyl-3,(5 or 6)-dimethylpyrazine
2-acetyl-3-ethylpyrazine
methyl salicylate
#13 pyrazine (~1000 pM)
response (median)
2-methoxypyrazine
pyrazineethanethiol
2-methylpyrazine
2-ethylpyrazine
2-chloropyrazine
2,5-dimethylpyrazine
5H-5-methyl-6,7-dihydrocyclopenta[b]pyrazine
pyrazine
2,3-diethylpyrazine
OMP-IRES-tTA; tetO-GCaMP6s
F
0 max.
ΔF
0 max.
glomerulus position (mean ± st. dev.)
2.5
anterior (mm)
midline 0.0
0.0
2.5
lateral (mm)
caudal sinus
4 5 7 24 15 3
11 9 1
23 13
20 6 8
10
25 26
2 17
19 22
14 16 12 18 21


page 1 of 2

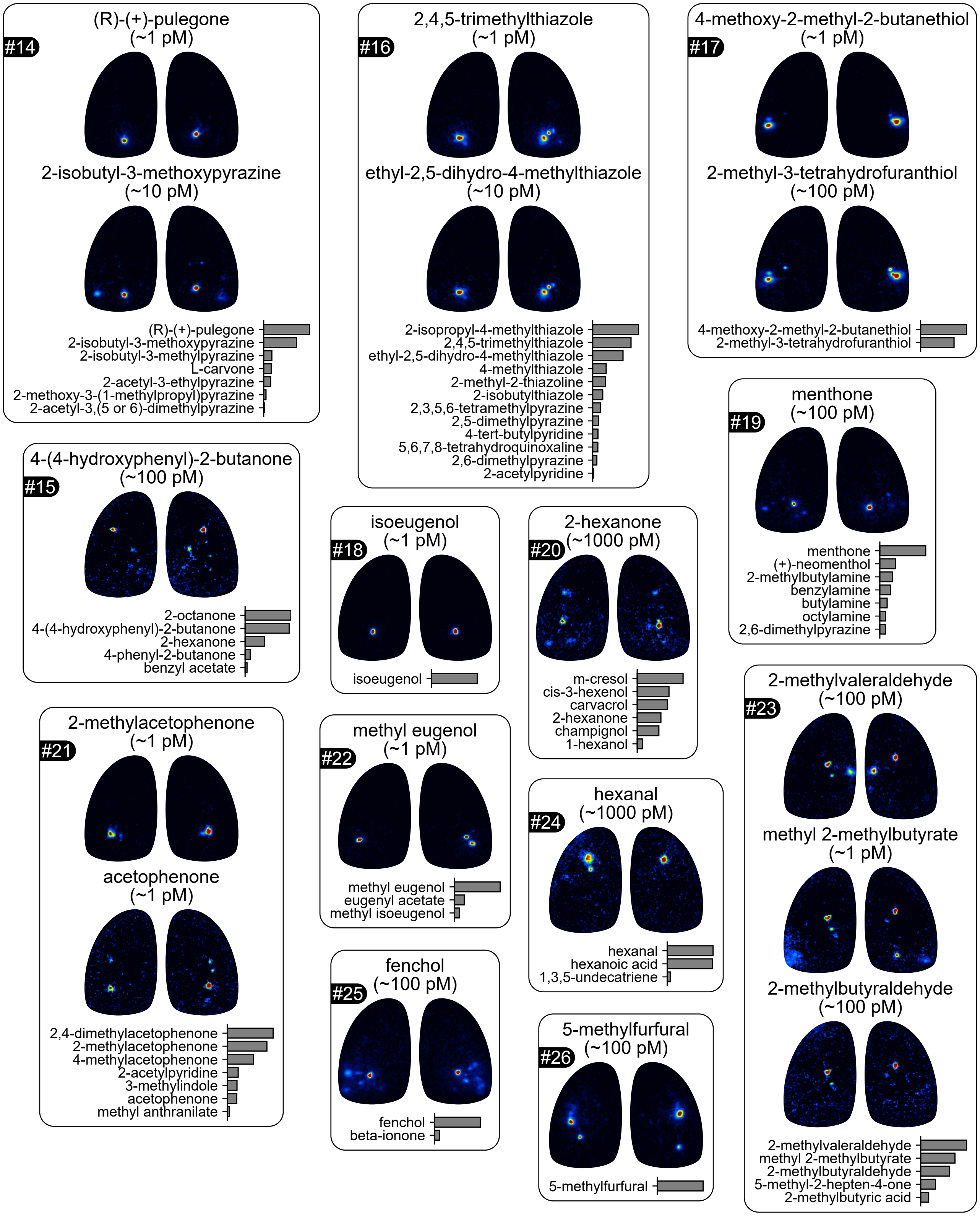
#14
(R)-(+)-pulegone
(~1 pM)
2-isobutyl-3-methoxypyrazine
(~10 pM)
(R)-(+)-pulegone
2-isobutyl-3-methoxypyrazine
2-isobutyl-3-methylpyrazine
L-carvone
2-acetyl-3-ethylpyrazine
2-methoxy-3-(1-methylpropyl)pyrazine
2-acetyl-3,(5 or 6)-dimethylpyrazine
#16
2,4,5-trimethylthiazole
(~1 pM)
ethyl-2,5-dihydro-4-methylthiazole
(~10 pM)
2-isopropyl-4-methylthiazole
2,4,5-trimethylthiazole
ethyl-2,5-dihydro-4-methylthiazole
4-methylthiazole
2-methyl-2-thiazoline
2-isobutylthiazole
2,3,5,6-tetramethylpyrazine
2,5-dimethylpyrazine
4-tert-butylpyridine
5,6,7,8-tetrahydroquinoxaline
2,6-dimethylpyrazine
2-acetylpyridine
#17
4-methoxy-2-methyl-2-butanethiol
(~1 pM)
2-methyl-3-tetrahydrofuranthiol
(~100 pM)
4-methoxy-2-methyl-2-butanethiol
2-methyl-3-tetrahydrofuranthiol
#19
menthone
(~100 pM)
menthone
(+)-neomenthol
2-methylbutylamine
benzylamine
butylamine
octylamine
2,6-dimethylpyrazine
#15
4-(4-hydroxyphenyl)-2-butanone
(~100 pM)
2-octanone
4-(4-hydroxyphenyl)-2-butanone
2-hexanone
4-phenyl-2-butanone
benzyl acetate
#18
isoeugenol
(~1 pM)
isoeugenol
#20
2-hexanone
(~1000 pM)
m-cresol
cis-3-hexenol
carvacrol
2-hexanone
champignol
1-hexanol
#23
2-methylvaleraldehyde
(~100 pM)
methyl 2-methylbutyrate
(~1 pM)
2-methylbutyraldehyde
(~100 pM)
2-methylvaleraldehyde
methyl 2-methylbutyrate
2-methylbutyraldehyde
5-methyl-2-hepten-4-one
2-methylbutyric acid
#21
2-methylacetophenone
(~1 pM)
acetophenone
(~1 pM)
2,4-dimethylacetophenone
2-methylacetophenone
4-methylacetophenone
2-acetylpyridine
3-methylindole
acetophenone
methyl anthranilate
#22
methyl eugenol
(~1 pM)
methyl eugenol
eugenyl acetate
methyl isoeugenol
#24
hexanal
(~1000 pM)
hexanal
hexanoic acid
1,3,5-undecatriene
#25
fenchol
(~100 pM)
fenchol
beta-ionone
#26
5-methylfurfural
(~100 pM)
5-methylfurfural
